# iAstrocytes model cytokine influences on complement expression and neuronal network synchronization

**DOI:** 10.64898/2026.06.04.730242

**Authors:** Nader Morshed, Matthew Demers, Ana Gonzalez-Ramos, Henna Jäntti, Jordan Doman, Sean D’Souza, Letian Li, Adam J. Granger, Matthew B. Johnson, Beth Stevens

## Abstract

Astrocytes play essential roles in neuronal development, function, and disease, yet existing methods to derive astrocytes from human pluripotent stem cells (hPSCs) are complex and can involve months of *in vitro* maturation. We developed a genomic safe-harbor knock-in system for inducible expression of the astrogenic transcription factors NFIA, NFIB, and SOX9, enabling rapid and robust generation of functional induced astrocytes (iAstrocytes). Across five hPSC lines, NFIB-SOX9 and NFIA-NFIB-SOX9 combinations efficiently generated highly pure populations expressing astrocyte-specific and synaptogenic genes. iAstrocytes displayed cytokine-induced expression of complement factors C3 and C4 and were amenable to CRISPR interference (CRISPRi) gene expression knockdown. Optimization of culture conditions enabled survival of NFIB-SOX9 iAstrocytes in co-culture with human induced neurons (iNeurons). Through pharmacological and genetic perturbations, we uncovered a previously undescribed phenomenon in which co-culture with iAstrocytes promoted the development of synchronized iNeuron network calcium activity mediated by specific gap junction proteins. This rapid and genetically tractable iAstrocyte platform provides a robust model to dissect human genetic and environmental effects on astrocyte-neuron interactions.

## Introduction

Astrocytes are essential regulators of neuronal development and function, supporting synaptogenesis, glutamate homeostasis, and lipid metabolism^1,2^. Dysregulated astrocyte signaling is increasingly implicated in brain dysfunction across the lifespan and in disease states^3–9^. In these contexts, astrocytes exhibit heightened cytokine and oxidative stress responses^4,10–13^ and increased expression of complement components^9,12,14^, a pathway genetically and functionally linked to neuropsychiatric and neurodegenerative disease risk and pathologies^15–17^. However, the upstream cellular mechanisms controlling astrocytes’ regulation of synapse formation, neuronal function, and disease-associated states remain poorly understood.

Although co-cultures of human induced neurons (iNeurons) with mouse astrocytes have revealed insights into neuronal and synaptic development^18–20^, species differences in astrocyte biology^21–23^ and the astrocytic expression of human-specific disease risk proteoforms such as APOE and complement C4 underscore the need for fully human systems. Human pluripotent stem cell (hPSC)-derived astrocytes provide a tractable model to investigate how environmental factors and human genetics shape astrocyte function and neuronal support^24,25^. Earlier hPSC-to-astrocyte differentiation protocols rely on morphogen patterning^26–28^ or proprietary media^29,30^, and typically require 1-6 months to yield functional astrocytes. Recent studies have shown that induced expression of NFIA, NFIB, and SOX9 transcription factor (TF) combinations can accelerate astrocyte differentiation, generating functional cells within 2-4 weeks^31–38^. While these methods have demonstrated utility^39^, they have drawbacks limiting their use, including lentivirus transduction variability and biosafety concerns, and their applicability and reproducibility across diverse human genetic backgrounds has not been extensively tested.

Here we leveraged a genomic safe harbor knock-in system^40^ to induce expression of combinations of NFIA, NFIB, and SOX9. We generated iAstrocytes using NFIA-NFIB-SOX9 (“AB9”) and NFIB-SOX9 (“NBS9”) across both human embryonic stem cell (hESC) and induced pluripotent stem cell (iPSC) lines, including three male and two female lines. We comprehensively characterized iAstrocytes using flow cytometry, immunofluorescence (IF), transcriptomic profiling, and assays for synaptogenic factor secretion. To enable genetic perturbations, we established engineered hPSC lines for CRISPR interference (CRISPRi) knockdown (KD)^41^ in either iAstrocytes or iNeurons and demonstrated efficient KD in iAstrocytes in monoculture as well as in iNeurons in co-culture with iAstrocytes. We further examined cytokine-induced states in iAstrocytes, focusing on complement factors C4 and C3 which are upregulated by astrocytes in neurodegeneration and aging^9,12,14^ and are involved in complement-mediated synapse engulfment^42^. We demonstrated the capacity of iAstrocytes to upregulate C4 and C3, established specificity of stimulus-response relationships and used CRISPRi to efficiently suppress induction of complement factors and other disease-associated responses.

To examine the ability of iAstrocytes to promote iNeuron maturation and synaptic connectivity, we used fluorescent labels and calcium sensors to identify optimal conditions for co-culture attachment, survival, and neuronal calcium imaging. These experiments revealed synchronized neuronal calcium activity that emerged as early as 9-14 days in co-culture (12-17 days post-differentiation of iNeurons). As proof of principle of the use of this platform for functional studies we performed pharmacological and genetic perturbations to identify the molecular mechanisms mediating this synchronous neuronal activity. We found that pharmacological inhibition of gap junctions or neuron-specific CRISPRi KD of *GJC1* (connexin 45) disrupted synchronous neuronal calcium activity. Together, these findings establish a rapid, fully human, and perturbable model for investigating astrocyte-neuron interactions and provides a robust system to study genetic and environmental influences.

## Results

### Safe harbor astrogenic transcription factor expression generates fast, reproducible iAstrocytes across hPSC lines

We first optimized a one-step genomic safe harbor knock-in strategy for iAstrocyte induction, adapting a vector previously used to generate NGN2-iNeurons^30,40,43^. Electroporation or lipofection delivery of Cas9 RNP and homology dependent repair-mediated plasmid integration targeting AAVS1 or CLYBL loci achieved stable reporter expression in 5-10% of H1 hESCs (**Fig S1A**), which increased to >80% following primary antibiotic selection with geneticin (**Fig S1B**). Expression efficiency increased to >99% of cells after secondary doxycycline-dependent antibiotic selection using zeocin (**Fig S1B**). Reporter expression was induced after 48 hours to a similar degree across doxycycline concentrations ranging from 500-2000 ng/mL (**Fig S1C**). Dual-resistance selection (e.g., NeoR-ZeoR, HygroR-PuroR, BlastR-PuroR) enabled flexible orthogonal selection schemes in H1 hESCs (**Fig S1D-E**). This dual selection approach enabled stable and efficient inducible expression of transgenes across hPSC lines without clonal isolation.

We next introduced inducible NFIB only, NBS9, or AB9 constructs into AAVS1 or CLYBL loci of H1 hESCs (**Fig 1A, S2A**). Within 7 days of induction in media containing serum, morphogens, and growth factors (**Fig 1B**), H1 AAVS1-NBS9 iAstrocytes expressed canonical astrocyte markers (CD44, CD49f, EGFR, S100B, VIM, ALDH1L1/2) (**Fig 1C-D**). These cells lacked markers for myeloid (CD45, P2RY12, CX3CR1, CD43, CD41a, CD235a), neuronal (CXCR4, CD325), and oligodendrocyte precursor cells (O4, PDGFRa, NG2), confirming lineage specificity. By day 21, iAstrocytes also expressed osteonectin (SPARC) and thrombospondin 1 (THBS1) (**Fig 1E**), and IF analysis revealed widespread VIM, S100B, CD44, SOX9, SPARC, and APOE expression as well as sporadic GFAP expression (**Fig 1F**). We confirmed that most iAstrocytes retained AAVS1 safe harbor-driven expression through day 21, as indicated by detection of the V5 epitope tag specific to exogenous SOX9 (**Fig 1F**). iAstrocytes expressing NFIB, NBS9, or AB9 from the CLYBL locus all expressed CD44, EGFR, and other surface markers by day 7 indicating acquisition of astrocyte identity (**Fig S2B**). CLYBL-NBS9 iAstrocytes proliferated more rapidly in the period between days 7 and 15 compared to those produced using CLYBL-AB9 (**Fig S2C**).

**Figure 1:**
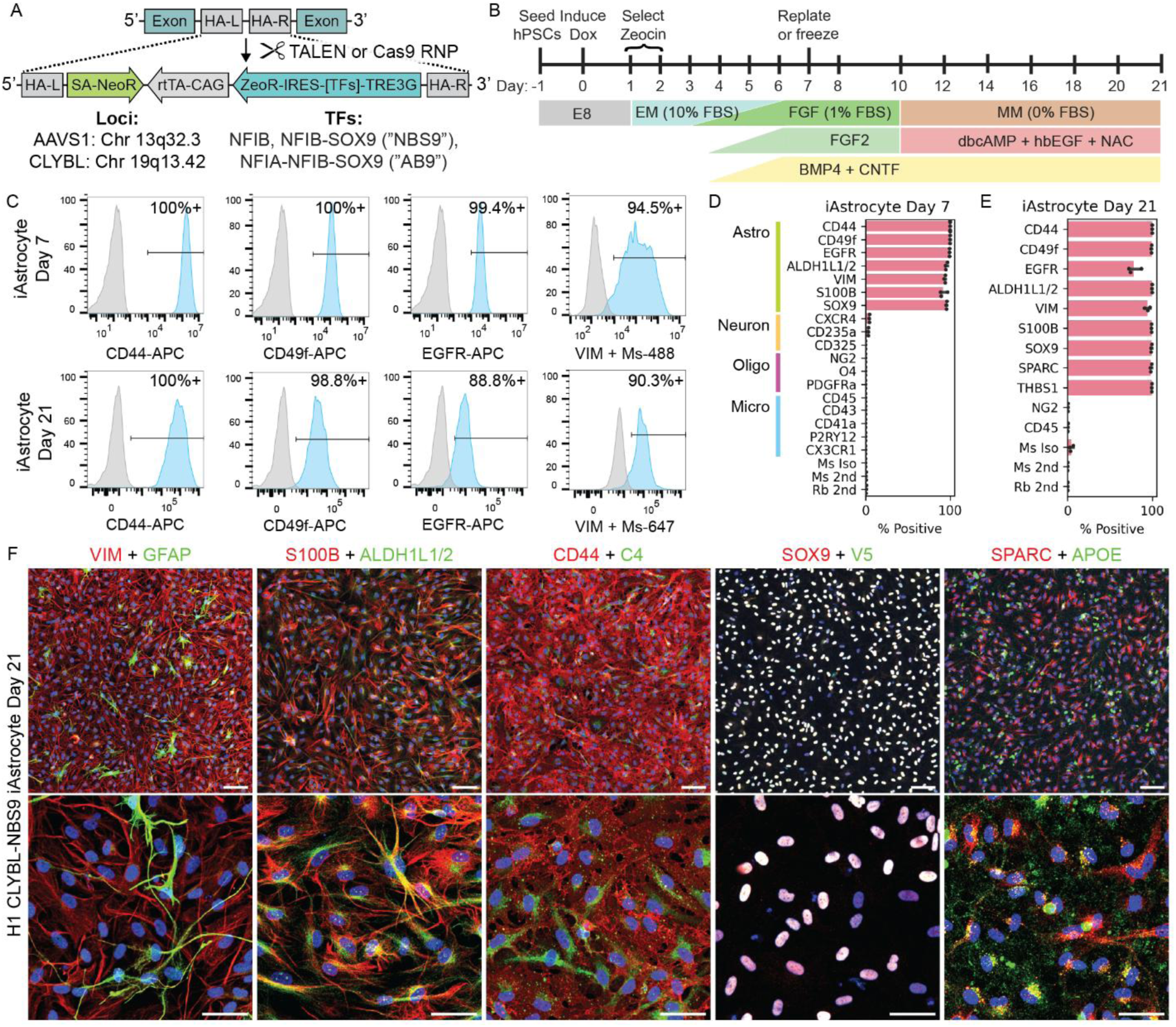
Safe harbor knock-in of astrogenic transcription factors enables rapid astrocyte differentiation. (A) Schematic for knock-in of astrogenic transcription factors to AAVS1 or CLYBL genomic safe harbor loci using TALENs or Cas9. HA, homology arm; SA, splice acceptor; rtTA, reverse tetracycline-controlled transactivator; TRE3G: Tet-On 3rd generation; IRES: internal ribosomal entry site; NeoR: neomycin/geneticin resistance; ZeoR: zeocin resistance. (B) Timeline of astrocyte differentiation protocol. EM: expansion medium; FGF: medium containing fibroblast growth factor; DM: differentiation medium. (C) Flow cytometry quantification of CD44, CD49f, EGFR, and VIM in H1 AAVS1-NBS9 iAstrocytes on day 7 (top row) and day 21 (bottom row). Blue = Stained population, Grey = No primary control. (D-E) Percentage of H1 AAVS1-NBS9 iAstrocytes positive for surface and internal markers at (D) day 7 and (E) day 21, measured by flow cytometry. (n=3 replicates per timepoint from 1 differentiation) (F) IF images of iAstrocyte markers in H1 CLYBL-NBS9 iAstrocytes on day 21 at 10x and 20x magnification: GFAP and VIM, S100B and ALDH1L1/2, CD44 and C4, total SOX9 and inducible V5-tagged SOX9, and SPARC and APOE. Green = GFAP, ALDH1L1/2, C4, SOX9, or APOE, Red = VIM, S100B, CD44, V5, or SPARC, Blue = Nuclei. Scale = 100 μm (top row) or 50 μm (bottom row).

We assessed the reproducibility of iAstrocyte induction across three male and two female hESC and iPSC lines (H1, H9, KOLF2.1J, FA13.1J, WTC11). We generated hPSCs with NFIB, NBS9, or AB9 in the CLYBL locus and obtained 11 total engineered lines that survived both chemical selection steps (see: **Key Resources Table**). Across all engineered lines, differentiation yields on day 7 varied between 5-50 iAstrocytes per seeded hPSC (**Fig S2D**). In nine engineered hPSC lines that were assessed across at least three independent differentiation batches, iAstrocytes consistently upregulated CD44 and EGFR and downregulated TRA-1-60R and TRA-1-81 pluripotency markers by day 7 (**Fig S2E**). Inducible SOX9 with V5-epitope tag was expressed and P2A linker was efficiently cleaved across all NBS9 and AB9 lines and not present in lines expressing only NFIB (**Fig S2F**).

### iAstrocytes resemble synapse-supporting human astrocytes and are amenable to CRISPRi gene perturbation

To assess cell identity and compare gene expression profiles with other models, we performed transcriptomic profiling of 7 NBS9 and AB9 iAstrocyte lines from 4 genetic backgrounds at day 21 and 1 uninduced line (stem cell state). One differentiation (H1 CLYBL-NBS9) exhibited an interferon response associated with overconfluency, which could be prevented by a lower day 7 seeding density (**Fig S2G**), and was excluded from downstream analyses. Principal component analysis showed a uniform transcriptional shift from pluripotency to astrocytic identity along PC1, accounting for nearly half of transcriptional variance, while inter-line variability along PC2 only accounted for 10% of variance (**Fig 2A, Table S1**). Comparing hPSCs to iAstrocytes on the same genetic background revealed 2,625 differentially expressed genes (DEGs) (**Fig 2B, Table S2**), whereas CLYBL-NBS9 iAstrocytes from two different genetic backgrounds differed by only 103 DEGs (**Fig S3A, Table S2**), and NBS9 versus AB9 iAstrocytes, pooled across two backgrounds yielded only 68 DEGs, including *NFIA* itself (**Fig S3B, Table S2**). Although NBS9 did not induce significant NFIA expression, both TF combinations produce highly transcriptionally similar astrocytes.

**Figure 2.**
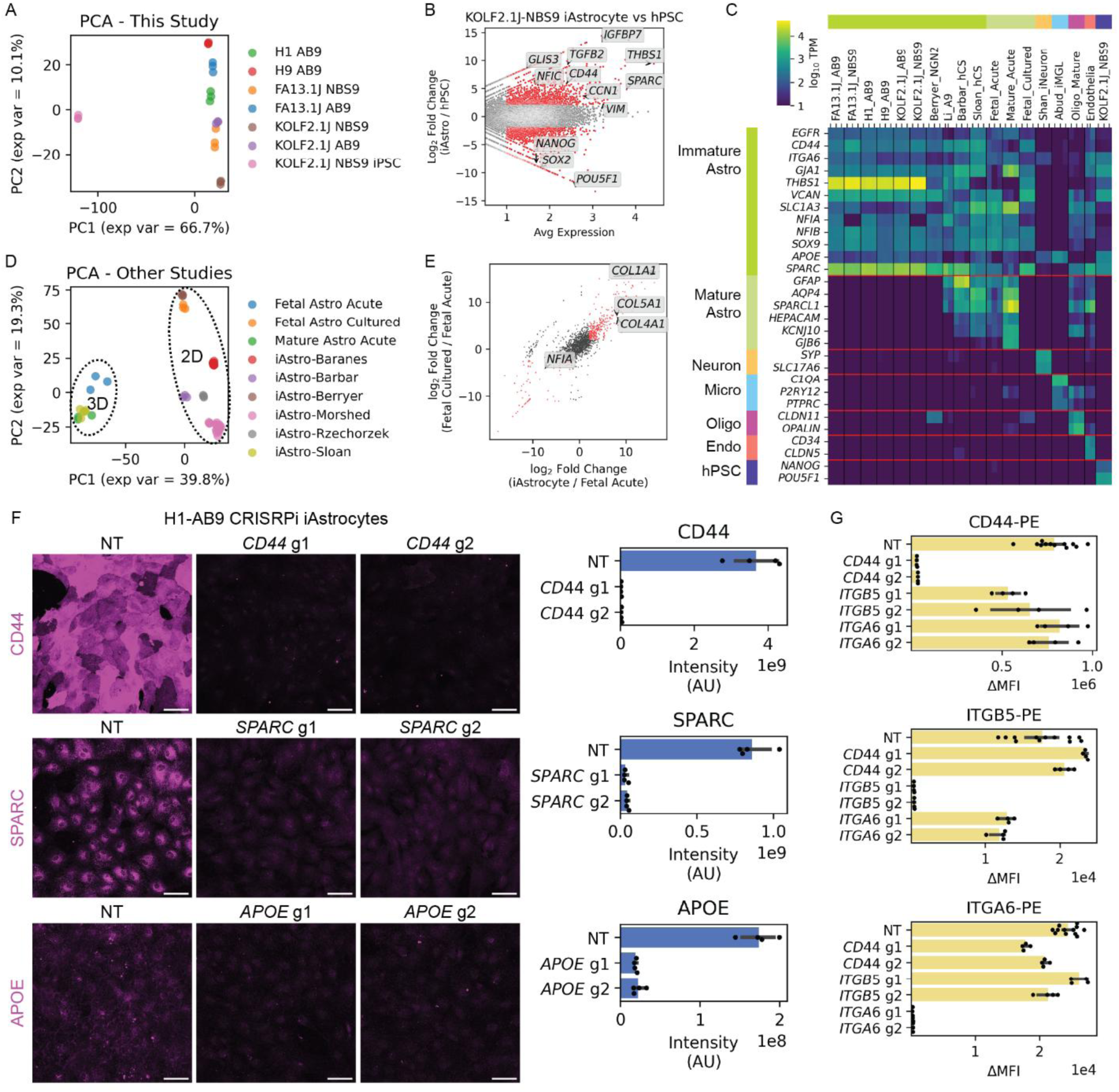
RNA gene expression and CRISPRi gene knockdown in iAstrocytes. (A) Principal component analysis (PCA) comparing KOLF2.1J iPSCs with iAstrocytes induced with CLYBL-NBS9 or AB9 from multiple hPSC backgrounds. (B) Volcano plot for RNA markers enriched in KOLF2.1J-NBS9 iAstrocytes relative to uninduced hPSCs. (C) Bulk RNA-seq quantification of immature astrocyte, mature astrocyte, neuron, microglia (Micro), endothelia (Endo), oligodendrocyte (Oligo), and hPSC marker genes in iAstrocytes from this study and other published studies of iAstrocytes, iNeurons, iMGLs, primary astrocytes, primary oligodendrocytes, and primary endothelia. Datasets are normalized to transcripts per million (TPM). (D) PCA comparing this study’s iAstrocytes with previously published iAstrocyte models, acutely isolated astrocytes, and primary cultured human astrocytes. (E) Fold change in gene expression between acutely isolated fetal astrocytes and either cultured fetal astrocytes or KOLF2.1J-NBS9 iAstrocytes. (F) (left) IF images of CD44, SPARC, and APOE in H1-AB9 CRISPRi iAstrocytes on day 21 after LV transduction on day 9. Magenta = Secondary-647 + iRFP670. Scale = 100 μm; (right) Quantification of IF intensities of CD44, SPARC, and APOE KD. (N=4 replicates per condition from 1 differentiation) (G) Flow cytometry quantification of CD44, ITGA6, and ITGB5 on H1-AB9 CRISPRi iAstrocytes on day 21 after LV transduction on day 9. (n=4 replicates per condition from 1 differentiation)

iAstrocytes consistently upregulated astrocyte genes including *CD44*, *EGFR*, and *VIM* (**Table S2**), and showed greater expression of *THBS1*, *CCN1*, *CCN2*, *TGFB2*, *SPARC*, *NFIC*, and *NFIX* relative to hPSCs, iNeurons^44^, and iMicroglia^45^ (**Fig S3C**, **Table S2**). Marker genes for neurons (*SYP, SLC17A6*), microglia (*C1QA*, *P2RY12*), oligodendrocytes (*CLDN11, OPALIN*), and endothelia (*CD34*, *CLDN5*) were absent in iAstrocytes (**Fig 2C**). Among genes involved in synaptic development, all iAstrocyte lines expressed glypicans (*GPC1*, *GPC2*, *GPC4*, and *GPC6*), thrombospondins (*THBS1*, *THBS2*, *THBS3*), *SPARC*, and cellular communication network factors (*CCN1*, *CCN2*, *CCN3*) (**Fig S3D**). Together, these data indicate that iAstrocytes express astrocyte-specific genes and genes supporting neuronal synaptic development.

Integration with bulk RNA-seq from multiple iAstrocyte models^27,30,33,46,47^, acutely isolated human fetal and adult astrocytes^48^, and primary cultured human fetal astrocytes^30^ showed that our iAstrocytes were most similar to other TF-based iAstrocyte models^30,33,47^ and to CD49f^+^ iAstrocytes isolated from 3D neural spheres and further differentiated in 2D^27^ (**Fig 2C-D, Table S2**). In contrast, iAstrocytes from long-term 3D cortical organoid suspension cultures^46^ expressed higher levels of late maturity markers (*SPARCL1*, *AQP4*, *GFAP*, *HEPACAM*, *KCNJ10*; **Fig 2C**) and clustered more closely with acutely profiled primary fetal and adult astrocytes^48^ (**Fig 2D**). Notably, primary fetal astrocytes cultured in 2D prior to sequencing^30^ were more similar to 2D cultured iAstrocyte models than to either acutely profiled primary astrocytes^48^ or iAstrocytes from long-term 3D organoid cultures^46^, suggesting that 2D versus 3D culture conditions drive a significant component of *in vitro* expression signatures across models. NBS9 iAstrocytes and cultured fetal astrocytes expressed higher levels of collagen factors (*COL1A1*, *COL4A1*, *COL5A1*) and lower *NFIA* compared to acutely isolated fetal astrocytes (**Fig 2E**, **Table S2**).

We next established a platform to study iAstrocyte functions using CRISPRi perturbation. We generated CRISPRi-compatible hPSC lines co-expressing ZIM3-KRAB-dCas9 plus iRFP670 or a HaloTag from piggyBac vectors^49^. We isolated H1 clones of CLYBL-AB9 with CRISPRi-iRFP and CLYBL-NBS9 with CRISPRi-Halo (see: **Methods**). Flow cytometry confirmed unimodal iRFP670 or HaloTag signals across all clones (**Fig S4A-B**). To validate CRISPRi activity in iAstrocytes, we selected astrocytic surface glycoprotein CD44, integrin α6 (*ITGA6*, encoding CD49f), and integrin β5 (*ITGB5*)^27,50^. We first targeted the surface integrin ITGB5 in uninduced hPSCs using lentiviral (LV) delivery of a single guide RNA (sgRNA) and co-expressing mNeonGreen^51^. We observed efficient KD of ITGB5 protein in transduced hPSCs (**Fig S4C**). We next tested KD in differentiated iAstrocytes, using lentivirus (LV) to transduce cells with sgRNA at day 9 post differentiation. We targeted CD44, ITGA6 and ITGB5 as well as SPARC and APOE, and observed efficient KD using imaging analysis (**Fig 2F**) or surface flow cytometry (**Fig 2G**). We observed similar KD efficiency for CD44 in both CRISPRi-iRFP and CRISPRi-Halo iAstrocytes (**Fig S4C**) and validated downregulation of additional targets *GPC4*, *COL1A1*, *S100B*, *VIM*, and *SLC1A3* by qPCR (**Fig S4D**). Together, these results demonstrate efficient, specific, and reproducible CRISPRi-mediated KD of target genes in iAstrocytes, establishing a robust platform for genetic perturbation.

### Specificity of cytokine-induced complement and immune responses in iAstrocytes

We tested the ability of our iAstrocyte model to recapitulate established reactive astrocyte responses involving the complement pathway^10,12,24^, which has been linked to neurodegeneration and synapse loss in conditions including Alzheimer’s disease^52–54^, Huntington’s disease^55^, and others^15,42,56–58^. Although the context-specific cellular mechanisms of complement expression and regulation in the brain are incompletely understood, prior studies have shown that astrocytes upregulate complement component C3 in response to co-treatment with TNFα, IL1α, and C1q (TIC)^10,59,60^ and complement component C4 in response to IFNγ^24,61^. Reactive iAstrocytes can also upregulate the surface proteins MHC class I (MHC-I)^62^ and Intercellular Adhesion Molecule 1 (ICAM1)^27,59,63^ which are both elevated in Alzheimer’s disease brain tissues^64–69^. However, it remains unclear whether other interferons also induce these responses. To explore this, we treated iAstrocytes for 24 hours with TIC, IFNγ, IFNβ, or IFNα variants (IFNαU, IFNα2b, IFNα6) in a serum-free medium and performed IF analysis for C4, C3, CD44, and ICAM1. TIC robustly induced C3 and ICAM1 protein expression in iAstrocytes, with IFNγ also eliciting a small ICAM1 response (**Fig 3A-B, S5A-B**). In contrast, IFNα variants, IFNβ, and IFNγ all induced C4 protein expression (**Fig 3A-B, S5C**). We performed CRISPRi KD with two independent sgRNAs per target to validate our assays for ICAM1, C3, and C4 and observed robust repression of cytokine-induced responses (**Fig S5A-D**). As the C4 sgRNAs used here target the promoters of both *C4A* and *C4B* isoforms, we overexpressed C4A and C4B individually in HEK293T cells and found that the C4 antibody recognizes both isoforms (**Fig S5E**), suggesting that cytokine induction and knockdown results reflect the cumulative expression levels of both isoforms in these experiments.

**Figure 3:**
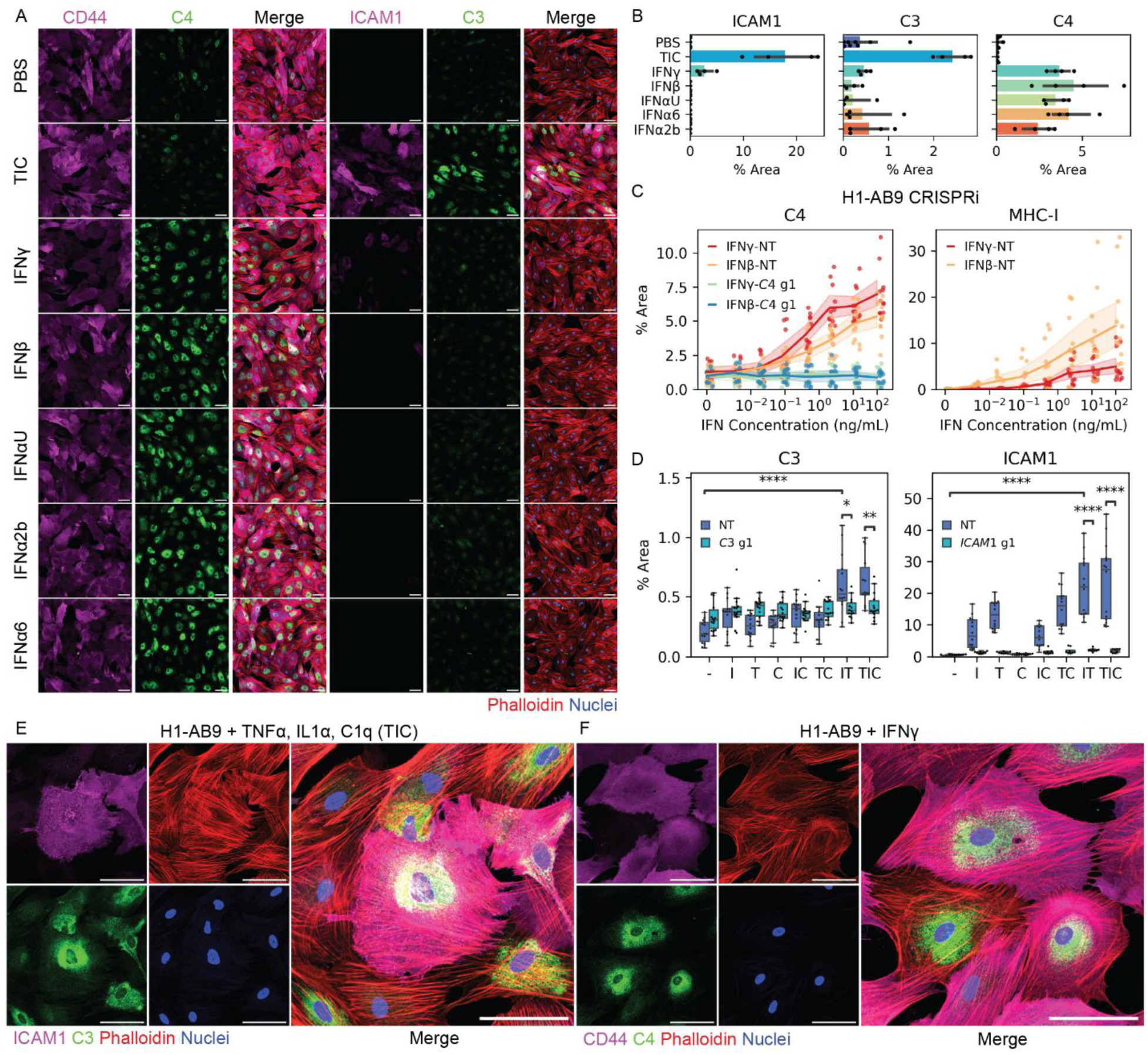
TIC and Interferon treatments induce iAstrocyte expression of reactivity markers. (A) IF images of day 21 H1-AB9 iAstrocytes after 24-hour treatment with TIC (TNFα, IL-1α, and C1q), IFNγ, IFNβ, IFNαU, IFNα2b, or IFNα6. CD44, C4, merge (left) and ICAM1, C3, merge (right). Red = Phalloidin, Green = C3 or C4, Magenta = ICAM1 or CD44. Scale = 100 μm. (B) Quantification of ICAM1, C3, and C4 protein expression across replicates of conditions shown in (A). *p < 0.05, **p < 0.01, ***p < 1e-3, ****p < 1e-4, unpaired Student’s t-test, two-sided. (n=4-8 replicates per condition from 1 differentiation) (C) Quantification of C4 and MHC-I expression in H1-AB9 CRISPRi iAstrocytes transduced with LV targeting *C4* or NT and treated with descending doses of IFNβ or IFNγ for 24 hours. *p < 0.05, **p < 0.01, ***p < 1e-3, ****p < 1e-4, unpaired Student’s t-test, two-sided. (n=48 replicates per condition from 4 differentiations) (D) Quantification of C3 and ICAM1 expression in H1-AB9 CRISPRi iAstrocytes transduced with LV targeting *C3*, *ICAM1,* or NT and treated with combinations of IL1α “I”, TNFα “T”, and C1q “C” for 24 hours. *p < 0.05, **p < 0.01, ***p < 1e-3, ****p < 1e-4, unpaired Student’s t-test, two-sided. (n=32-56 replicates per condition from 3 differentiations) (E-F) High-magnification IF images of H1-AB9 iAstrocytes stimulated for 24 hours with (E) TIC and stained for C3 and ICAM1 expression or (F) IFNγ and stained for C4 and CD44. Red = Phalloidin, Green = C3 or C4, Magenta = CD44. Scale = 100 μm.

Titration experiments revealed that iAstrocytes upregulated C4 and MHC-I proteins after 24 hour treatment with IFNβ or IFNγ at concentrations of 10-50 ng/mL (**Fig 3C, S5F**). C4 protein expression was suppressed by CRISPRi KD across all tested IFN concentrations (**Fig 3C**). Dissection of the TIC response revealed that IL-1α and TNFα were each individually sufficient to induce ICAM1 expression, with combined treatment further enhancing ICAM1 response and inducing C3 expression above background (**Fig 3D**). C1q had no detectable effect on ICAM1 or C3 expression either alone or in combination with IL-1α and/or TNFα (**Fig 3D**). CRISPRi KD of either target efficiently suppressed C3 or ICAM1 induction (**Fig 3D**). Together, these results establish that our iAstrocyte model can be used to investigate the molecular pathways linking disease-relevant stimuli to complement system activation and other aspects of reactive astrocyte states, and reveal novel insights into the specificity of astrocyte responses to individual cytokines.

### iAstrocytes secrete key factors supporting neuron development and synaptogenesis

We next used antibody staining and imaging to examine the ability of iAstrocytes to secrete protein factors reported to regulate neuronal development and synaptogenesis, key functions of astrocytes in the developing brain^1,70^. Our transcriptomic analysis revealed robust expression *THBS1* (**Fig 2C, S3C-D)** encoding a secreted protein that supports neuronal synaptic development. We performed immunofluorescence (IF) imaging of THBS1 and observed an extracellular basement membrane structure beneath iAstrocytes as well as small intracellular pockets (**Fig 4A-B**). This IF signal did not appear in vessels coated with matrigel but lacking iAstrocytes (**Fig S6A**). We tested the effects of vessel coating substrate and observed THBS1 protein deposition on vessels coated with matrigel or laminin substrates, but not on vessels with polymer substrates (PDL, PLL, PLO, dPGA) alone (**Fig 4C, S6A-B**). THBS1 was also deposited by iAstrocytes cultured on matrigel-coated plastic or glass coverslips (**Fig S6C**). To further validate the source of this signal, we performed CRISPRi KD of *THBS1* in iAstrocytes. We observed a decrease in THBS1 protein by IF in iAstrocytes transduced with sgRNA against the primary transcript start site (TSS) of *THBS1*, but not for an alternative TSS or in NT controls (**Fig 4D**).

**Figure 4.**
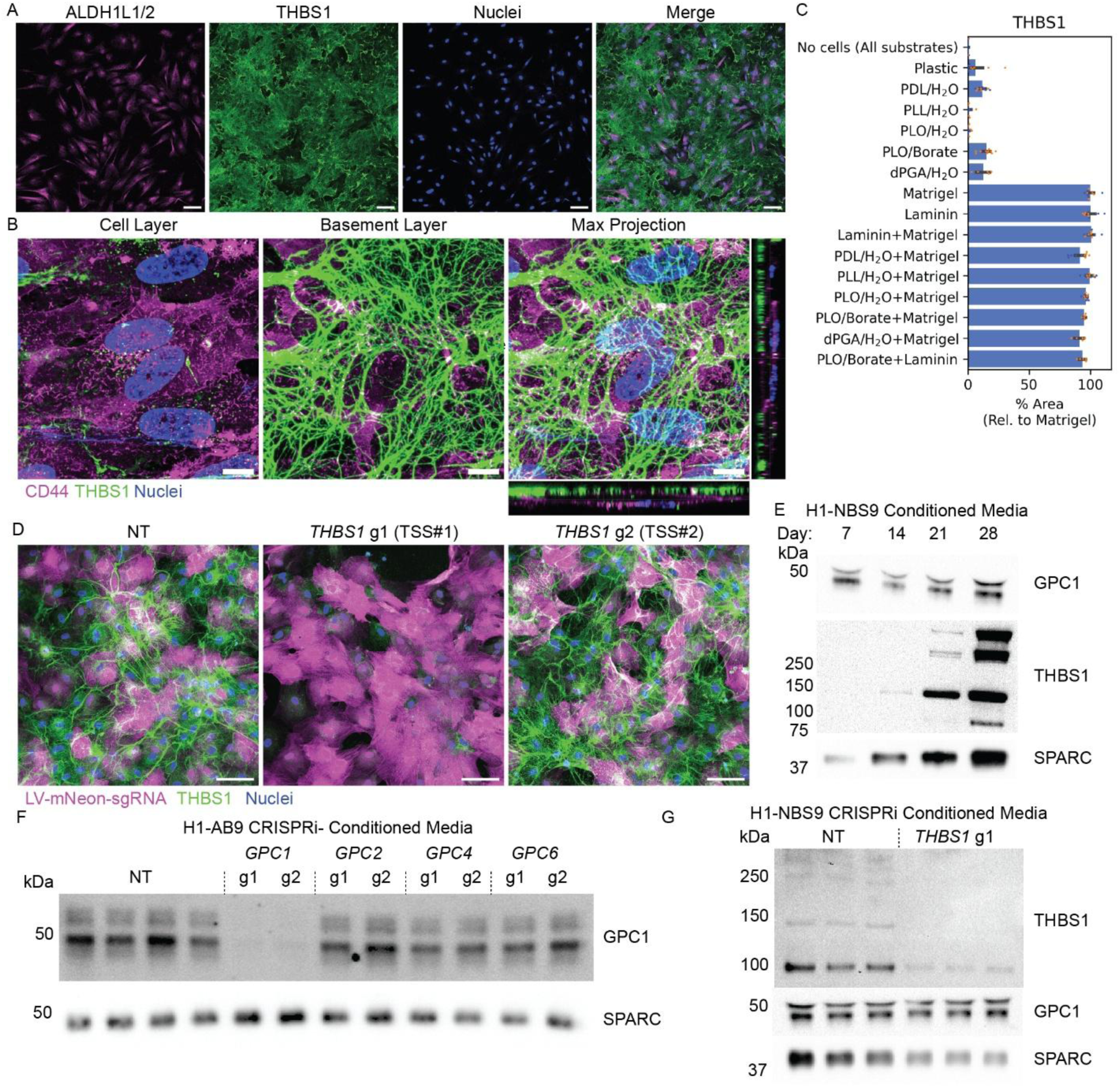
iAstrocytes secrete glypicans, thrombospondins, and other synaptogenic factors. (A) IF images of THBS1 and ALDH1L1/2 after culturing H1-NBS9 iAstrocytes to day 21. Green = ALDH1L1/2, Magenta = CD44, Blue = Nuclei. Scale = 100 μm. (B) High-magnification IF z-stack images of THBS1 and CD44 on H1-NBS9 iAstrocytes showing cell layer (left) basement layer (middle), and maximum z-stack projection with orthogonal views (right). Green = THBS1, Magenta = CD44, Blue = Nuclei. Scale = 10 μm. (C) IF quantification of THBS1 area on tissue culture-treated plastic vessels coated with mouse laminin, poly-D-lysine (PDL), poly-L-lysine (PLL), or poly-L-ornithine (PLO), or dendritic polyglycerol amine (dPGA), mouse laminin, with or without matrigel and H1-NBS9 iAstrocytes on day 21. (n=5-47 replicates per condition from 2 differentiations; replicates are colored by batch) (D) IF images of THBS1 expression in H1-AB9 CRISPRi iAstrocytes on day 21 after transduction with non-targeting or sgRNAs targeting *THBS1* transcription start sites (TSS) #1 or #2. Green = THBS1, Magenta = mNeonGreen, Blue = Nuclei. Scale = 100 μm. (E) Western blot detection of GPC1, THBS1, and SPARC in conditioned media from H1-NBS9 iAstrocytes between days 7 and 28. (F) Western blot detection of GPC1, and SPARC in conditioned media from H1-AB9 CRISPRi iAstrocytes on day 21 after CRISPRi LV knockdown of *GPC1*, *GPC2*, *GPC4*, or *GPC6* on day 9. (G) Western blot detection of THBS1, GPC1, and SPARC in conditioned media from H1-NBS9 CRISPRi iAstrocytes on day 21 after CRISPRi LV knockdown of *THBS1* on day 9.

Other proteins involved in brain development and expressed by iAstrocytes include APOE^1^, SPARC^1^, and glypican 1 (GPC1)^71^. We measured THBS1, GPC1, and SPARC proteins in conditioned media from iAstrocytes by western blot and observed the presence or increase of these factors between days 7 and 28 post induction (**Fig 4E**). We validated the specificity of our assays for GPC1, THBS1, and APOE in cell lysate or conditioned media using CRISPRi KD (**Fig 4F-G, S7A-B**). These experiments demonstrate that iAstrocytes secrete thrombospondins, glypicans, and other proteins involved in synaptic development and provide a method for functional manipulation of this process using CRISPRi.

### iAstrocytes promote synchronized neuronal network activity

We next asked whether iAstrocytes support the functional maturation of co-cultured iNeurons, focusing on cell attachment and morphology, survival, and calcium activity. iNeurons were generated from H1 hESCs by NGN2 expression and labeled using piggyBac-mediated expression of membrane-targeted fluorescent proteins to visualize cell morphology in live co-cultures (**Fig 5A**). iNeurons were differentiated to day 3 and iAstrocytes were differentiated to day 7 before both cell types were replated together. Co-culture days (ccD) refers to subsequent days post-replating. Co-culture of iNeurons with H1 CLYBL-NBS9 or -AB9 iAstrocytes on matrigel reduced neuronal clumping compared to monocultures (**Fig S8A**). Distribution of iNeurons was also more homogenous on substrates coated with poly-L-ornithine followed by matrigel (**Fig S8B**). Both iNeurons and iAstrocytes remained viable in Neurobasal with B27 and doxycycline (NBM-min) and did not require additional GlutaMAX, glucose, or non-essential amino acid supplements (**Fig S8C-D**). Neurobasal medium reformulated to maintain a target concentration of 5-10 mM glucose in co-cultures to promote mitochondrial oxidative phosphorylation^72^ (NBM-A g10) improved iAstrocyte survival relative to the original Neurobasal formulation (**Fig S8E**). Among the TF combinations tested, CLYBL-AB9 iAstrocytes showed poor survival after ccD 10, while CLYBL-NBS9 iAstrocytes had improved viability, maintaining 76-91% of their initial plating density by ccD 30 (**Fig S8E**).

**Figure 5.**
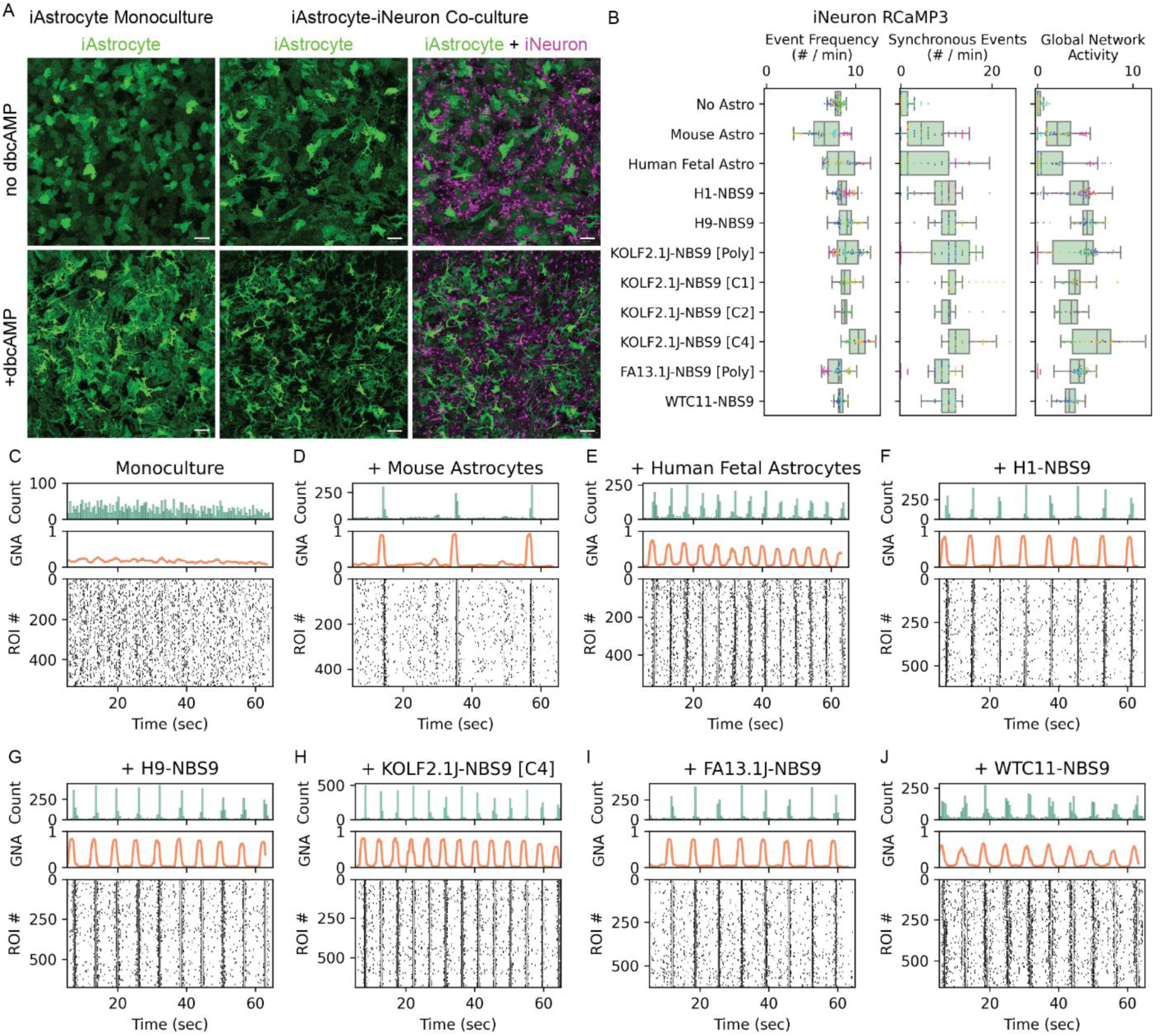
Co-culture with iAstrocytes enables synchronous neuronal calcium activity. (A) IF images of H1-NBS9 iAstrocytes in monoculture (left) or co-culture with iNeurons (bottom) without dbcAMP (top) or with dbcAMP (bottom) on ccD 12. Magenta = iNeurons, Green = iAstrocytes. Scale = 100 μm. (B) Quantification of calcium average single-neuron event frequency, synchronous event frequency, and GNA for RCaMP3 iNeurons in monoculture or co-culture with mouse primary, human primary, or human induced astrocytes on ccD 13-14. (N=20-83 replicates per condition from 2-6 differentiations per line; replicates are colored by batch) (C-J) Raster plots showing calcium event activity in RCaMP3 iNeurons in (C) monoculture, or co-culture with (D) mouse astrocytes, (E) human fetal astrocytes, (F) H1-NBS9, (G) H9-NBS9, (H) KOLF2.1J-NBS9 [C4], (I) FA13.1J-NBS9 [Poly], or (J) WTC11-NBS9 iAstrocytes on ccD 14.

To assess the impact of iAstrocytes on neuronal activity, we expressed GCaMP8f or RCaMP3 in iNeurons via piggyBac to monitor calcium activity in real time. We tested co-culture medium supplements used in prior studies reporting synchronous neuronal activity^18,44,73,74^ and found that addition of dibutyryl-cAMP (dbcAMP) and ascorbic acid increased GCaMP8f signal intensity compared to media containing only neurotrophic factors (NTFs: BDNF, GDNF, CNTF) (**Fig S9A**). iAstrocytes co-cultured with iNeurons in NBM-A g10 with dbcAMP displayed a more complex, process-bearing morphology, compared to iAstrocytes in monoculture without dbcAMP (**Fig 5A**), with both co-culture and dbcAMP contributing to this effect. To optimize calcium recording on ccD 14, we evaluated BrainPhys imaging (BPI) and NBM-A g10 as imaging media and compared continuous dbcAMP supplementation throughout the co-culture period or acutely prior to calcium imaging. Acute dbcAMP addition was necessary and sufficient to promote synchronized neuronal activity in BPI (**Video S1**). Synchronous activity was most frequent in NBM-A g10 with dbcAMP and was observed infrequently in NBM-A g10 lacking dbcAMP.

With this proof-of-concept functional co-culture assay, we next sought to establish an optimal imaging protocol and data analysis pipeline to reproducibly quantify iNeuron network calcium activity. We used a semi-automated pipeline to detect individual neuron calcium events, synchronous calcium events involving more than 30% of active neurons, and global network activity (GNA), a cumulative metric of network connectivity and activity defined as the sum of the fractions of neurons involved in all synchronous events over a set recording time (see **Methods**). We recorded calcium activity in NBM-min, NBM-A g10, or BPI and calculated average event frequency of individual neurons, frequency of synchronous events, and GNA (**Fig S9B, Video S2**). Co-culture with iAstrocytes in either NBM-min or NBM-A g10 supported the emergence of synchronized neuronal calcium activity, but synchronous events occurred less frequently and were more variable between wells after medium change to BPI (**Fig S9B-F**). Co-culture in NBM-A g10 promoted more robust network activity compared to NBM-min (**Fig S9B**). We examined network synchronous activity in a time course assay using NBM-A g10 and found that co-cultures with iAstrocytes develop synchronous activity between ccD 7 and 14, while co-cultures with mouse astrocytes developed synchronous activity between ccD 14 and 21 (**Fig S9G**). These results highlight an optimal medium in which iAstrocytes can reproducibly support iNeuron network formation and activity at an early co-culture stage.

To benchmark our system against primary astrocytes as well as across hPSC genetic backgrounds, we compared iNeuron monocultures with co-cultures containing primary mouse astrocytes, human fetal astrocytes, or iAstrocytes derived from five hPSC lines. On ccD 13-14, monocultures and co-cultures had similar single-neuron event frequencies, however co-cultures exhibited more frequent synchronous events and higher GNA compared to monocultures (**Fig 5B-J**). iAstrocytes derived with NBS9 from H1, H9, KOLF2.1J, FA13.1J, and WTC11 all supported network synchronization (**Fig 5F-J, Video S3**). Expression of GCaMP8f in iNeurons and RCaMP3 in iAstrocytes enabled dual-color imaging of calcium activity in both cell types (**Video S4**). Together, these findings demonstrate that iAstrocyte co-cultures drive the emergence of synchronized neuronal network activity, establishing a robust and perturbable platform for dissecting human astrocyte-neuron interactions.

### Neuronal gap junctions support synchronized calcium activity in iAstrocyte co-cultures

Previous studies demonstrated development of synchronous electrical and calcium activity of iNeurons co-cultured with mouse astrocytes or iAstrocytes mediated by AMPA and NMDA receptors^18,75–77^. During development, neurons can also propagate calcium signals through direct electrical coupling mediated by gap junctions formed by connexins^78–80^. To identify mechanisms contributing to synchronized activity in our co-cultures, we applied pharmacological perturbations at ccD 14. The AMPA receptor antagonist NBQX reduced synchronous event frequency and GNA without altering average single-neuron event frequency in co-cultures with primary mouse astrocytes (**Fig S9H-J**), but had no effect in co-cultures with iAstrocytes (**Fig 6A-C**). The NMDA receptor antagonist D-AP5 did not alter activity in either model (**Fig 6A, S9H, S10A-B**), consistent with previous reports that NMDAR-mediated currents appear later in co-culture with mouse astrocytes^18^. In contrast, in iAstrocyte co-cultures, the connexin inhibitors carbenoxolone^81^, meclofenamic acid^81^, mefloquine^81,82^, and TAT-Gap19^83^ reduced average single-neuron event frequency, synchronous event frequency, and GNA (**Fig 6A, 6D-E, S10C-D, Video S5**). Application of GABA suppressed all activity metrics, whereas the GABA_A_ receptor antagonist gabazine had no effect (**Fig 6A, S10E-F**), consistent with previous reports that NGN2 produces homogenous excitatory neurons that can be modulated by GABA but do not synthesize it^84^ and indicating that our iAstrocytes do not secrete GABA in this context either. The sodium channel blocker tetrodotoxin (TTX) abolished all coordinated activity, confirming these events represent action potential-dependent signaling (**Fig 6A, S10G**). Similarly, the intracellular calcium chelator BAPTA-AM blocked all coordinated activity (**Fig 6A, S10H**). Comparable pharmacological responses were observed in GCaMP8f iNeurons cultured in BPI medium (**Fig S10I-O**). Together, these results indicate that gap junction coupling and sodium channel activity drive spontaneous synchronous calcium dynamics in iNeuron-iAstrocyte co-cultures.

**Figure 6.**
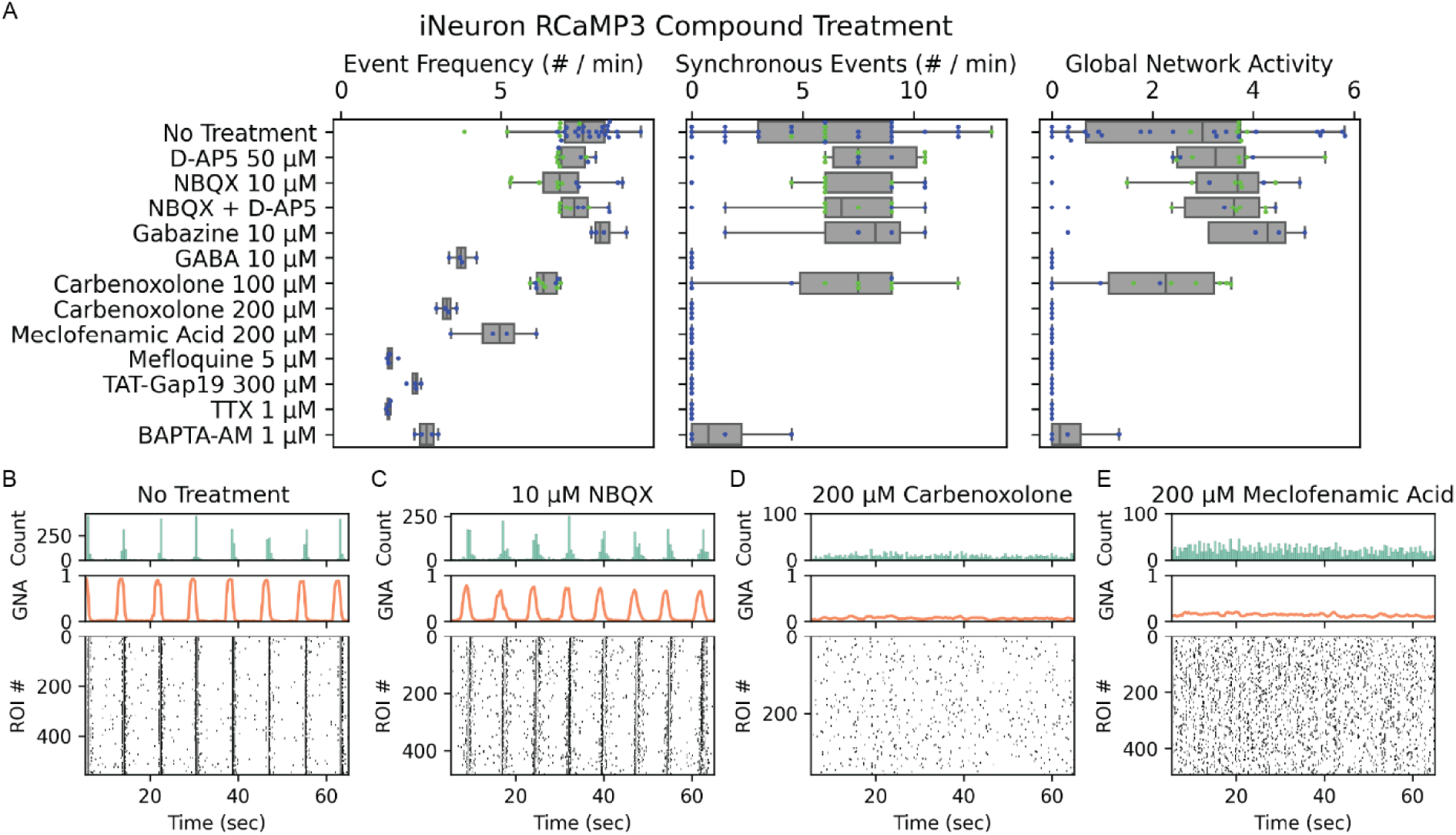
Pharmacological inhibition of gap junctions disrupts synchronous calcium activity. (A) Quantification of calcium average single-neuron event frequency, synchronous event frequency, and GNA in co-cultures with RCaMP3 iNeurons and H1-NBS9 iAstrocytes in NBM-A g10 medium across pharmacological treatments. (N=4-30 replicates per condition from 1-2 differentiations; replicates are colored by batch) (B-E) Raster plots showing calcium event activity in co-cultures with (B) no treatment or treatment with (C) NBQX, (D) Carbenoxolone, (E) Meclofenamic Acid.

To establish a genetically tractable model to investigate the mechanisms underlying synchronized neuronal activity in our co-cultures, we generated clonal iNeuron lines with CRISPRi-iRFP or CRISPRi-Halo. To validate CRISPRi activity in iNeurons, we selected genes encoding proteins localized to synaptic vesicles (*SYN1*; *SYP*), presynaptic boutons (*BSN*), and postsynaptic densities (*HOMER1; DLG4* encoding PSD95). We validated KD in co-cultures with mouse astrocytes at ccD 42 when synaptic marker expression and electrical activity have been reported^18^. We observed efficient transduction of iNeurons with LV one day after co-plating cells and tested monoclonal antibodies against SYN1, SYP, BSN, PSD95, and HOMER1. We found that the IF signal for each target was reduced in KD conditions, albeit with higher background staining for PSD95 and HOMER1 antibodies (**Fig S11A-B**). IF analysis showed that CRISPRi KD effectively reduced SYP protein levels in both CRISPRi-iRFP and CRISPRi-Halo iNeurons co-cultured with iAstrocytes at ccD 14 (**Fig S12A**). Together, these results demonstrate efficient, long-term CRISPRi-mediated KD of synaptic proteins in iNeurons in co-cultures.

Gap junction-mediated synchronization may involve neuronal connexin 45 (*GJC1*), connexin 47 (*GJC2*), or connexin 36 (*GJD2*), which are expressed by iNeurons (**Fig S12B**). To identify neuronal mediators of network synchronization, we introduced RCaMP3 into a CRISPRi-Halo clonal neuronal line. These CRISPRi-Halo + RCaMP3 iNeurons were then co-cultured with NBS9 iAstrocytes, followed by arrayed lentiviral KD of connexins *GJC1*, *GJC2*, and *GJD2* (**Fig S12C**). Neuronal CRISPRi KD of *GJC1*, but not *GJC2* or *GJD2*, reduced average neuron event frequency, frequency of synchronous events, and GNA (**Fig 7B-J, Video S6**). Knockdown of *GJC1* was validated by qPCR (**Fig S12D**). These results support a model in which neuronal gap junctions, specifically connexin 45, coordinate early network activity of iNeurons co-cultured with iAstrocytes, a novel mechanism of *in vitro* human neuronal synchronization that may be useful for probing the role of gap junctions in neuronal network formation.

**Figure 7.**
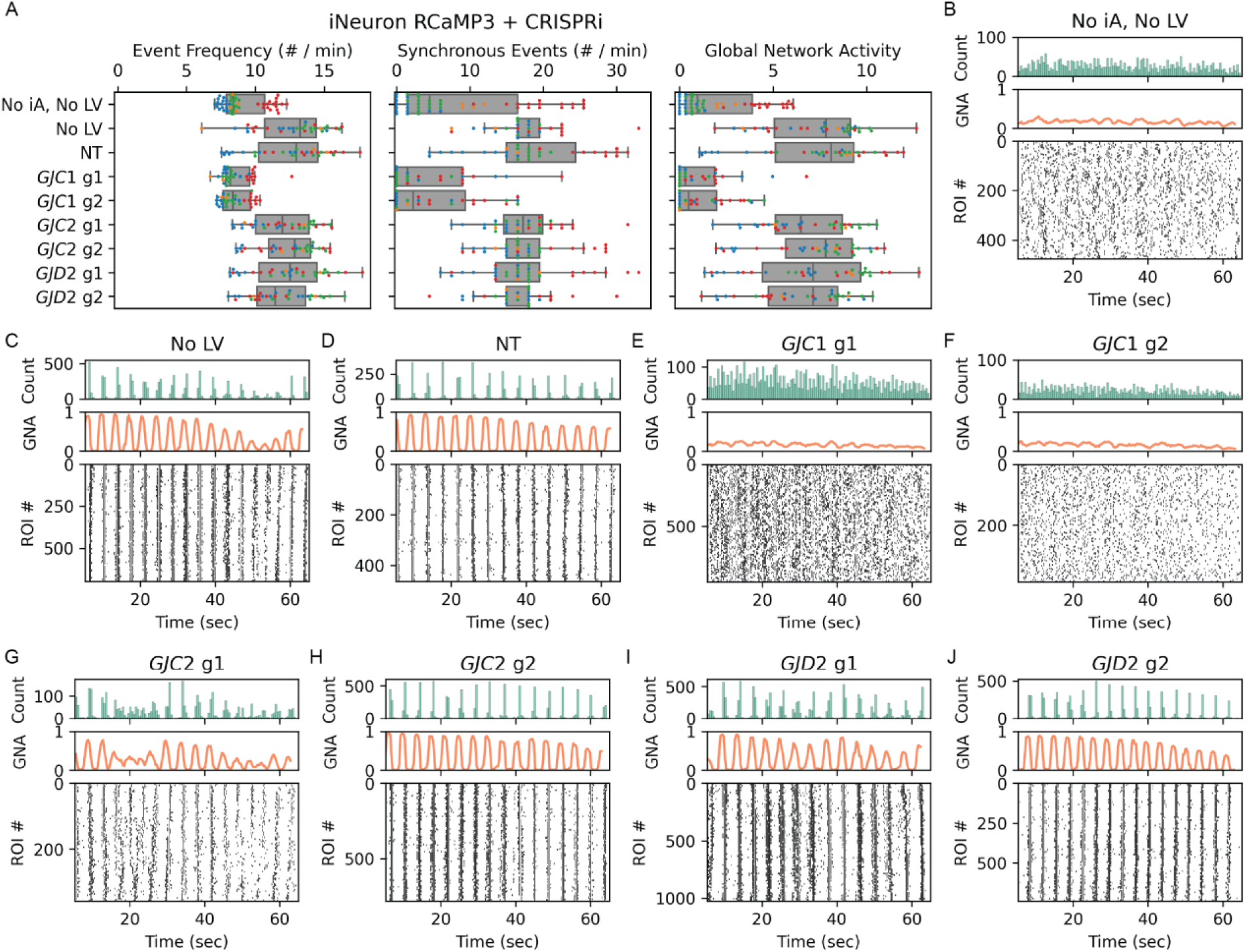
CRISPRi knockdown of GJC1 disrupts synchronous calcium activity. (A) Quantification of calcium average single-neuron event frequency, synchronous event frequency, and GNA in co-cultures with CRISPRi + RCaMP3 iNeurons and H1-NBS9 iAstrocytes on ccD 9-10 with KD of gap junctions *GJC1*, *GJC2*, or *GJD2*. (N=32-34 replicates per condition from 4 differentiations; replicates are colored by batch) (B-J) Raster plots of RCaMP3 calcium activity in co-cultures (B) no astrocyte (C) no LV, or (D) non-targeting controls, or after CRISPRi KD (E-F) *GJC1*, (G-H) *GJC2*, or (I-J) *GJD2*.

## Discussion

We present an optimized and scalable strategy for rapid generation of iAstrocytes from hPSCs. Genomic safe harbor knock-in of astrogenic transcription factors combined with dual chemical selection enables efficient, reproducible differentiations across multiple male and female hESC and iPSC lines. Comparison of NFIB-SOX9 and NFIA-NFIB-SOX9 induction revealed that both TF combinations yielded iAstrocytes expressing canonical astrocyte markers and secreting factors involved in neuronal development. Transcriptomic comparison with existing iAstrocyte models indicates that our iAstrocytes are broadly similar to other 2D models, though additional time or cues, such as 3D culture conditions or co-culture with neurons, may be required to reach a more mature state. Using genomically integrated CRISPRi machinery^41,49^ and lentiviral guide RNA delivery, we achieved efficient gene expression knockdown in both differentiated iAstrocytes and iNeurons. We validated antibodies against key proteins expressed by each cell type, supporting the system’s utility for systematic functional genomics in defined human cell types. Together, these advances establish a scalable and genetically tractable platform for studying human neuron-glia interactions across diverse genetic backgrounds. iAstrocytes can be generated within one week for cryopreservation or direct use in co-cultures, facilitating high-throughput studies. The robustness of this approach across diverse hPSC lines further supports its application to the investigation of human-specific genetic influences on astrocyte function and neuron-glia interactions in health and disease. Compared to existing TF-based induction methods^30,32–34^, our approach offers advantages in biosafety, scalability, and reproducibility on diverse genetic backgrounds.

Astrocyte dysfunction and reactivity has been implicated in a variety of neurodevelopmental and neurodegenerative disease pathologies^85^. In disease contexts, astrocytes upregulate complement components C4 and C3^5,12,86^ and downregulate genes involved in synaptic support^7^, and astrocytes *in vitro* respond to interferon and cytokine treatments by upregulating C4 and C3^10,12,24,86^ and modulating the function of co-cultured neurons^12,60^. However, a fully human model to investigate how astrocyte-derived complement and other reactive astrocyte states influence neuronal and synaptic function has been lacking. Here, we characterized iAstrocyte responses to inflammatory cues under serum-free conditions and observed stimulus-specific reactive phenotypes consistent with those previously reported in other induced^24,27^ and primary astrocytes^12^. iAstrocytes upregulated C4 and MHC-I following interferon treatment, and C3 and ICAM1 following IL1α and TNFα co-treatment. Notably, C1q treatment did not individually or additively induce these iAstrocyte responses. CRISPRi-mediated repression efficiently reduced the induction of C4, C3, and ICAM1. This platform is thus poised to enable functional dissection linking astrocyte reactivity to neuronal and synaptic dysfunction. Future studies will examine how cytokine treatments or other manipulations of astrocytic complement expression modulate neuronal network formation or activity in different co-culture conditions.

Support of synapse formation and function is a primary role of astrocytes in the developing and mature brain, and disruption of neuron-astrocyte communication is implicated in neurodevelopmental and neurodegenerative disorders^1–9^. Astrocytes are the primary cellular source of two proteins, APOE and complement C4, that are centrally involved in synapse development and disease and confer disease risk through human-specific protein variants that cannot easily be studied functionally in non-human models. Our iAstrocytes express both APOE and C4, in addition to thrombospondins, glypicans, and other synapse-supporting factors, enabling molecular functional studies of human-specific variants and their impact on astrocyte-neuron interactions. To investigate iAstrocyte support of human neurons *in vitro*, we optimized co-culture conditions for attachment, survival, and fluorescent calcium indicator recordings. We focused on NFIB-SOX9 iAstrocytes which exhibited superior viability in co-cultures compared to those produced with addition of NFIA. Calcium imaging demonstrated that co-culture with mouse astrocytes, human fetal astrocytes, or NFIB-SOX9 iAstrocytes from five independent hPSC lines each promoted synchronization of spontaneous neuronal activity not observed in iNeuron monocultures. Pharmacological perturbation revealed that iAstrocytes supported gap junction-mediated neuronal synchronization, in contrast to mouse astrocytes which supported AMPA receptor-mediated synchronization at a similar co-culture stage. CRISPRi KD of *GJC1* (connexin 45) in iNeurons disrupted iAstrocyte-dependent synchronized activity. These findings demonstrate the use of iAstrocytes to interrogate the molecular mechanisms by which astrocytes support neuronal development and connectivity. These findings identify neuronal connexin 45 as a critical mediator of astrocyte-supported network synchronization and suggest that astrocytes regulate an early developmental stage of circuit formation that precedes mature excitatory synaptic transmission. Together, these findings establish a tractable human model for dissecting the molecular mechanisms through which astrocytes shape neuronal network development and function.

Gap junction-mediated neuronal synchronization appears mechanistically distinct from the AMPA receptor-driven activity observed in co-cultures with mouse astrocytes and other iAstrocyte models^18,75–77^. Calcium waves and dye coupling have been observed in cortical, thalamic, and retinal neurons at late gestational and perinatal stages and are abolished by gap junction inhibitors^78–80,87–89^. Neuronal *GJC1* expression peaks at this same stage in rodents and humans before declining during adolescence^90–92^. Support for gap junction-mediated calcium wave propagation may represent an early function of astrocytes that shapes neuronal network development. Our iAstrocyte-iNeuron co-cultures are, to our knowledge, the first *in vitro* model of gap junction-mediated neuronal network activity and provide a platform to investigate how astrocytes support this early developmental form of neuronal circuit activity. Our hPSC-derived co-culture system thus provides a tractable platform to dissect astrocyte contributions to early neuronal network development and plasticity, capturing features of fetal circuit maturation not recapitulated in co-cultures with postnatal primary mouse astrocytes.

Significant gaps remain in our understanding of how astrocytes regulate neuronal development or disease-related dysfunction. Our fully human, genetically tractable, and scalable co-culture system is well positioned to address these questions. Using genetic, pharmacologic, and cytokine perturbations, we can systematically test how iAstrocyte-derived factors influence early gap junction-mediated network activity and later AMPA receptor-driven activity, including how disruptions to this developmental progression may contribute to circuit-level dysfunction in neurodevelopmental or psychiatric disorders. This system further enables cell type-specific interrogation of disease-linked genetic variation or molecular mechanisms driving synapse or network dysfunction. Overall, this work establishes a scalable platform for the study of human neuron-astrocyte interactions.

## Supporting information

Table S1_This study

## Author Contributions

N.M., M.B.J., and B.S. designed the study. N.M., J.L.D., L.L. cloned plasmids. N.M., M.J.D. cultured, modified, and differentiated stem cells. N.M., M.J.D., H.J. performed immunofluorescence stains and analysis. N.M. performed RNA data analysis. N.M., M.J.D. collected calcium activity measurements and A.G.-R. analyzed data with supervision from A.J.G.. S.D. performed C4A/B antibody validation. N.M. performed reactive astrocyte assays. N.M. assembled figures. N.M., M.B.J., and B.S. wrote the manuscript.

## Acknowledgements

We thank members of Stevens laboratory for numerous discussions and feedback; Isaac Canals, William C. Skarnes, Erik Ullian, Marius Wernig, and Takeshi Uenaka for feedback on experimental results; Maria Alimova for management of Opera Phenix High-Throughput imaging system at Broad Institute, funded by S10 Grant NIH OD-026839-01; members of Eroglu, Allen, Nehme, and Khurana laboratories for feedback on astrocyte culture methods, Ralda Nehme and Ross D. Jones for manuscript comments, and Krishna Narayanan for administrative support. We also thank Kathleen Worringer for AAVS1-NGN2 donor plasmid, Takeshi Uenaka for TK4 plasmids, and Michael Ward for S41 plasmid. This work was supported by Stanley Center for Psychiatric Research at Broad Institute, NIMH P50 MH112491 (to B.S.), Sigrid Juselius Foundation (to H.J.), Instrumentarium Science Foundation (to H.J.), Howard Hughes Medical Institute (to B.S.), 5T32AG222-30 (to N.M.), 1F32AG079666-01 (to N.M.).

## Materials & Correspondence

Correspondence and requests for materials should be addressed to N.M., or B.S..

## Supplementary Figures

**Supplementary Figure S1.**
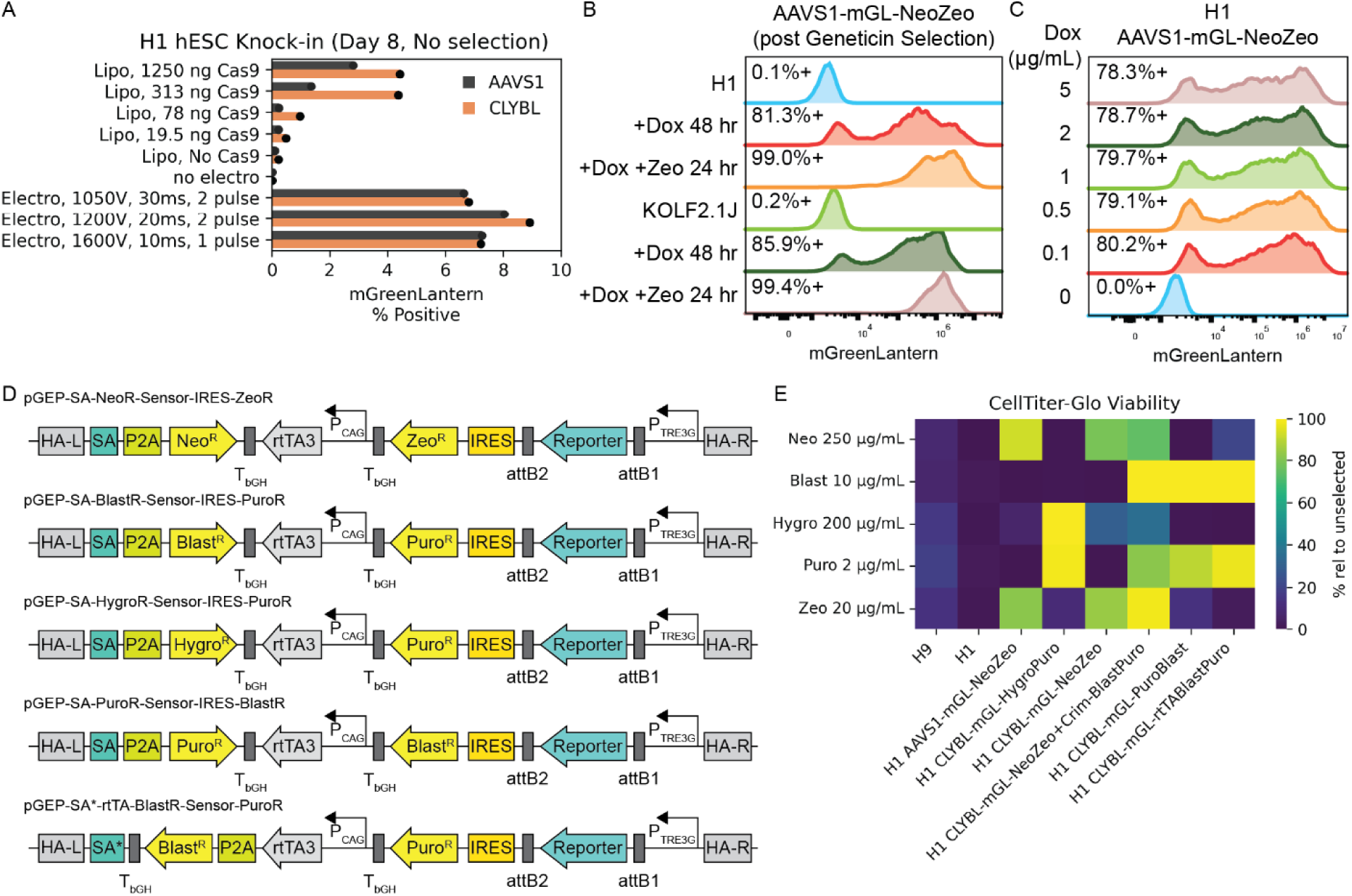
Safe Harbor knock-in and induced expression optimization. (A) Percentage of H1 hESCs positive for mGreenLantern, 8 days after electroporation or lipofection with donor DNA (AAVS1-mGreenLantern-NeoZeo or CLYBL-mGreenLantern-NeoZeo) and Cas9 RNP targeting AAVS1 or CLYBL. (B-C) Flow cytometry histogram of mGreenLantern fluorescence in H1 and KOLF2.1J hPSCs with AAVS1-mGreenLantern-NeoZeo knock-in with (B) 48 hours 500 ng/mL dox or 48 hours 500 ng/mL dox + 24 hour 20 μg/mL zeocin treatment or (C) 24 hours dox dosages from 100 - 5000 ng/mL. (D) Schematics for safe harbor knock-in vectors with dual selection markers: SA-NeoR-IRES-ZeoR (NeoZeo), SA-BlastR-IRES-PuroR (BlastPuro), SA-HygroR-IRES-PuroR (HygroPuro), SA-PuroR-IRES-BlastR (PuroBlast), and SA*-rtTA-P2A-BlastR-IRES-PuroR (rtTABlastPuro). SA: Splice acceptor, SA: Splice acceptor with stop code, HA: Homology arm, P2A, T2A: self-cleaving peptides, rtTA3: reverse tetracycline-controlled transactivator 3, P_CAG: CAG promoter, P_TRE2G: tet-response element 2nd gen, IRES: internal ribosomal entry site, attB1, attB2: gateway cloning scars, T_bGH: bovine growth hormone terminator. (E) Relative viability of hPSCs, cultured for 72 hours with each antibiotic. H1 and H9 parental hESCs and H1 hESCs with knock-in of BlastPuro (BP), NeoZeo (NZ), PuroBlast (PB), or rtTABlastPuro (RBP) constructs. hPSC lines include H1 and H9 hESC parental lines and H1 lines with CLYBL or AAVS1 knock-in of vectors with mGreenLantern or Crimson expression. hESCs were cultured for 72 hours with E8 and geneticin, blasticidin, hygromycin, puromycin, or zeocin selection. Intensity measurements are normalized to unselected controls for each line.

**Supplementary Figure S2.**
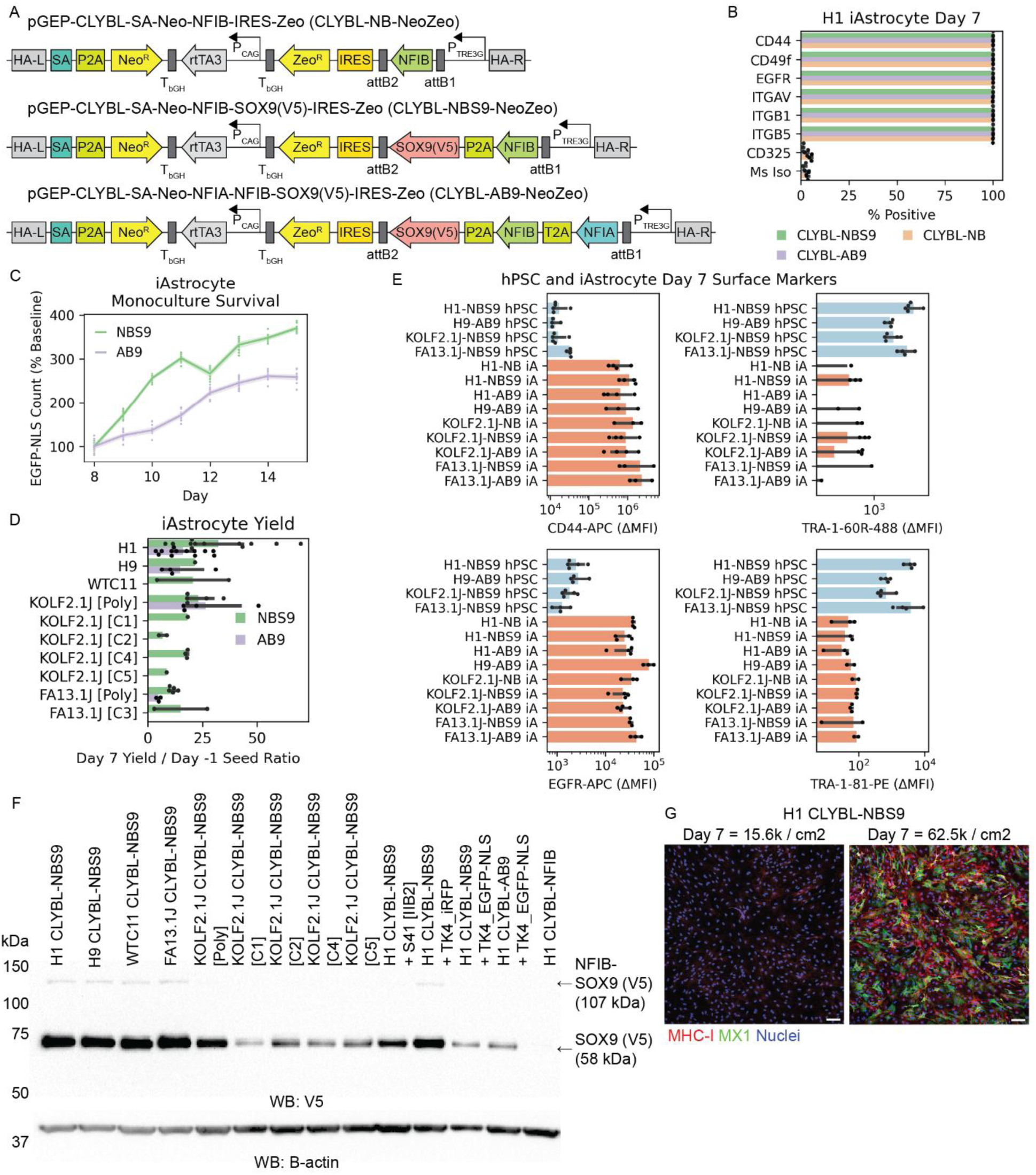
Safe harbor astrocyte model validation. (A) Vector schematic for geneticin-selectable knock-in, dox-inducible expression of NFIB (NB), NFIB-SOX9 (NBS9), or NFIA-NFIB-SOX9 (AB9) with zeocin-selection. (B) Flow cytometry quantification of iAstrocyte surface markers across H1 CLYBL-NB, NBS9, and AB9 lines. (n=3 replicates per condition from 1 differentiation) (C) Quantification of nuclear area of H1 CLYBL-NBS9 or AB9 iAstrocytes labeled with EGFP-NLS between days 8 and 15. (D) Quantification of iAstrocyte cell count ratio comparing iAstrocyte harvest count on day 7 with hPSC seeding count on day -1. (n=1-11 differentiations per line) (E) Flow cytometry quantification of iAstrocyte surface markers across H1, H9, KOLF2.1J, and FA13.1J day 7 iAstrocytes and uninduced hPSCs. (n=3-4 differentiations per line) (F) Western blot detection of V5 epitope tag on inducible SOX9 and B-actin in cell lysates from CLYBL-NB, NBS9, or AB9 iAstrocytes on day 7. (G) IF images of H1 CLYBL-NBS9 iAstrocytes on day 21 after seeding at 1.56e4 or 6.25e4 cells / cm^2^ on day 7. Red = MHC-I, Green = MX1, Blue = Nuclei. Scale = 100 μm.

**Supplementary Figure S3.**
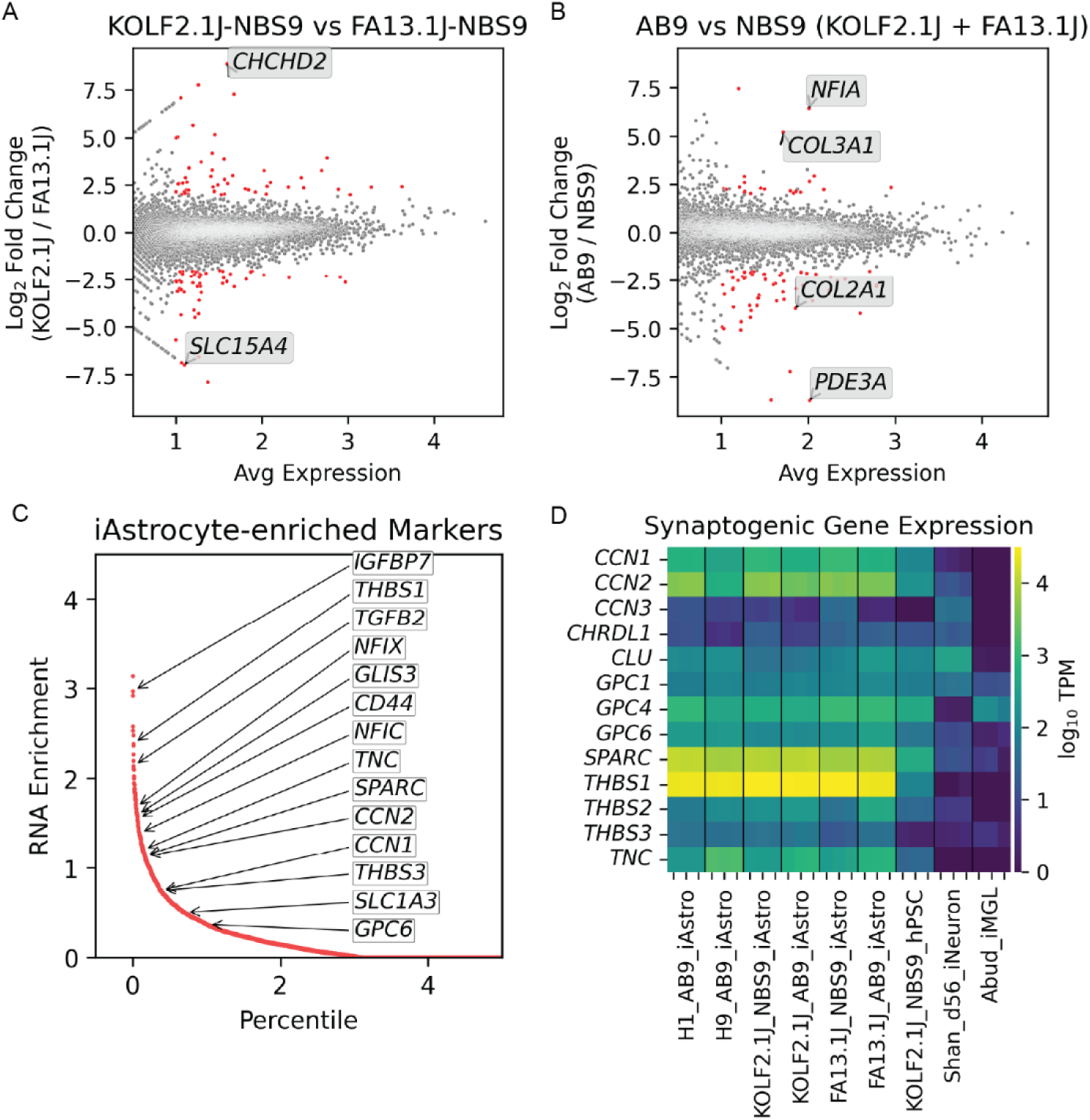
iAstrocyte RNA-seq comparison with published models. (A) Volcano plot for differentially expressed genes between KOLF2.1J-NBS9 and FA13.1J-NBS9 iAstrocytes. (B) Volcano plot for differentially expressed genes between NBS9 and AB9 iAstrocytes for KOLF2.1J and FA13.1J lines. (C) Enrichment scores for genes expressed in NBS9 or AB9 iAstrocytes compared to hPSCs, iNeurons, and iMGLs. (D) Heatmap showing expression of select synaptogenic genes across iAstrocytes, iNeurons, iMGLs, and hPSCs. Colors correspond to log10 transcripts per million (TPM).

**Supplementary Figure S4.**
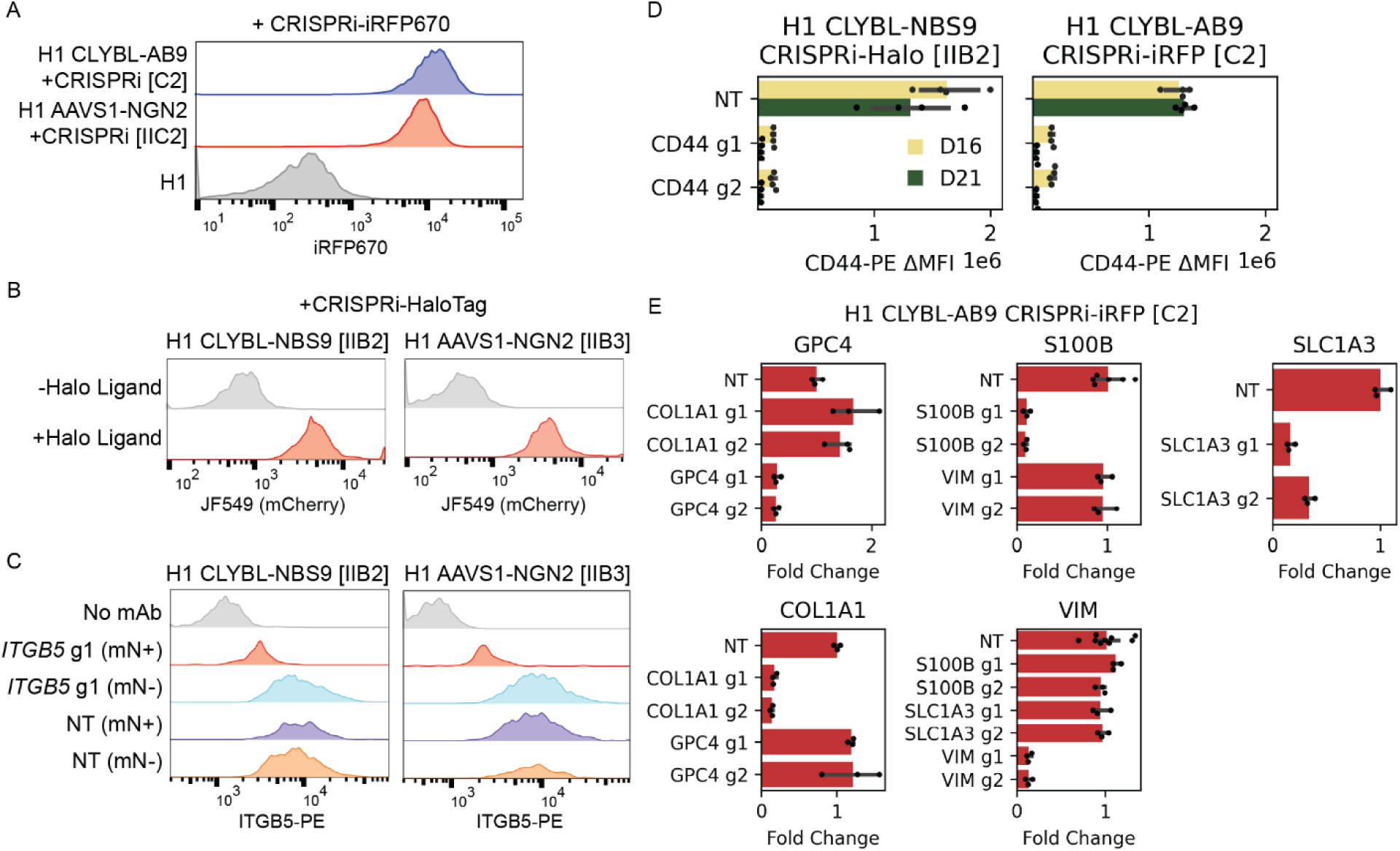
Validation of CRISPRi activity in iAstrocytes. (A) Flow cytometry quantification of iRFP670 expression in H1 CLYBL-AB9 and H1 AAVS1-NGN2 clonal lines with CRISPRi-iRFP670 and non-fluorescent parental H1 line. (B-C) Flow cytometry validation of H1 CLYBL-NBS9 and AAVS1-NGN2 stable clonal lines with CRISPRi-HaloTag. (B) mCherry fluorescent signal with HaloTag Ligand JF549 or (C) ITGB5-PE surface staining on mNeonGreen (mN) positive or negative hPSCs after LV KD targeting *ITGB5* or NT control. (D) Flow cytometry quantification of CD44 expression on days 16 and 21 CRISPRi iAstrocytes (H1-NBS9 CRISPRi-Halo and H1-AB9 CRISPRi-iRFP). iAstrocytes were transduced LV targeting *CD44* or NT. (N=4 replicates per condition from 1 differentiation) (E) qPCR quantification of *COL1A1*, *GPC4*, *S100B*, *SLC1A3*, or *VIM* in H1-AB9 CRISPRi-iRFP iAstrocytes on day 21. iAstrocytes were transduced with LV targeting each gene or NT. (N=3 replicates per condition from 1 differentiation)

**Supplementary Figure S5.**
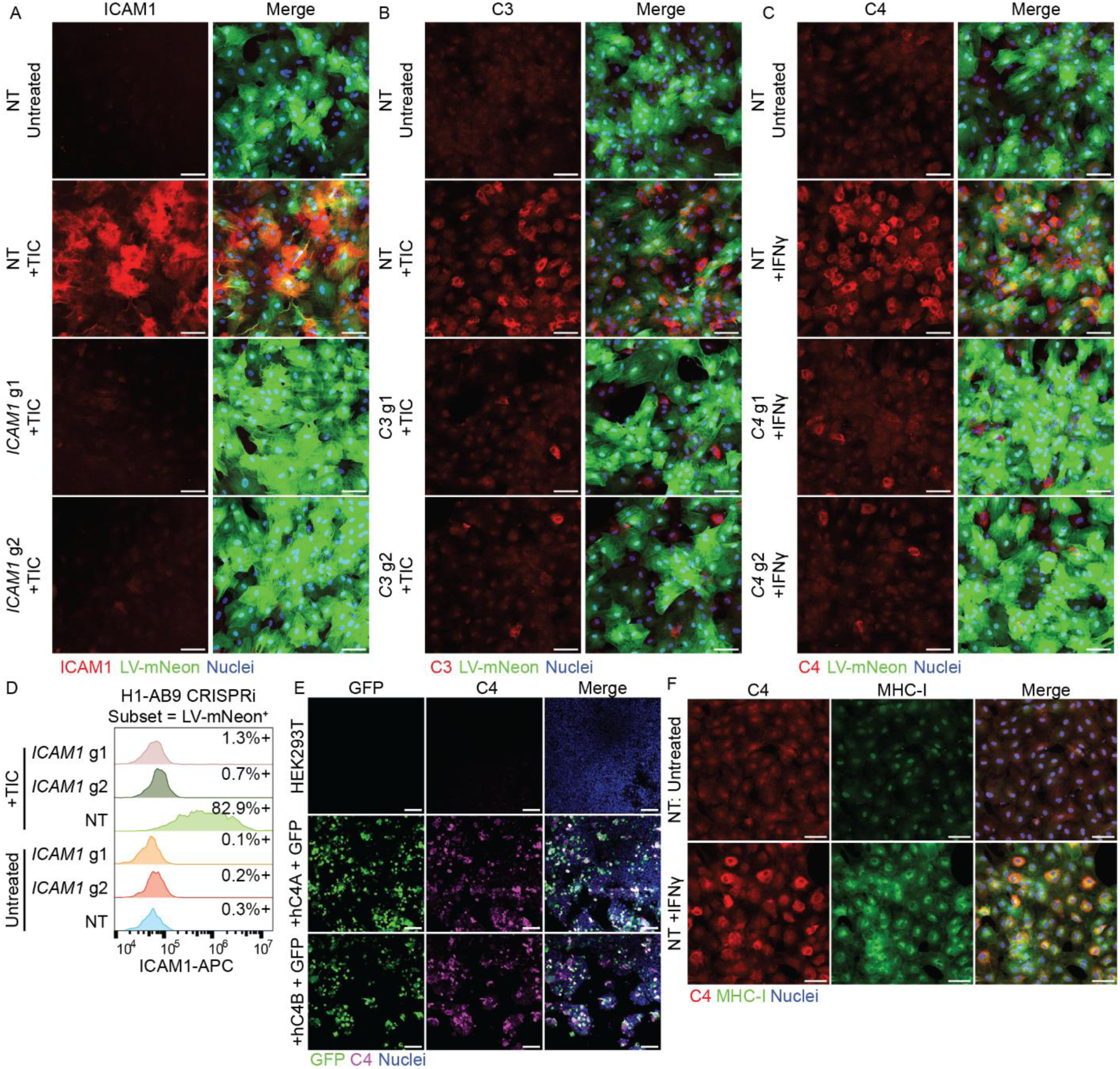
Validation of reactive astrocyte markers using CRISPRi. (A-C) IF images of ICAM1 (A), C3 (B), and C4 (C) in H1-AB9 CRISPRi iAstrocytes with or without TIC or IFNγ treatment and non-targeting (NT) or on-target LV. Red = ICAM1, C3, or C4, Green = mNeonGreen, Blue = Nuclei. Scale = 100 μm. (D) Flow cytometry quantification of ICAM1 expression on day 21 H1-AB9 CRISPRi iAstrocytes treated with TIC and transduced with LV targeting *ICAM1* or NT. (E) IF images of C4 protein in HEK293T cells with piggyBac expression of EGFP and hC4A or hC4B. Green = EGFP, Magenta = C4, Blue = Nuclei. Scale = 100 μm. (F) Representative IF images of C4 and MHC-I in day 21 H1-AB9 CRISPRi iAstrocytes treated with IFNγ for data shown in **Fig 3C**. Red = C4, Green = MHC-I and iRFP670, Blue = Nuclei. Scale = 100 μm.

**Supplementary Figure S6.**
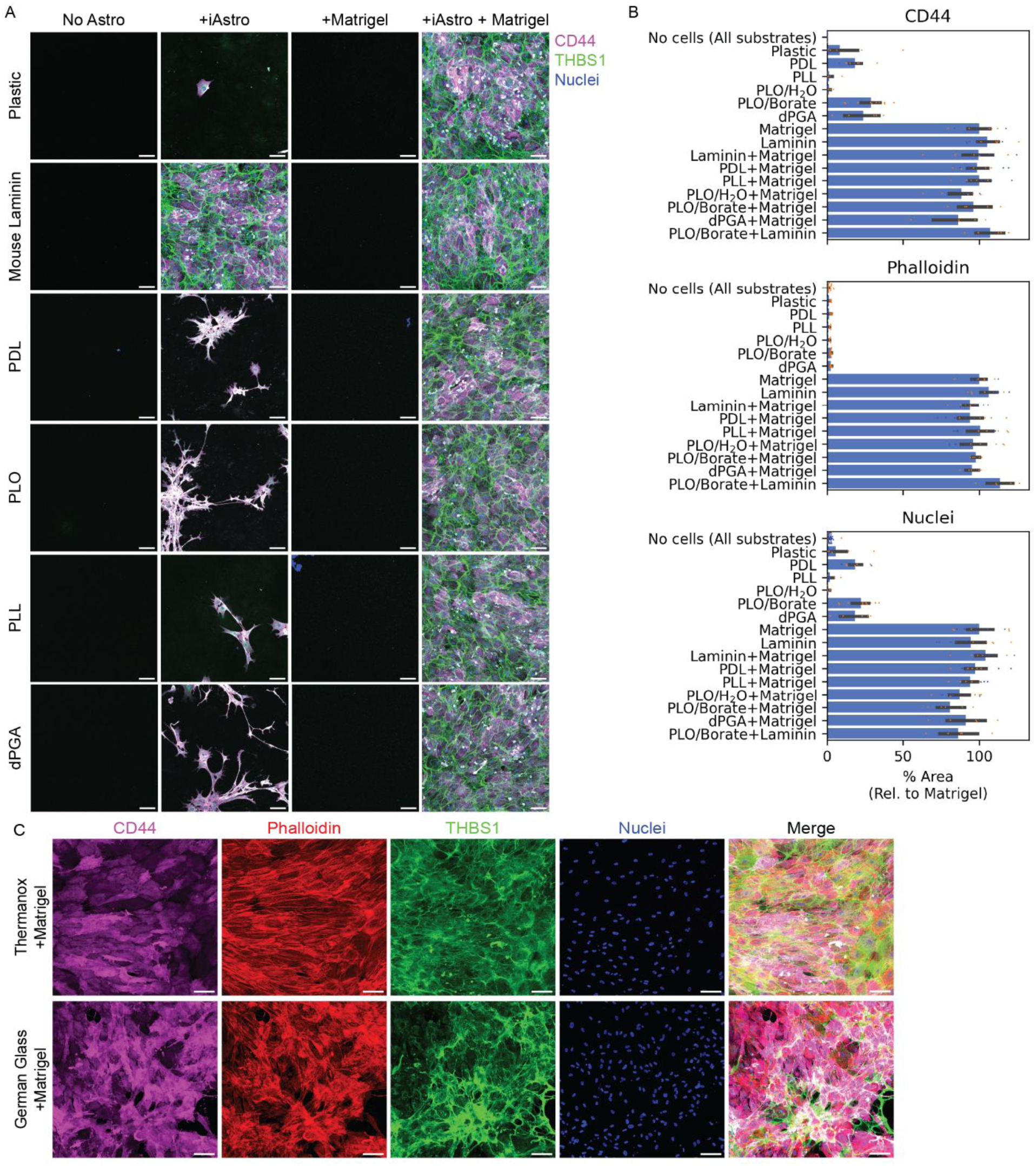
iAstrocyte thrombospondin deposition across vessel coating substrates. (A) IF images of CD44 and THBS1 on plastic vessels coated with mouse laminin, poly-D-lysine (PDL), poly-L-lysine (PLL), poly-L-ornithine (PLO), or dendritic polyglycerol amine (dPGA), mouse laminin, with or without matrigel and H1-NBS9 iAstrocytes on day 21. Green = THBS1, Magenta = CD44, Blue = Nuclei. Scale = 100 μm. (B) Quantification of CD44, Phalloidin, and THBS1 area across conditions shown in (A). (N=5-14 replicates per condition from 2 differentiations) (C) IF images of VIM and THBS1 from day 21 H1-NBS9 iAstrocytes cultured on Thermanox or German Glass coverslips with matrigel coating. Red = Phalloidin, Green = THBS1, Magenta = CD44, Blue = Nuclei, Scale = 100 μm.

**Supplementary Figure S7.**
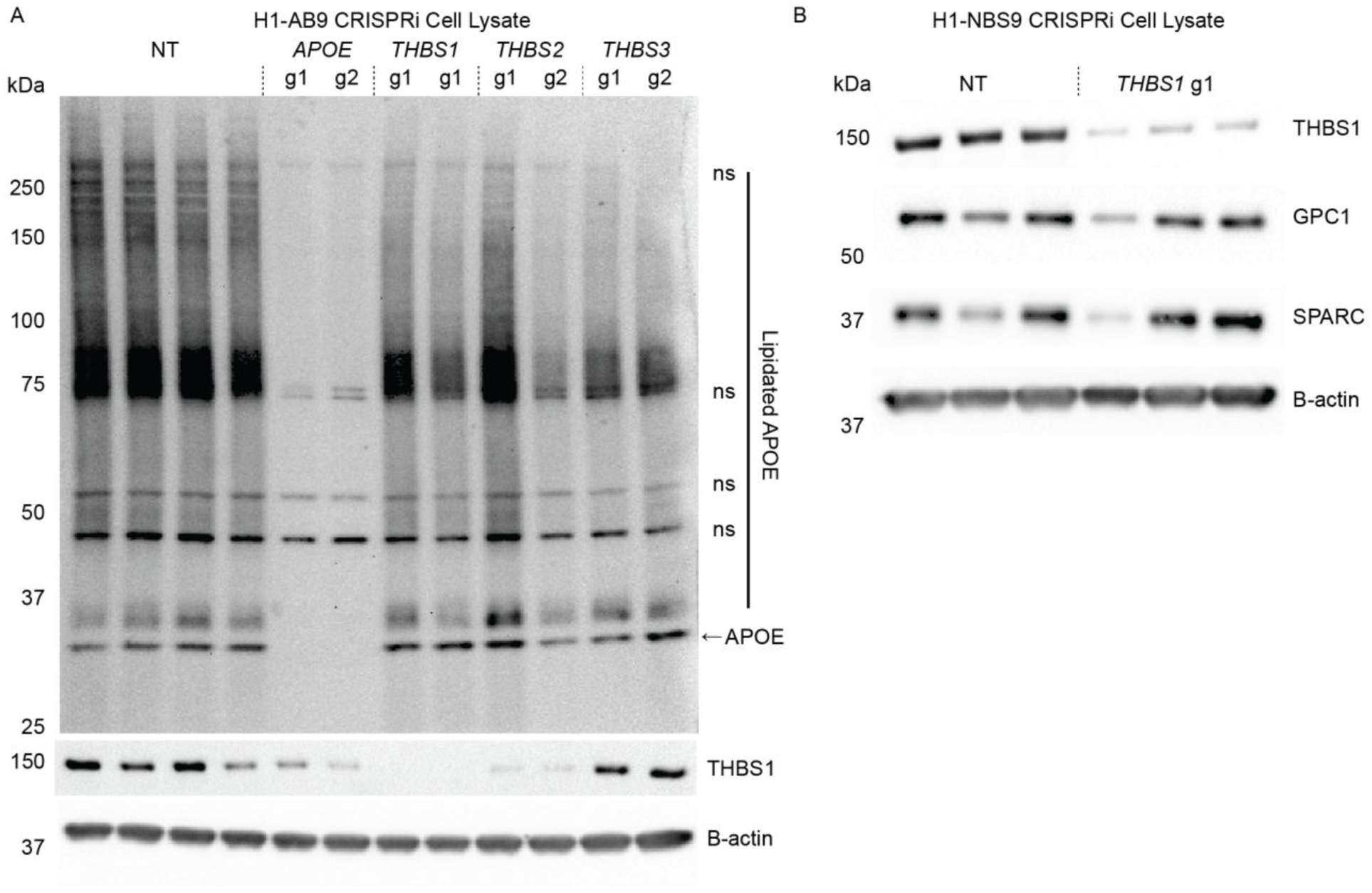
iAstrocyte cell lysate protein validation. (A) Western blot detection of APOE, THBS1, and B-actin in cell lysate from H1-AB9 CRISPRi iAstrocytes on day 21 after LV knockdown of *APOE*, *THBS1*, *THBS2*, *THBS3*, or NT. (B) Western blot detection of THBS1, GPC1 and SPARC, and B-actin in cell lysate from H1-NBS9 CRISPRi iAstrocytes on day 21 after LV knockdown of *GPC1*, *GPC2*, *GPC4*, *GPC6*, or NT.

**Supplementary Figure S8.**
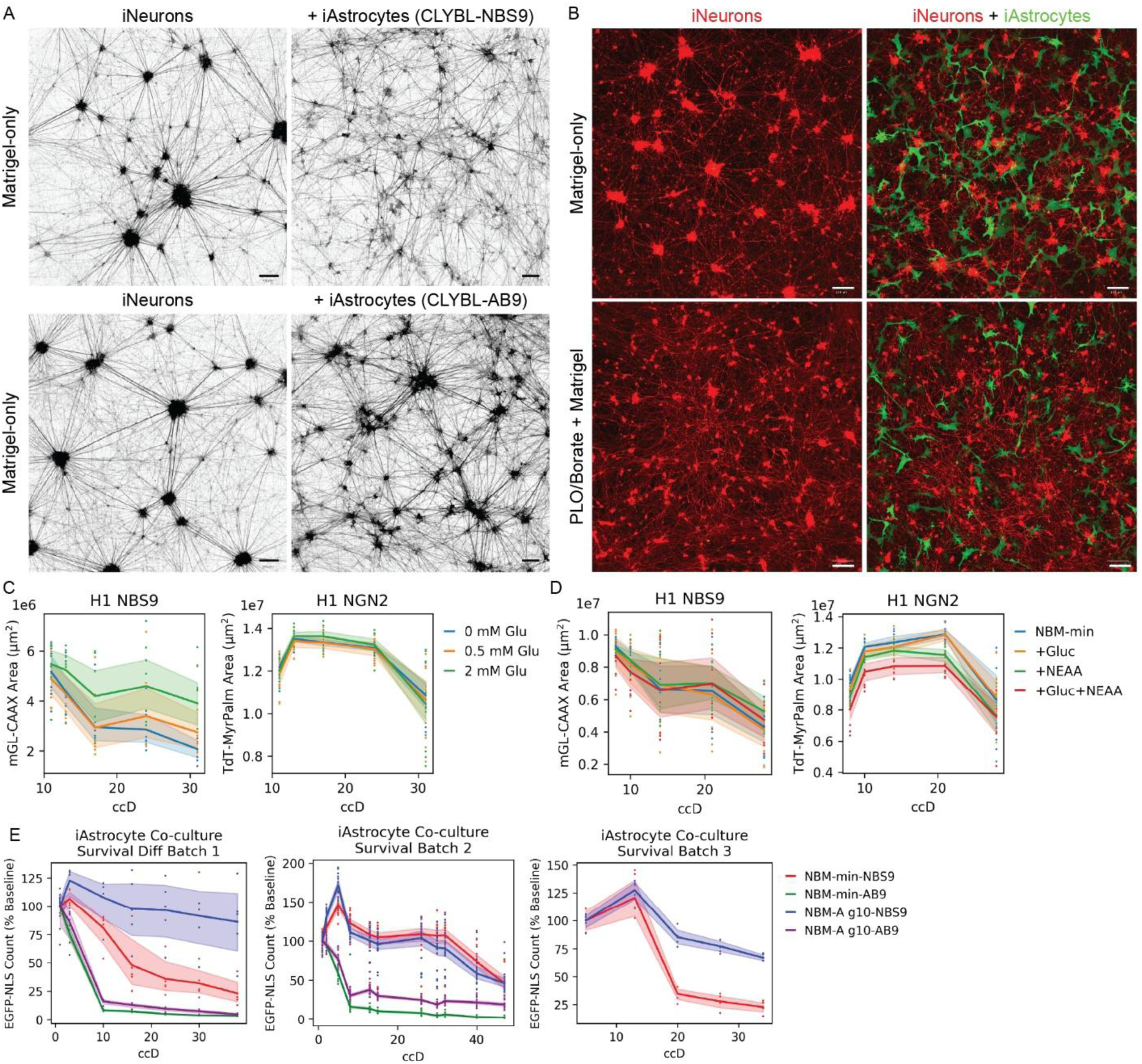
Optimization of co-culture attachment and survival conditions. (A) IF images comparing iNeurons labeled with iRFP670-CAAX on vessels coated with matrigel in monoculture (left) or co-culture with H1-NBS9 or AB9 iAstrocytes (right) on ccD 14. Black = iRFP670. Scale = 100 μm. (B) IF images of RCaMP3 iNeurons in monoculture (left) or co-culture with H1-NBS9 iAstrocytes labeled with iRFP-CAAX (right) on ccD 14 on vessels coated with matrigel-only or PLO followed by matrigel. Red = iNeurons, Green = iAstrocytes. Scale = 100 μm. (C-D) Quantification area coverage in co-cultures with H1-NBS9 iAstrocytes labeled with mGreenLantern-CAAX and iNeurons labeled with TdTomato-MyrPalm with (C) varying concentrations of Glutamax with glucose and NEAA present or (D) varying combinations of glucose and NEAA in absence of Glutamax. (N=8-12 replicates per condition from 1 differentiation for each panel) (E) Quantification of nuclear area of H1-NBS9 or AB9 iAstrocytes labeled with EGFP-NLS in co-culture with iNeurons in NBM-min or NBM-A g10 media between ccDs 1 and 35. (N=25-40 replicates per condition from 3 (NBS9) or 2 (AB9) differentiations)

**Supplementary Figure S9.**
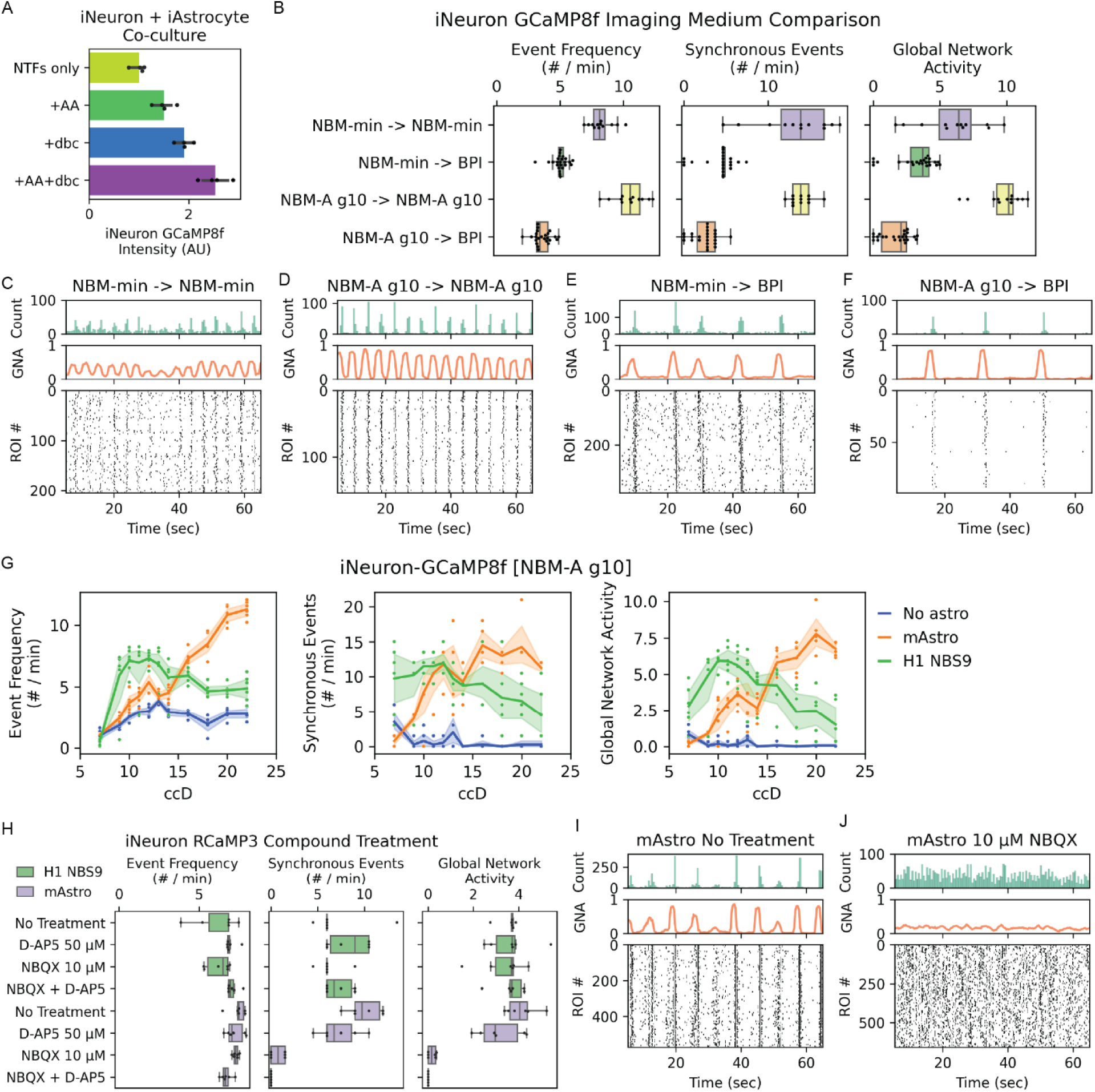
Optimization of co-culture neuronal calcium activity. (A) IF quantification of GCaMP8f intensity in iNeurons co-cultured with H1-NBS9 iAstrocyte on ccD 6 with neurotrophic factors (NTFs) alone or with combinations of ascorbic acid (AA) and dbcAMP (dbc). (N=4 replicates per condition from 1 differentiation) (B) Quantification of calcium average single-neuron event frequency, synchronous event frequency, and GNA in co-cultures with GCaMP8f iNeurons and H1-NBS9 iAstrocytes across media conditions on day 14. (N=12 replicates per condition from 1 differentiation) (C-F) Raster plots of calcium activity in co-cultures with GCaMP8f iNeurons and H1-NBS9 iAstrocytes using base media: (C) NBM-min, (D) NBM-A g10, or (E-F) after switching to each to BrainPhys Imaging medium (BPI). (G) Quantification of calcium average single-neuron event frequency, synchronous event frequency, and GNA in GCaMP8f iNeurons between ccDs 7 and 22 in monoculture or co-culture with H1-NBS9 iAstrocytes in NBM-A g10 medium. (N=6 replicates per condition from 1 differentiation) (H) Quantification of calcium average single-neuron event frequency, synchronous event frequency, and GNA in RCaMP3 iNeurons in co-culture with H1-NBS9 or mouse astrocytes in NBM-A g10 medium across pharmacological treatments. (N=6 replicates per condition from 1 differentiation; NBS9 conditions are also shown in **Fig 6E**) (I-J) Raster plots of iNeuron RCaMP3 calcium activity in co-culture with mouse astrocytes after (I) no treatment or (J) treatment with NBQX.

**Supplementary Figure S10.**
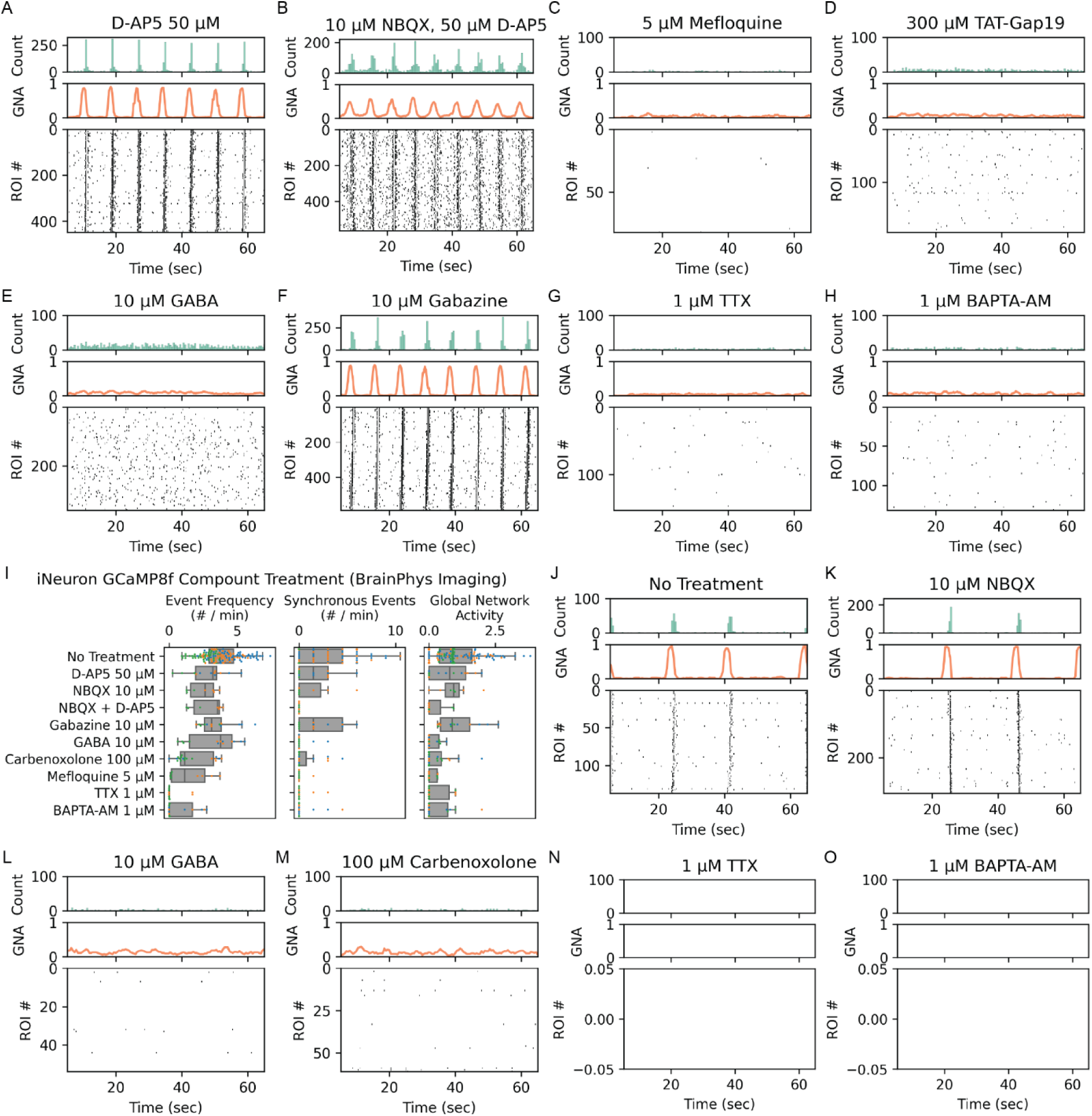
Co-culture pharmacology raster plots. (A-H) Raster plots of iNeuron RCaMP3 calcium activity in co-cultures with H1-NBS9 iAstrocytes after treatment with (A) D-AP5, (B) NBQX and D-AP5, (C) Mefloquine, (D) TAT-Gap19, (E) GABA, (F) Gabazine, (G) TTX, (H) BAPTA-AM. (I) Quantification of calcium average single-neuron event frequency, synchronous event frequency, and GNA in co-cultures with GCaMP8f iNeurons and H1-NBS9 iAstrocytes in BrainPhys Imaging medium across pharmacological treatments. (N=7-137 replicates per condition from 2-3 differentiations; replicates are colored by batch) (J-O) Raster plots of iNeuron GCaMP8f calcium activity in co-cultures with iAstrocytes after (J) no treatment, or treatment with (K) NBQX, (L) GABA, (M) Carbenoxolone, (N) TTX, (O) BAPTA-AM.

**Supplementary Figure S11.**
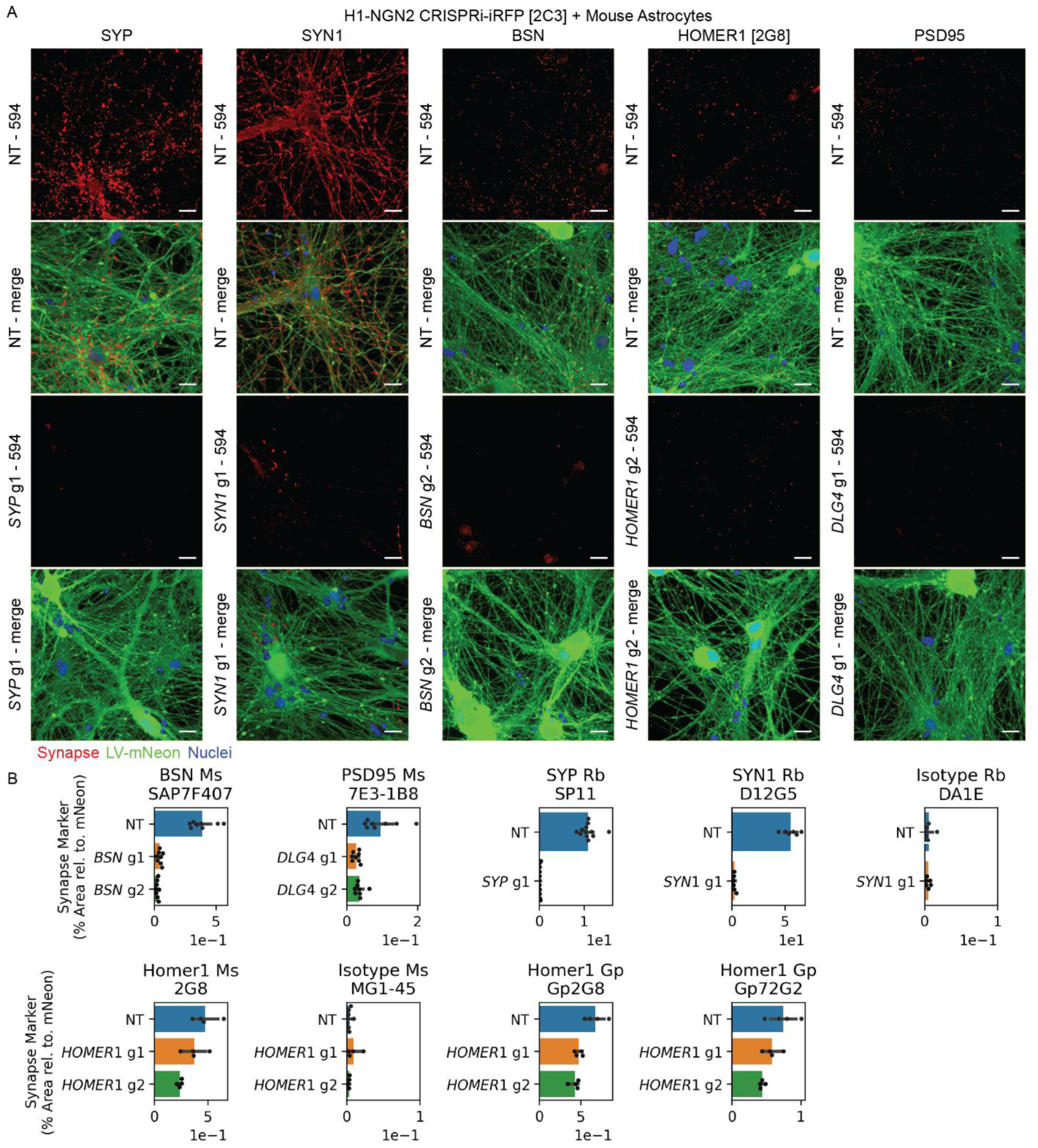
Validation of synaptic antibodies using CRISPRi iNeurons. (A) IF images of synaptophysin (SYP), Synapsin-1 (SYN1), Bassoon (BSN), Homer1, and PSD-95 (DLG4) in CRISPRi iNeurons on ccD 42 after LV transduction targeting each gene or NT. Red = Synapse Protein, Green = mNeonGreen, Blue = Nuclei. Scale = 20 μm.Quantification of synaptic antibody staining area across conditions shown in (A). (N=4-12 replicates per condition from 1 differentiation)

**Supplementary Figure S12.**
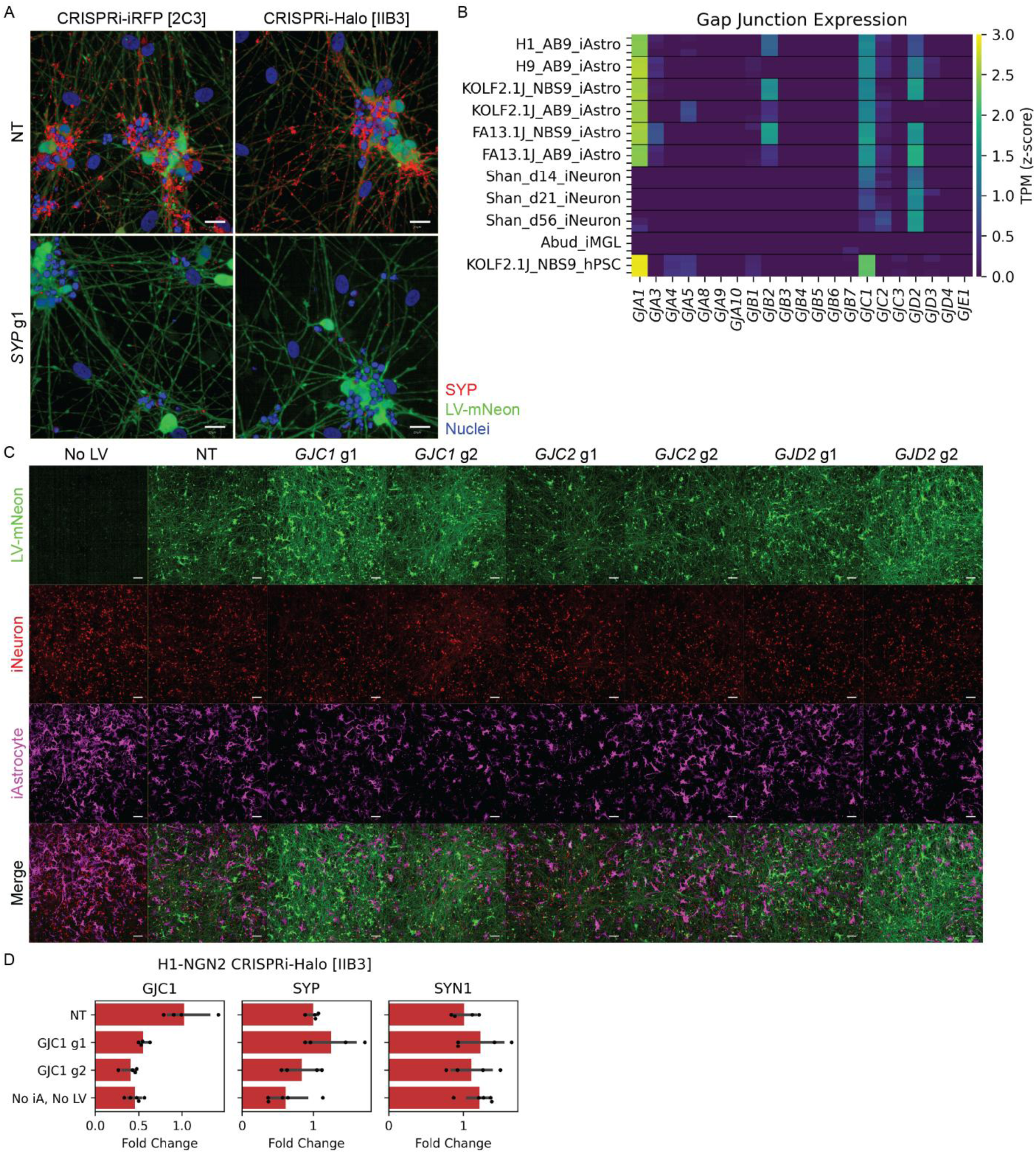
Co-culture neuronal gap junction CRISPRi characterization. (A) IF images of synaptophysin (SYP) in CRISPRi-iRFP or CRISPRi-Halo iNeurons on ccD 14 after transduction with LV targeting *SYP* or NT. Red = SYP, Green = mNeonGreen, Blue = Nuclei. Scale = 20 μm. (B) Heatmap showing expression of gap junction genes across iAstrocytes, iNeurons, iMGLs, and hPSCs. Colors correspond to log10 transcripts per million (TPM). (C) IF images of co-cultures with CRISPRi-Halo + RCaMP3 iNeurons and iAstrocytes labeled with iRFP-CAAX on ccD 10 after LV transduction targeting *GJC1*, *GJC2*, *GJD2*, or NT. Green = mNeonGreen, Red = RCaMP3, Magenta = iAstrocytes. Scale = 100 μm. (D) qPCR quantification of *GJC1*, *SYP*, or *SYN1* fold change normalized to *ACTB* in co-cultures with CRISPRi + RCaMP3 iNeurons and H1-NBS9 iAstrocytes on ccD 11. (N=4-5 replicates per condition from 1 differentiation)

## Supplementary Information

**Table S1.** Bulk RNA-seq data from nine iAstrocyte lines and uninduced hPSC control.

**Table S2.** RNA-seq differential gene expression, cell type enrichment scores, and PCA loadings.

**Table S3.** Cell lines and media reagents used for each figure.

**Video S1.** Calcium activity in RCaMP3 iNeurons co-cultured with H1-NBS9 iAstrocytes in (top) medium without dbcAMP or (bottom) medium with dbcAMP and imaged after medium change to (left) NBM-A g10 without or with dbcAMP, (middle) BPI without dbcAMP, or (right) BPI with dbcAMP.

**Video S2.** Calcium activity in GCaMP8f iNeurons co-cultured with H1-NBS9 iAstrocytes in (top-left) NBM-A g10, (top-right) NBM-min, (bottom-left) NBM-min switched to BPI, or (bottom-right) NBM-A g10 switched to BPI.

**Video S3.** Calcium imaging in RCaMP3 iNeurons in monoculture or co-culture with mouse astrocytes, human fetal astrocytes or five NBS9 iAstrocyte lines (Top, left to right: monoculture, mouse astrocytes, human fetal astrocytes, H1-NBS9; Bottom, left to right: H9-NBS9 KOLF2.1J-NBS9 [C4], FA13.1J-NBS9 [Poly], WTC11-NBS9).

**Video S4.** Simultaneous imaging of calcium activity in (left; magenta) GCaMP8f iNeurons, (middle; cyan) H1-NBS9 RCaMP3 iAstrocytes, and (right) merged channels.

**Video S5.** Calcium activity in RCaMP3 iNeurons co-cultured with H1-NBS9 iAstrocytes with (top-left) no treatment, (top-center) TTX, (top-right) GABA, (bottom-left) Carbenoxolone, (bottom-center) Meclofenamic Acid, or (bottom-right) Mefloquine.

**Video S6.** Calcium imaging in CRISPRi + RCaMP3 iNeurons across conditions with (top-left) no astrocytes or LV, (top-right) with iAstrocytes and NT LV, or (bottom-left, bottom-right) with iAstrocytes and LV targeting *GJC1*.

## STAR★Methods

### Key Resources Table

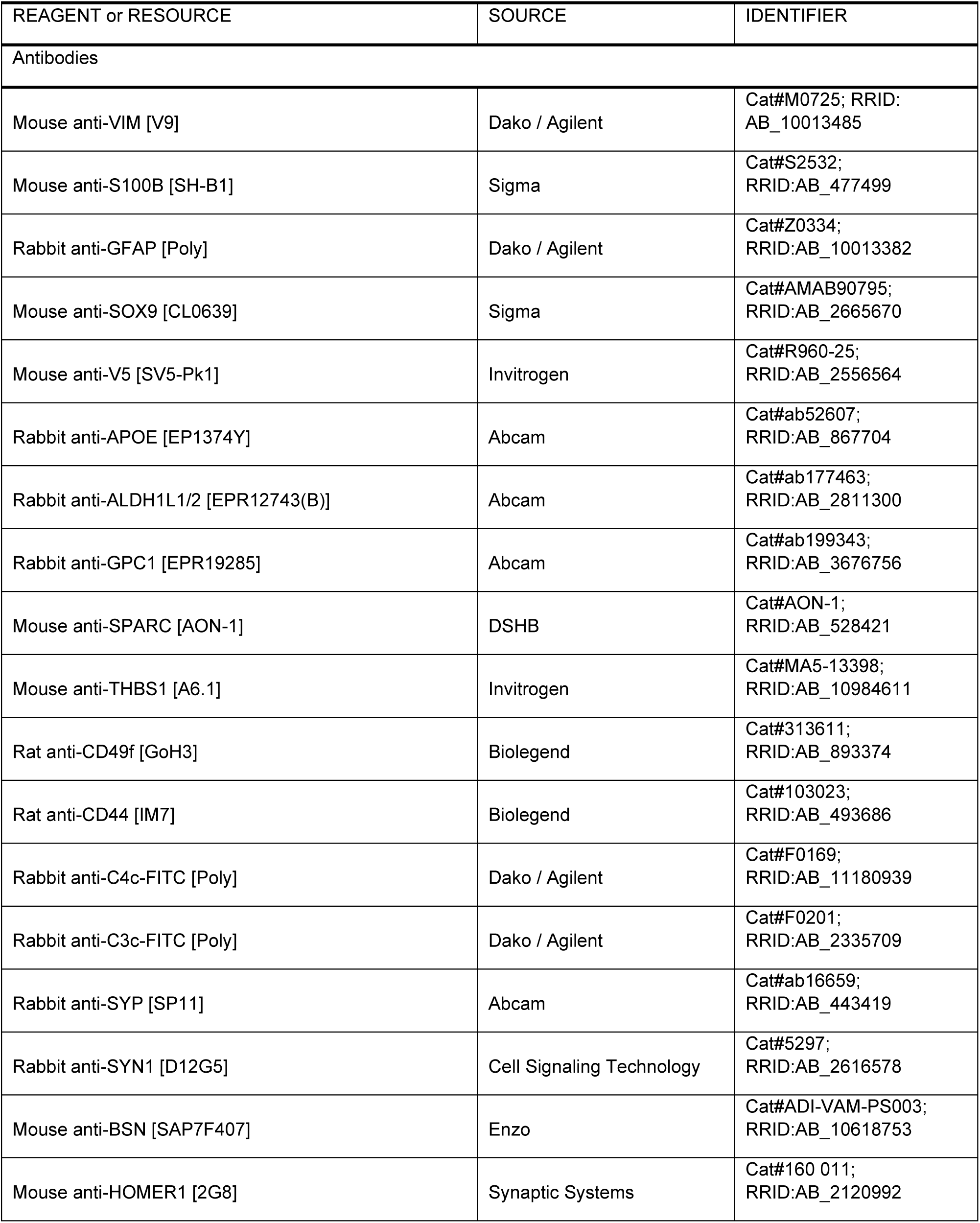
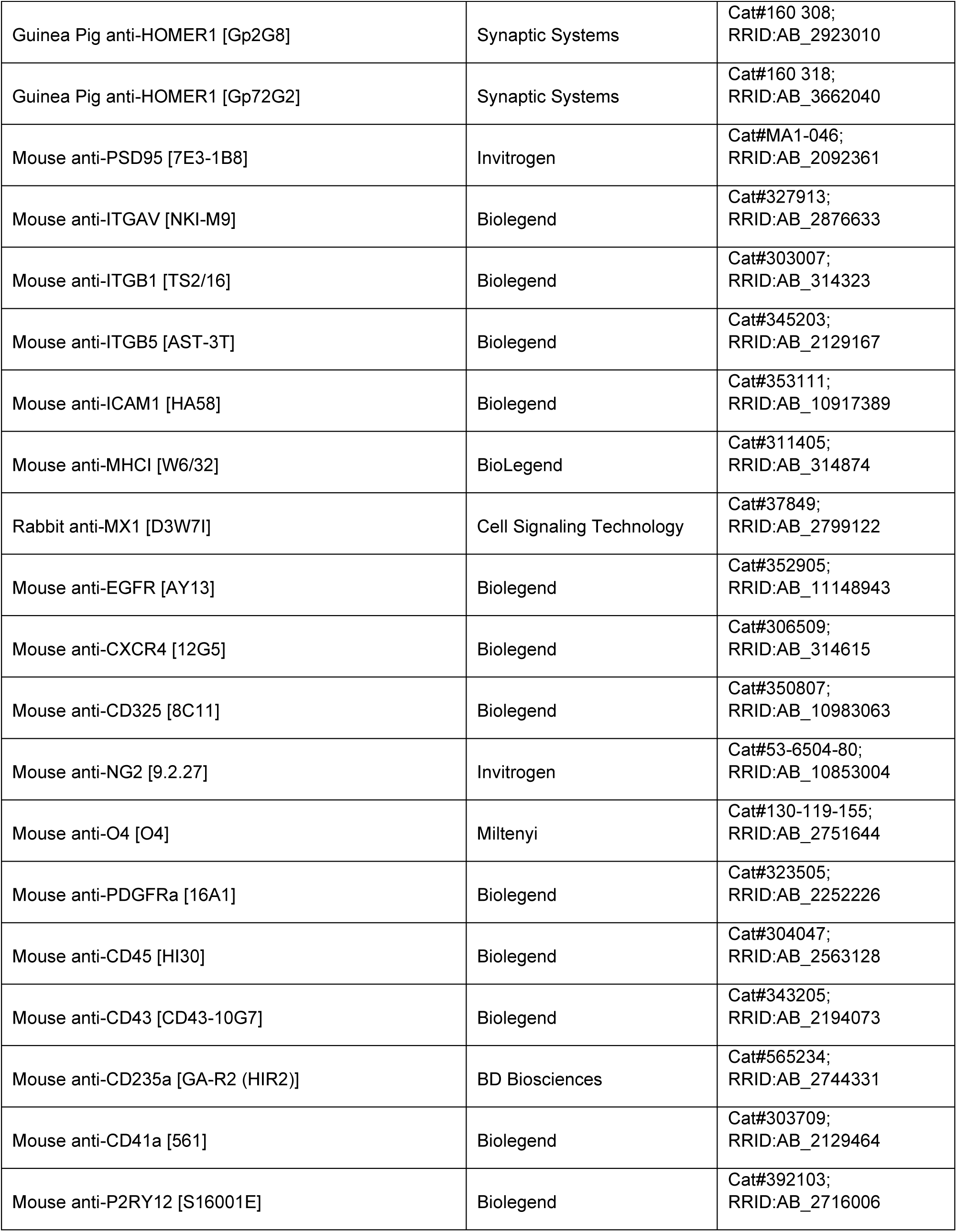
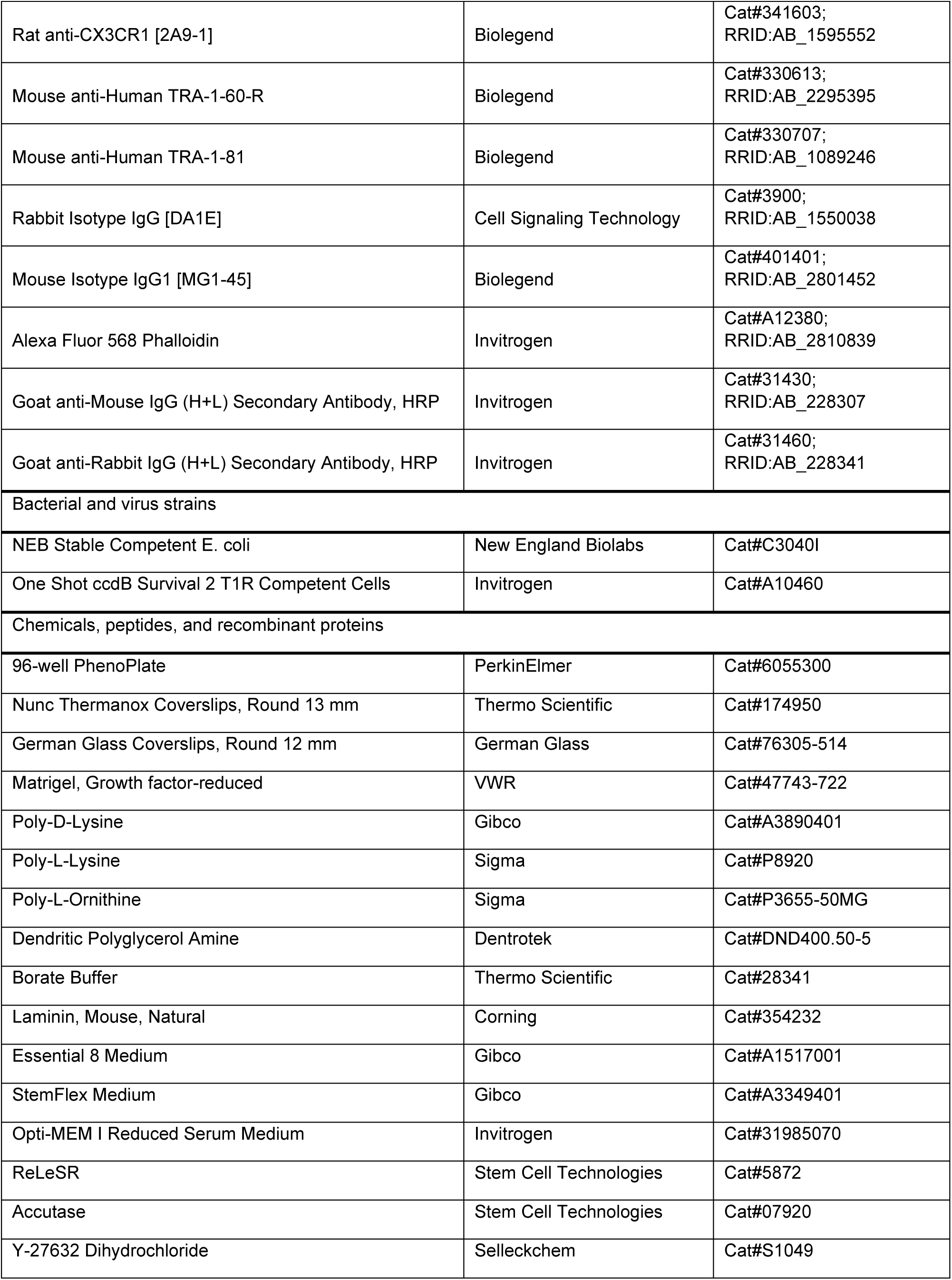
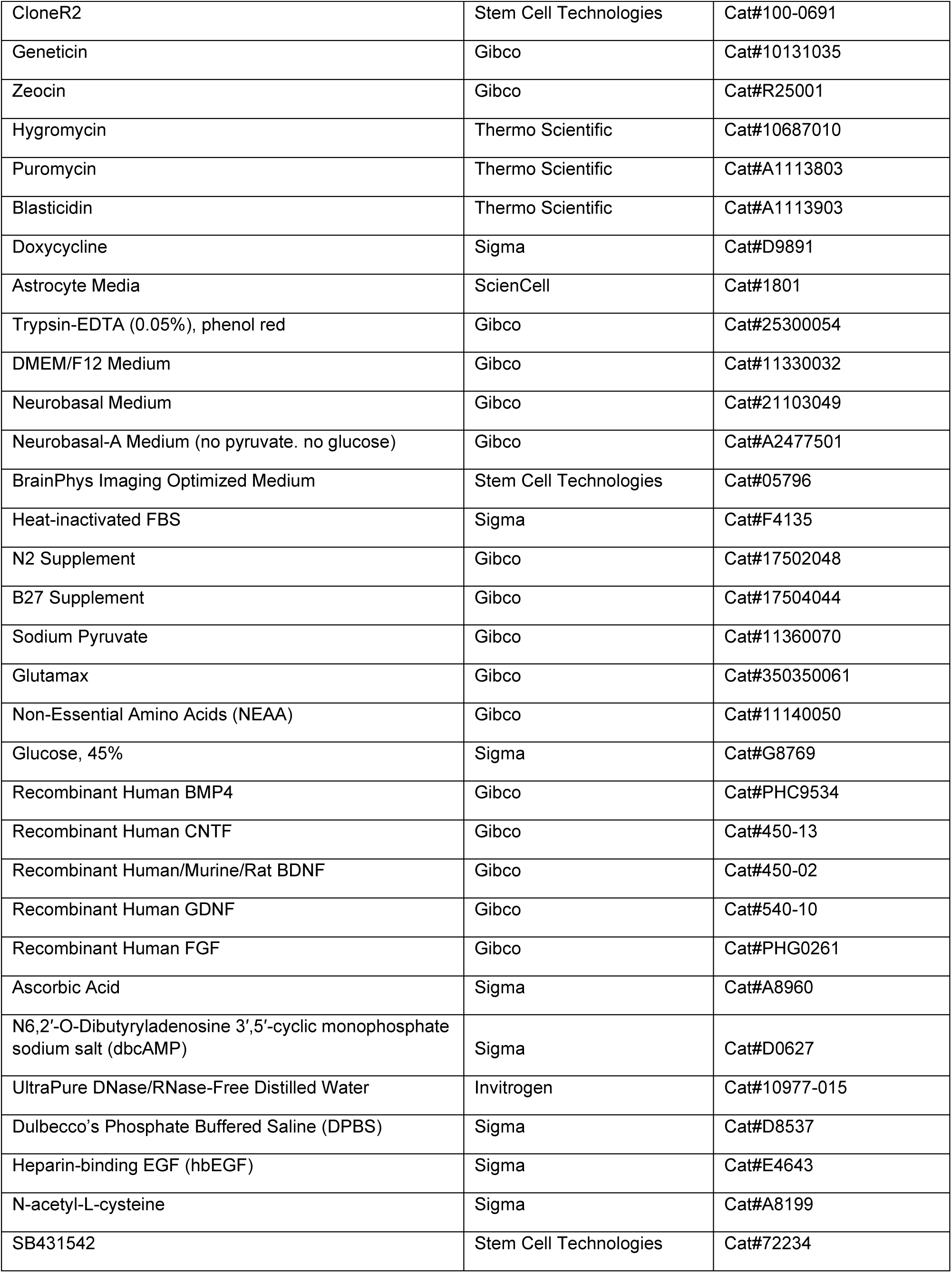
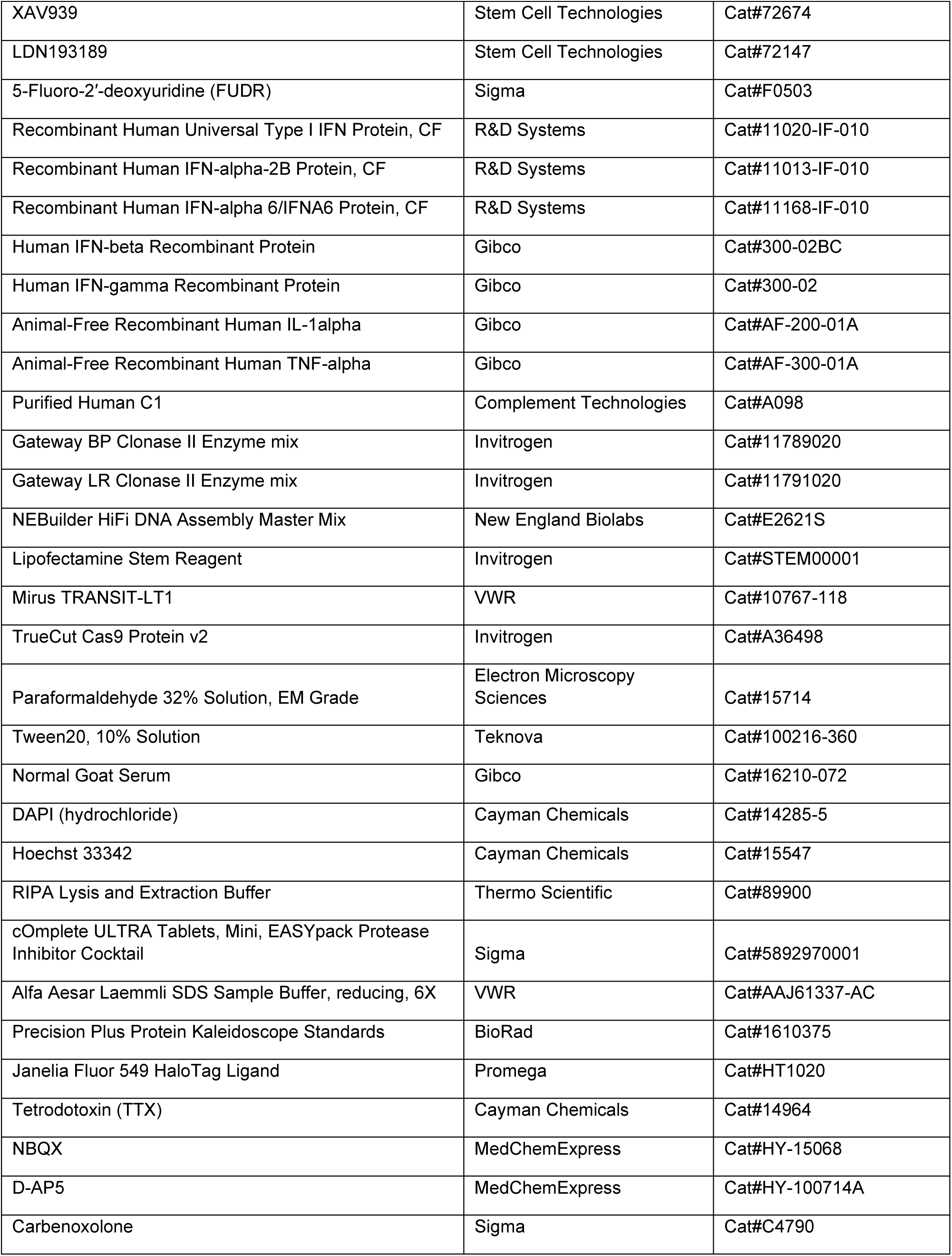
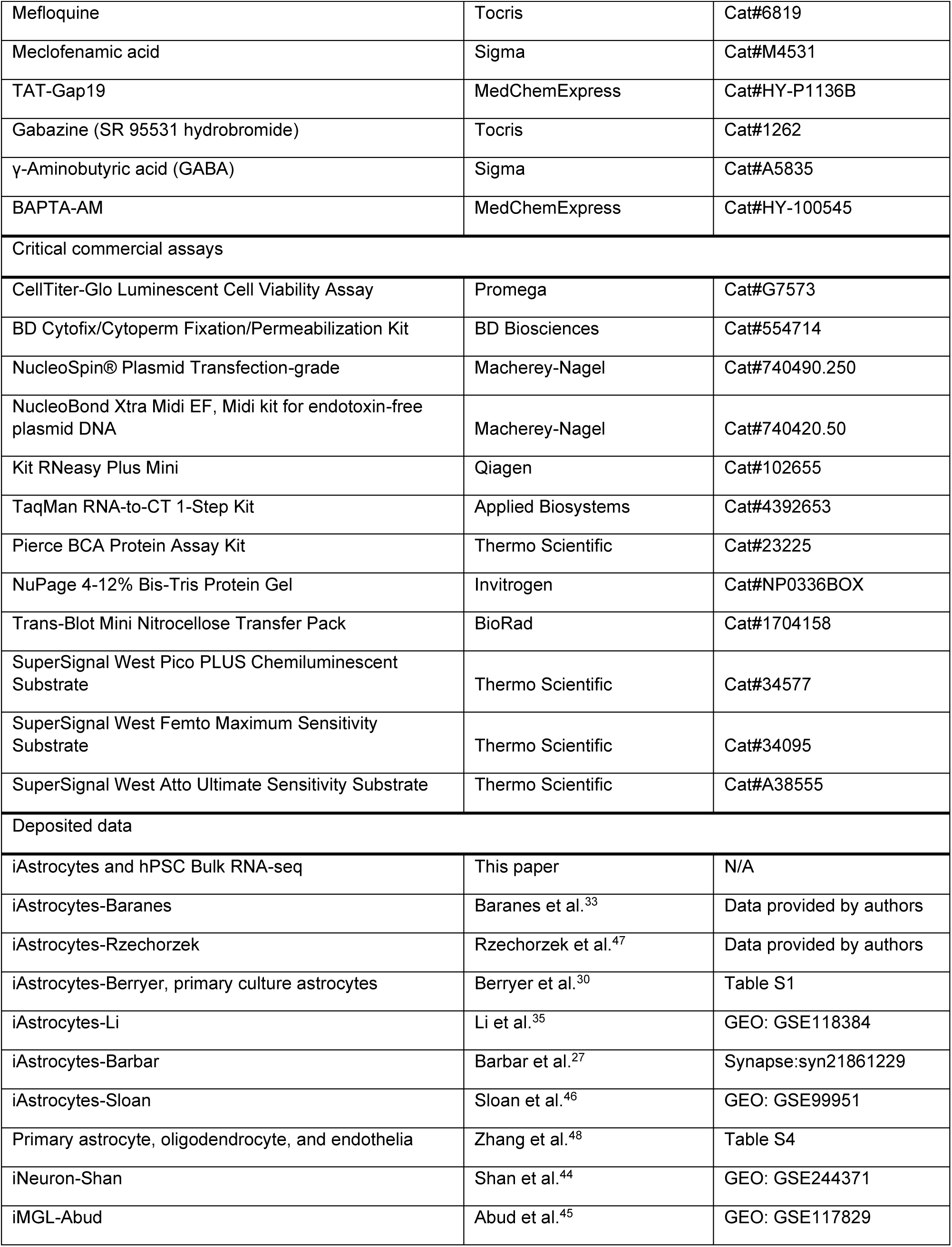
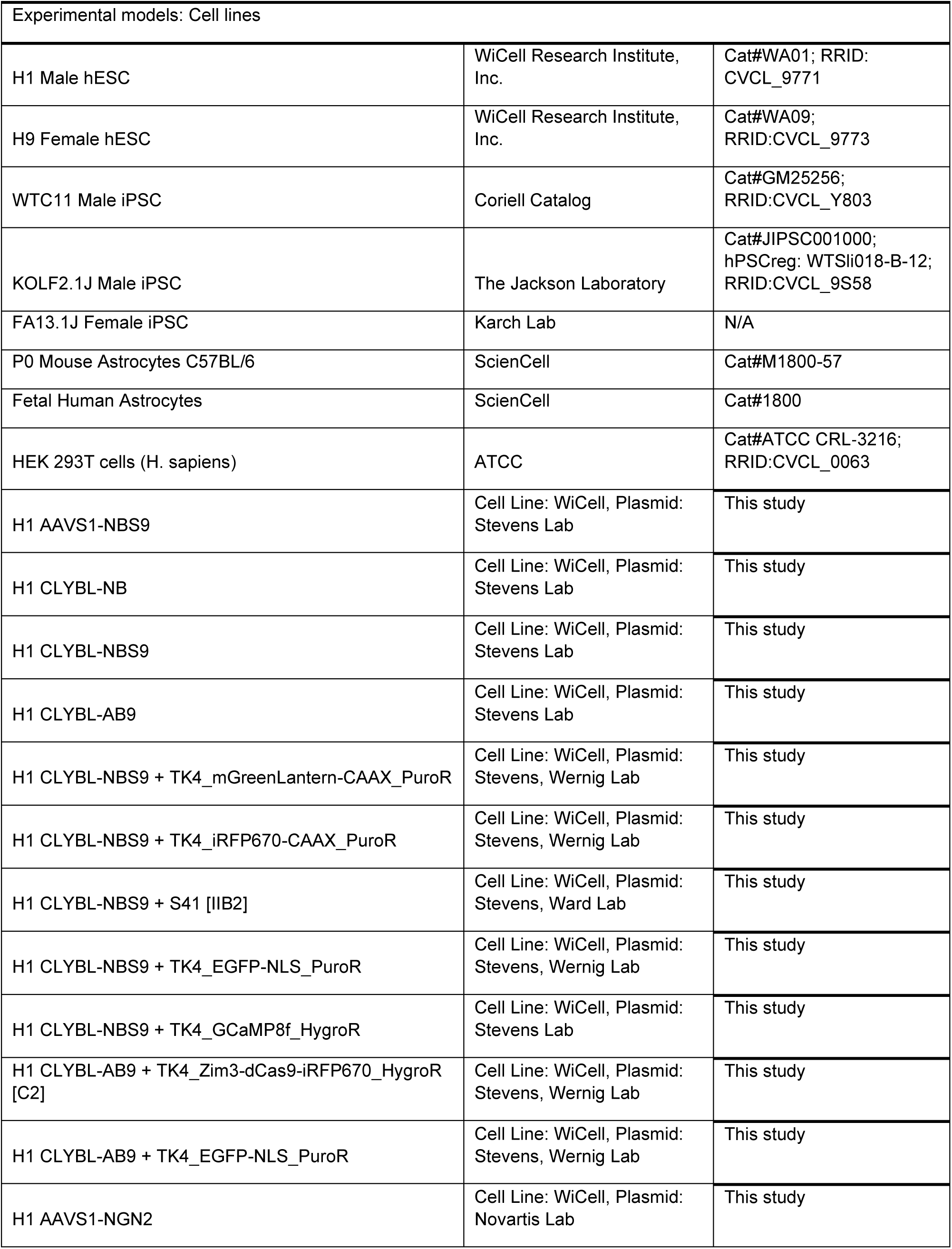
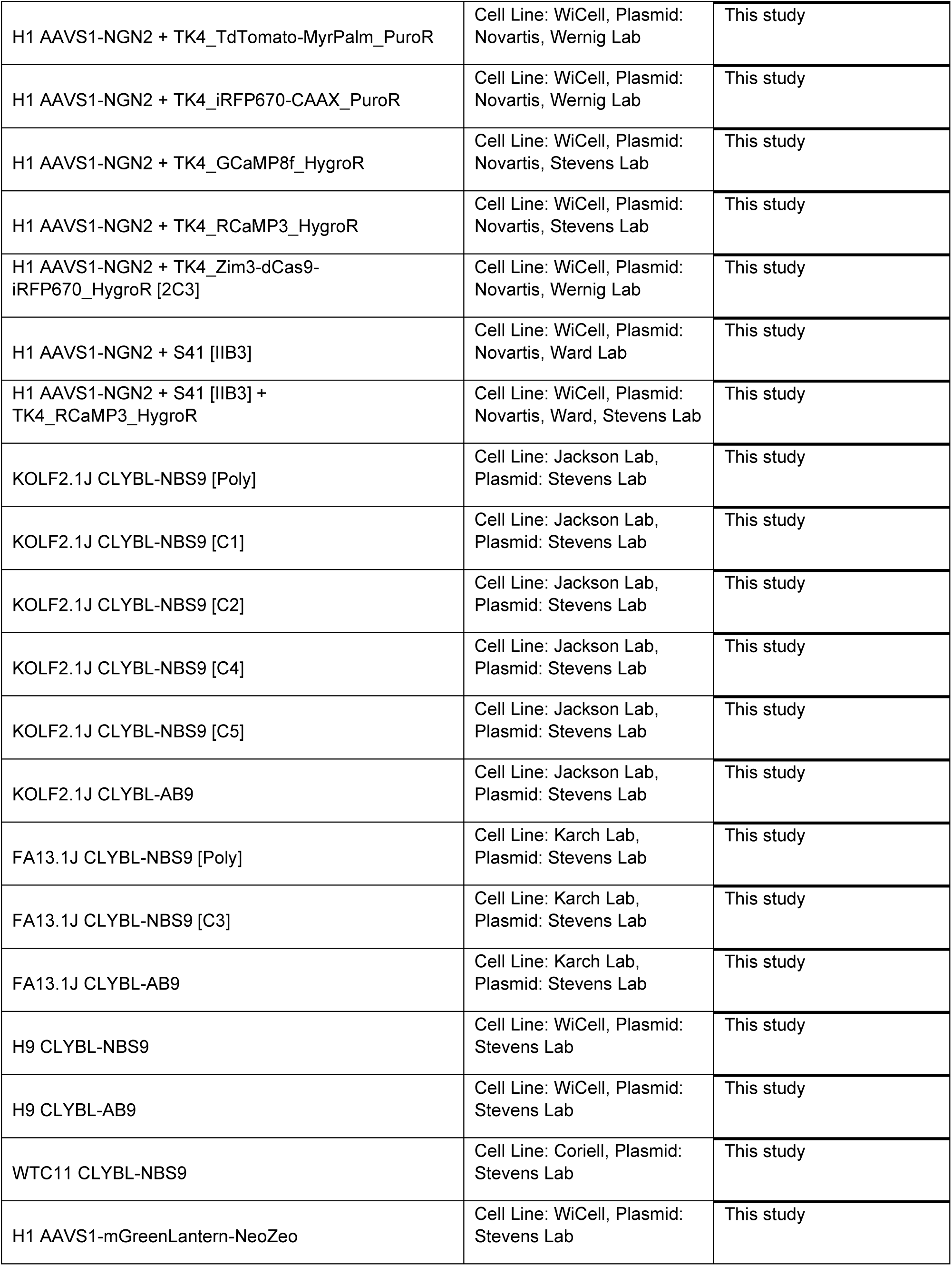
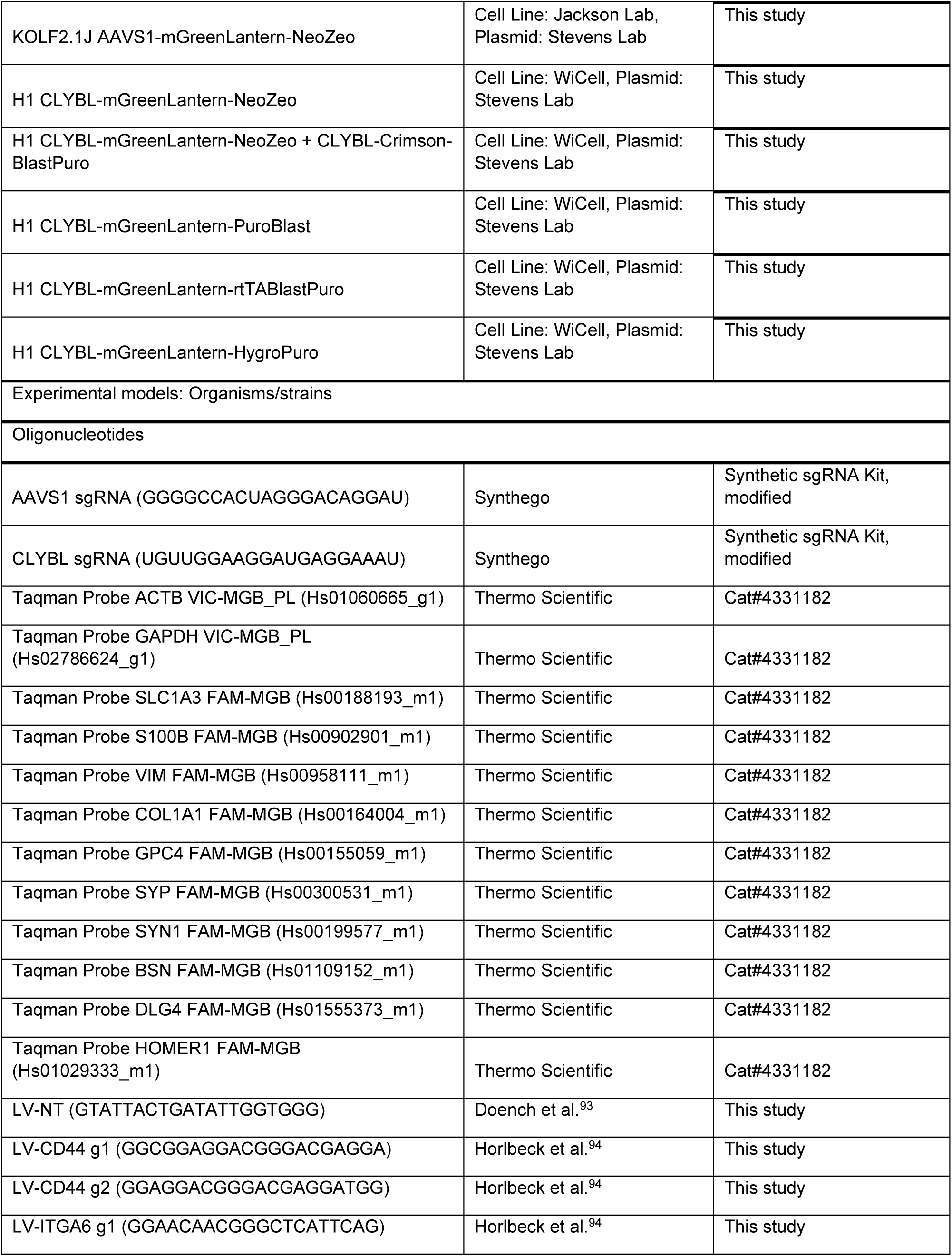
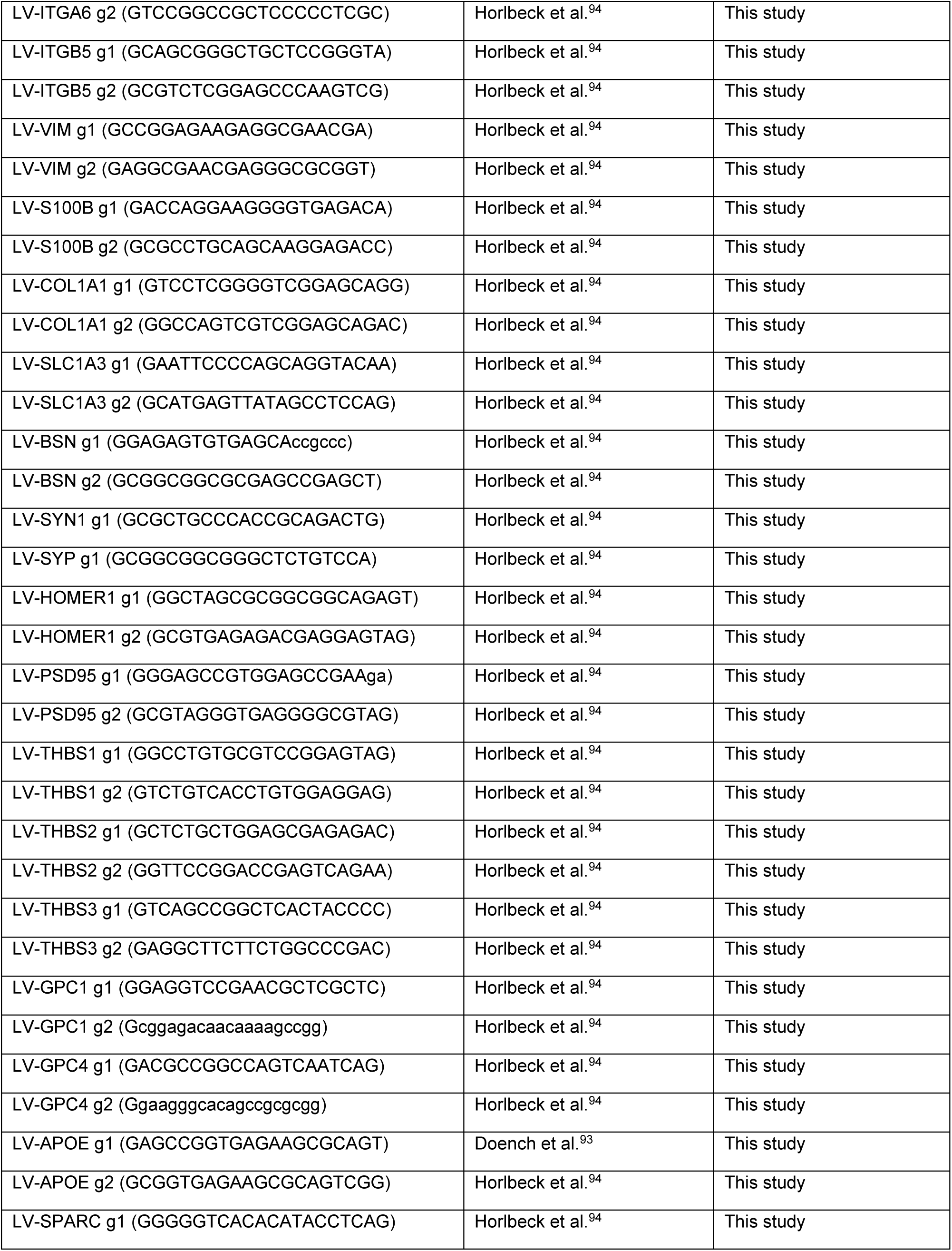
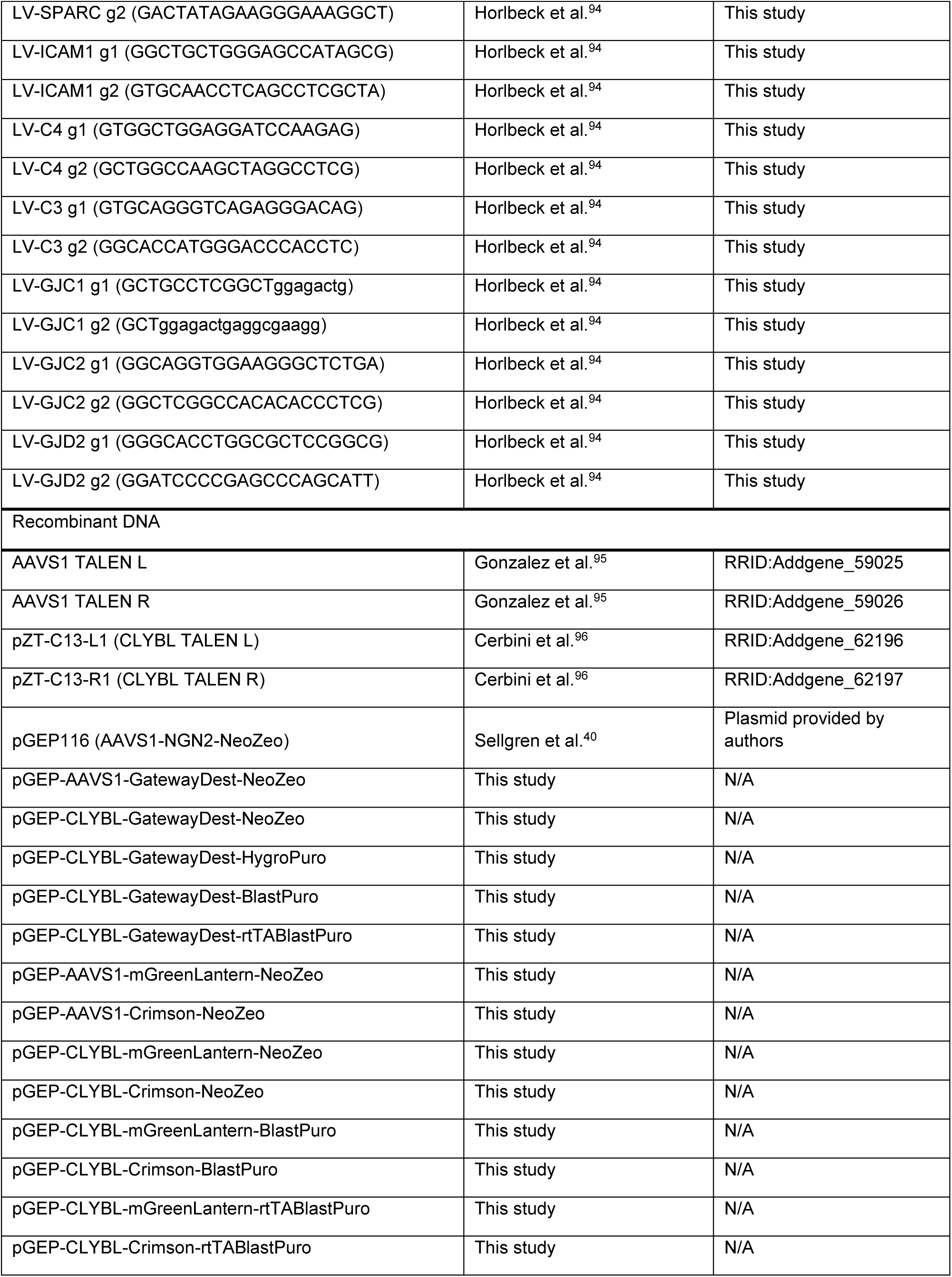
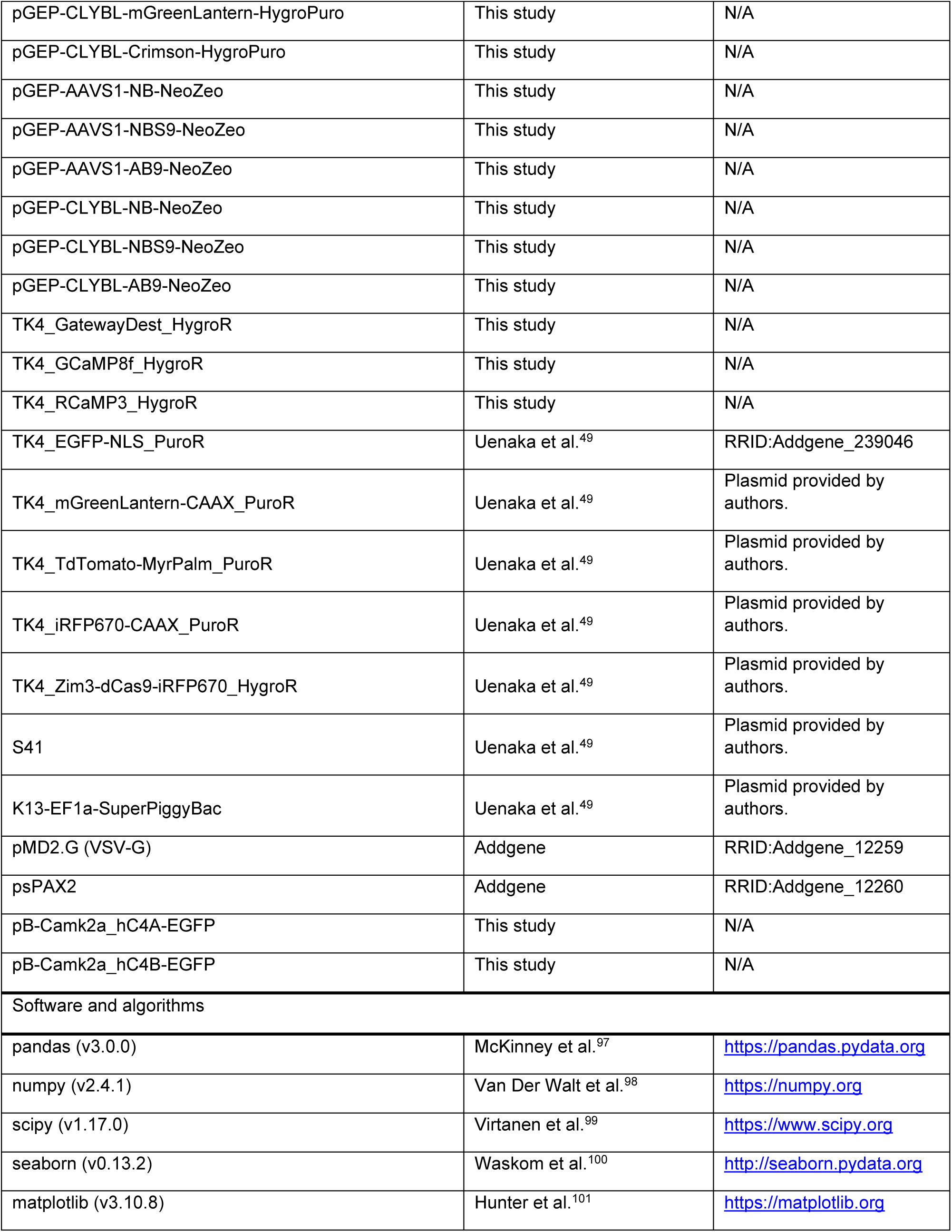
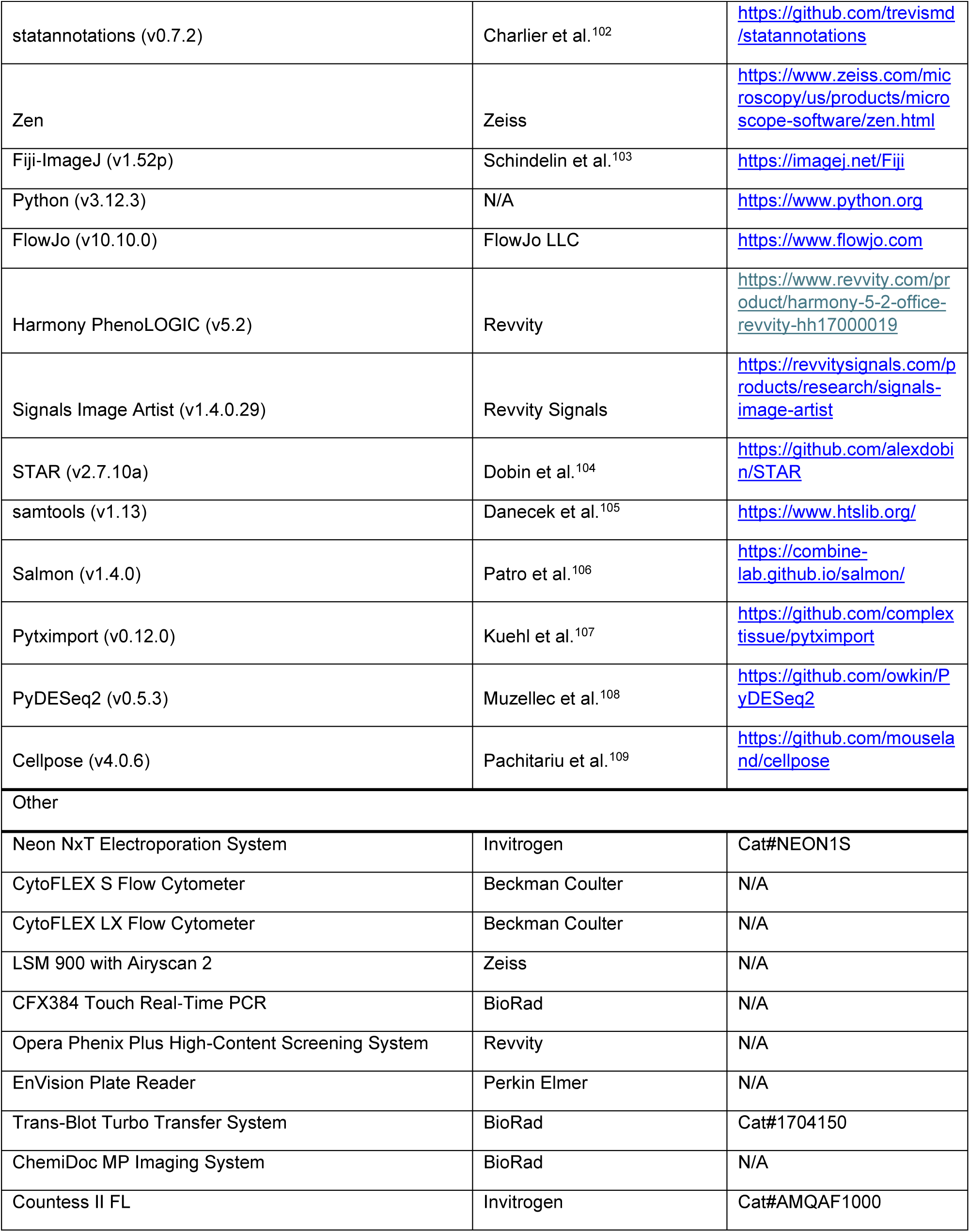

### Human pluripotent stem cell culture

All work involving stem cells underwent review and approval from the Broad Institute Office of Research Subject Protection (OSRP; NHSR-8617). Human pluripotent stem cells (hPSCs) used in this study include: H1 (male hESC) and H9 (female hESC) from WiCell^110^, WTC11/GM25256 (male iPSC) from Coriell^111^, KOLF2.1J (male iPSC) from Jackson Labs^112^, and FA13.1J (female iPSC) from Karch lab. hPSCs were cultured in Essential 8 (E8) medium on tissue culture-treated plates coated with growth factor-reduced matrigel diluted 1:100 in DMEM/F12. hPSCs were passaged with ReLeSR or accutase every 4-7 days based on colony appearance and the Rho kinase (ROCK) inhibitor Y-27632 dihydrochloride was added for 24 hours after thawing and passaging cells. hPSCs and differentiated cells were cryopreserved in CryoStor CS10.

Cytokines were prepared according to manufacturer instructions. Ascorbic acid and dbcAMP were dissolved in sterile water and filtered with a 0.22 μm filter to ensure sterility. Cells stained with trypan blue were counted automatically using Countess II. Experiments were conducted without antibiotics or antimycotics. hPSC media was tested for mycoplasma contamination.

### Safe harbor knock-in line generation

To generate engineered AAVS1 (PPP1R12C) and CLYBL safe harbor knock-in lines, hPSCs were electroporated or lipofected with “pGEP-AAVS1-*” or “pGEP-CLYBL-*” donor plasmids (see: Key Resources Table). Target genomic sites were cut using either an optimized Cas9 Ribonucleoprotein (RNP) mixture or published TALEN expression plasmids^95,96^. For electroporation, hPSCs were dissociated with accutase and 2e6 cells were mixed in Neon NxT Genome Editing (GE) Buffer with 10 μg donor plasmid and either 1.5 μg TALEN L and 1.5 μg TALEN R plasmids or 2 μg TrueCut Cas9 (Invitrogen) and 5.12 μg sgRNA targeting AAVS1 intron 1 or CLYBL intron 2 (see: **Key Resources Table**). hPSCs were electroporated using a Neon NxT system with 100 μL tips and settings of 1050V, 30 ms, 2 pulses. After electroporation, hPSCs were seeded in 6-well plates in E8 with CloneR2. For lipofection, hPSCs were dissociated with accutase and 1.5e5 cells were seeded in 12-well plates in E8 with CloneR2. Lipofection mixture was prepared with 100 μL OptiMEM, 1 μg donor plasmid, 1.25 μg TrueCut Cas9, and 3.2 μg sgRNA and mixed before adding 2.5 μL Lipofectamine Stem reagent. Lipofection mixtures were incubated for 10 minutes at RT before adding dropwise to cells. hPSC media was replaced 16-20 hours after lipofection. hPSCs were cultured in CloneR2 for 3 days to improve HDR and survival. After the first split, hPSCs were selected for genomic integration of donor plasmid using 100 μg/mL geneticin for 3 days and given 1 day without antibiotics to recover before splitting again.

To generate stable piggyBac knock-in lines, hPSCs were thawed from vials or dissociated with accutase and seeded in 12-well plates in E8 with ROCK inhibitor. Lipofection mixture was prepared with 100 μL OptiMEM, 332 ng donor plasmid, and 166 ng EF1a-SuperPiggyBac (Vector ID: K13) and mixed before adding 2.5 μL Lipofectamine Stem or TransIT-LT1 reagent. Lipofection mixtures were incubated for 10 minutes at RT before adding dropwise to cells. PiggyBac lipofections were performed in duplicate with 1x and 0.5x volume. For donor plasmids, “CRISPRi-iRFP” used vector ID: “TK4_Zim3-dCas9-iRFP670-HygroR” and “CRISPRi-Halo” used vector ID: “S41”. Two days after lipofection, hPSCs were selected with hygromycin (100 μg/mL) or puromycin (500 ng/mL) for 72 hours. hPSCs surviving with the lowest dose of transfection were split and expanded. Clonal lines were generated by splitting hPSCs with accutase and seeding 1e3 cells in a 10cm dish in StemFlex Medium with CloneR2 added for 3 days. Round colonies were scraped into accutase using a pipette tip and picking microscope. Expanded clones were analyzed for clonality by measuring fluorescent reporter signal by flow cytometry.

### Cell viability analysis

To test the functionality of NeoR, BlastR, HygroR, PuroR, and ZeoR vector elements, hPSCs with knock-in of safe harbor-resistance vectors were plated in white 96-well plates in 100 μL E8 medium with geneticin (G418), blasticidin, hygromycin, puromycin, or zeocin antibiotic selection. After 72 hours, 100 μL CellTiterGlo reagent was added to each well and plates were incubated for 15 minutes at RT with shaking. Luminescence was measured using a Perkin Elmer Envision plate reader.

### Vessel coating

Culture plates were coated with charged polymer and either growth factor-reduced matrigel or mouse laminin to promote adhesion. Polymers used include dendritic polyglycerol amine (dPGA, Dendrotek), 50 µg/mL in water; poly-L-lysine (PLL, Sigma), 100 µg/mL in water; and poly-L-ornithine (PLO, Sigma), 100 µg/mL in 100 mM borate buffer, pH 8.5 (PLO, Sigma). To coat, 50 µL polymer solution was added to each well of a 96-well plate. Plates were incubated for 1 hour at 37°C, washed thrice with water and dried in a biosafety cabinet. 50 µL per well of either matrigel (1:100 in DMEM/F12) or mouse laminin (1:100 in DMEM/F12) was added and plates were incubated at 37°C for at least one hour or until use. hPSCs, iAstrocytes (d-1 to day 7 induction, day 7 to day 21 monoculture), and induced Neurons (day -1 to day 3 induction) were plated on matrigel-coated plastic unless otherwise indicated. Primary mouse and human astrocytes were cultured on plastic. iNeuron-iAstrocyte optimized co-cultures were plated on PLO followed by matrigel unless otherwise indicated. HEK293T cells were plated on plastic for culture or PDL for immunofluorescence imaging.

### Astrocyte differentiation and culture

iAstrocytes were generated using a published differentiation media protocol^32,113^. Briefly hPSCs were split with accutase and seeded at 5.2e3 cells / cm^2^ in E8 with ROCK inhibitor (day -1). On day 0 hESCs were cultured in E8 medium with 500 ng/mL doxycycline to initiate TF expression. From days 1-6, iAstrocytes were cultured in expansion medium (EM: DMEM/F12, 10% heat-inactivated FBS, 1% N2, 1% Glutamax, 500 ng/mL doxycycline) which was gradually changed to FGF media (FGF: Neurobasal, 2% B27, 1% NEAA, 1% Glutamax, 1% heat inactivated FBS, 500 ng/mL doxycycline, with fresh 10 ng/mL BMP4, 5 ng/mL CNTF, 8 ng/mL FGF). On days 1 and 2 iAstrocytes were cultured in EM with 200 μg/mL zeocin. On days 3, 4, and 5, iAstrocytes were cultured in 3:1, 1:1, and 1:3 ratios of EM and FGF medium. On day 6, iAstrocytes were cultured in FGF medium. On day 7, iAstrocytes were dissociated with accutase and cryopreserved in CryoStor CS10. For monoculture maturation, day 7 iAstrocytes were thawed onto matrigel-coated plates in FGF medium and given a full medium change on day 8. On day 10, iAstrocytes were switched to maturation medium (MM: 50:50 Neurobasal:DMEM/F12, 1% N2, 1% Sodium Pyruvate, 1% Glutamax, 5 ng/mL heparin-binding EGF, 5 μg/mL N-acetyl-L-cysteine, 500 ng/mL doxycycline) with fresh 10 ng/mL BMP4, 10 ng/mL CNTF, 500 μg/mL dbcAMP. iAstrocytes were cultured in MM with half medium changes every 1-2 days until assay.

Primary mouse and human astrocytes were obtained from ScienCell (see: **Key Resources Table**). Primary astrocytes were cultured in Astrocyte Medium (ScienCell) and split using 0.05% Trypsin-EDTA (Gibco). Primary astrocytes were used for co-culture within 5 passages of thaw.

### Neuronal differentiation and culture

hPSCs were differentiated using a published iNeuron protocol using AAVS1 safe harbor inducible expression of NGN2 (Vector ID: pGEP116, a gift from Kathleen Worringer, Novartis)^40^ and dual-SMAD inhibition^18^. H1 AAVS1-NGN2 line was used for all iNeuron differentiations. hPSCs were split with accutase and seeded at 3.6e4 cells / cm^2^ on matrigel plates in E8 with ROCK inhibitor and 500 ng/mL doxycycline (day -1). From days 0-3, cells were cultured in N2 medium (N2: DMEM/F12, 1% N2, 0.3% Glucose, 500 ng/mL doxycycline), with 100 nM LDN193189, 2 μM XAV939, and 10 μM SB431542 added on days 0 and 1. Selection was performed with 200 μg/mL zeocin on days 1 and 2. On day 3, iNeurons were split with accutase and cryopreserved in CryoStor CS10 or used directly for co-culture.

For monocultures and co-cultures, day 3 iNeurons were seeded at a density of 4e4 iNeurons / cm^2^. For co-culture, astrocytes were seeded on the same day as iNeurons at a density ratio of 2:1 of neurons to astrocytes. Primary astrocytes were cultured for at least 3 days after thawing before re-seeding, while iAstrocytes were thawed and added immediately to co-cultures. Cultures were maintained in NBM-min medium (NBM-min: Neurobasal, 2% B27, 500 ng/mL doxycycline) or NBM-A g10 medium used Neurobasal-A (no pyruvate, no glucose) with 0.5 mM Pyruvate and 10 mM Glucose. Medium was supplemented with freshly added 10 ng/mL CNTF, 10 ng/mL GDNF, 10 ng/mL BDNF, 500 μg/mL dbcAMP, 200 μM ascorbic acid (Complete NBM-min or NBM-A g10: cNBM-min or cNBM-A g10). Co-culture day (ccD) refers to the day post seeding day 3 iNeurons and day 7 iAstrocytes or primary astrocytes. Co-cultures were given a full medium change on ccD 1 and 10 µM floxuridine (FUDR) pulse was added to suppress neuronal progenitor proliferation. Co-cultures for CRISPRi KDs were transduced with lentivirus on ccD 1 and given full medium change on ccD 2. Cultures were given half medium changes (without FUDR) every 2-3 days until time of assay.

### Production of lentivirus for CRISPRi knockdown

Protospacer sequences for CRISPRi knockdown were selected from hCRISPRi-v2.1^94^. The top 2 ranked guides per gene were cloned into a lentivirus vector with EF1α-mNeonGreen and U6-sgRNA expression (Vector ID: pJLD_LV_mNeon_sgRNA). sgRNAs for each target KD were validated by qPCR, WB, IF imaging, or flow cytometry analysis. To produce lentivirus, HEK293T cells were plated at 7.5e5 cells per well in a 12-well plate in 1 mL DMEM with 10% FBS. The next day cells were transfected with 800 ng sgRNA plasmid, 200 ng VSV-G, 500 ng psPAX2, and 4 μL TransIT-LT1 in 125 μL OptiMEM. 16-20 hours after transfection, medium was replaced with 1.5 mL OptiMEM. The next day, cell supernatant with lentivirus was collected and centrifuged in a swinging bucket at 1000g for 10 min to pellet debris. Cleared supernatant containing lentivirus was aliquoted, leaving at least 200 μL dead volume to avoid debris. Aliquots were frozen on dry ice before transfer to -80°C. Lentivirus was thawed on ice before transduction. Lentivirus batches were tested on iAstrocytes, iNeurons, or hPSCs to determine their functional titers. Knockdown experiments used 1 μL lentivirus / 100 μL medium.

### Flow cytometry analysis

Cells for flow cytometry analysis were lifted from vessels with accutase treatment for 5 min at 37°C. Cells were centrifuged at 1000g for 3 min and resuspended in 1% BSA in PBS and aliquoted for surface or internal stains. For surface stains, cells were incubated with pre-conjugated antibodies at 4°C for 15 min dark with shaking. Antibody dilutions were as follows: CD44 1:100, CD49f 1:100, ITGAV 1:100, ITGB5 1:100, ICAM1 1:100, EGFR 1:100, CXCR4 1:100, CD325 1:100, NG2 1:100, O4 1:100, PDGFRa 1:100, CD45 1:100, CD43 1:100, CD235a 1:100, CD41a 1:100, P2RY12 1:100, CX3CR1 1:100, TRA-1-60-R 1:100, TRA-1-81 1:100, Mouse Isotype 1:100. Cells were centrifuged and washed with PBS to remove excess antibodies. Cells were centrifuged and resuspended in flow buffer (0.1% BSA in PBS with DAPI 1:1000) before analysis. For internal stains, cells were centrifuged and resuspended in BD Cytofix buffer to fix for 20 min at 4°C. Cells were centrifuged, washed, and resuspended in BD Perm/Wash buffer. Cells were stained with internal antibodies for 30 min at room temperature (RT) in the dark with shaking. Cells were centrifuged, washed, and resuspended in secondary staining solution for 30 min at RT dark with shaking. Primary and secondary stains were performed in BD Perm/Wash buffer. Primary dilutions were as follows: ALDH1L1/2 1:500, VIM 1:1000, S100B 1:500, SOX9 1:500, SPARC 1:50, THBS1 1:200. Secondary antibodies were diluted 1:1000. Cells were analyzed on a CytoFLEX S or Cytoflex LX flow cytometer. Cells were gated on FSC-A/SSC-A, FSC-A/FSC-H (singlets), and V450A/SSC-A (DAPI-live cells or DAPI/33342+ fixed cells). DAPI-only conditions were included for each sample to subtract autofluorescence from antibody-stained cells resulting in delta mean fluorescent intensity (ΔMFI = MFI_stain+dapi_ - MFI_dapi_).

### Western blot analysis

For western blots, cells were lysed in ice-cold RIPA with protease inhibitors. Lysates were cleared by centrifugation at 20,000g for 15 min at 4°C. Protein concentration was quantified by BCA. 10-20 μg aliquots of sample were boiled in reducing SDS buffer for 10 min at 90°C. Protein lysates and IP eluates were separated on NuPAGE 4-12% Bis-Tris gels at 90-130V. Proteins were transferred to Trans-Blot nitrocellulose membranes using a Trans-Blot Turbo Transfer System. Membranes were blocked overnight at 4°C with 5% milk in TBST. Blocking buffer was replaced with primary antibody in 2.5% milk in TBST overnight at 4°C. Primary antibody dilutions were as follows: GPC1 1:1000, THBS1 1:1000, SPARC 1:500, APOE 1:1000, B-actin-HRP 1:10,000. Membranes were washed 3 x 5 min in TBST and stained with secondary antibody (Invitrogen: Goat anti-* Ig* (H+L) HRP Secondary Antibody) at 1:10,000 in 2.5% milk for 1 hour at RT. Membranes were washed 4 x 5 min in TBST and imaged with chemiluminescent substrate using a ChemiDoc MP Imaging System.

### Immunofluorescence staining and imaging

Cells for Immunofluorescence staining and imaging were cultured in 96-well PhenoPlate vessels (see: **Vessel coating**). At time of assay, medium was replaced with 4% paraformaldehyde (PFA) solution diluted in PBS from 32% stock solution to fix cells. Cells were fixed for 20 minutes at RT in the dark and then washed with PBS to remove residual PFA. Cells were blocked and permeabilized in 10% Normal Goat Serum (NGS) + 0.1% Triton for 1 hour at RT or overnight at 4°C. Primary antibodies were stained in 5% NGS + 0.1% Triton overnight at 4°C. Primary antibody dilutions were as follows: VIM 1:1000, S100B 1:500, GFAP 1:1000, SOX9 1:500, V5 1:1000, APOE 1:500, ALDH1L1/2 1:500, SPARC 1:50, THBS1 1:200, C4 1:1000, C3 1:1000, SYP 1:200, SYN1 1:1000, BSN 1:1000, HOMER1 [2G8] 1:500, HOMER1 [Gp2G8] 1:500, HOMER1 [Gp72G2] 1:500, PSD-95 1:500, MHC-I 1:100, MX1 1:250, Rabbit Isotype 1:250, Mouse Isotype 1:500, Phalloidin 1:500. Cells were washed 3x with PBS and then stained with secondary antibodies (Invitrogen: Goat anti-* Ig* (H+L) Cross-Adsorbed Secondary Antibody) and DAPI or Hoechst 33342 diluted 1:1000 in 5% NGS + 0.1% Triton for 1 hour at RT. Cells were washed 3x with PBS and stored dark until assay.

Live cells and synapse stains were imaged using Opera Phenix High Content Imaging System. Phenix images were collected using a 10x air- or 63x water-immersion objective and 405, 488, 568/594, and 647 fluorescent channels. EGFP-NLS nuclei were measured using SIMA “Find Nuclei” with parameter “Method=A”. mGL-CAAX and TdT-MyrPalm area were measured using SIMA “Find Image Region” with an absolute threshold set based on no cell controls. Synapse area overlapping with mNeonGreen was measured using SIMA “Find Image Region” with an absolute threshold set based on isotype antibody and untransduced controls. CRISPRi KD and reactive astrocyte assays were imaged using Zeiss LSM900 with AiryScan 4Y super-resolution imaging module. Zeiss images were collected using 10x air-, 20x air-, and 63x oil-immersion objectives. Intensity and percent area were quantified using ImageJ with an absolute threshold based on isotype antibody for KD assays or uninduced control for reactive assays.

### qPCR and RNA-seq analysis

Samples for RNA analysis were lysed in Buffer RLT with 40 mM DTT. RNA samples were isolated with AllPrep DNA/RNA or RNeasy Plus Micro kits. Samples for qPCR were prepared with RNA-to-Ct kit and RNA targets were detected using TaqMan probes listed in **Key Resources Table**. CFX384 Touch Real-Time PCR was used for qPCR amplification. qPCR Cq values were exported from the Bio-Rad CFX Manager and processed with pandas. Cq values < 15 or > 40 were discarded and median Cq values were calculated from 2 technical replicates of each sample-target combination. Fold changes were calculated as 2^−ΔΔ*Cq*^ for quantification cycle number relative to ACTB in each sample.

Bulk RNA-seq was performed using Novogene commercial sequencing service. Directional mRNA libraries were prepared using ultra-low input total RNA directional library preparation. Sequencing was performed using NovaSeq X Plus Series (PE150). In accordance with sharing guidelines by Jackson Laboratory, fastq data files were processed before alignment and deposition to remove Y chromosome reads. Reads were aligned to the human genome with STAR^104^ using Ensembl genome files: ‘Homo_sapiens.GRCh38.dna.primary_assembly.fa.gz’ and ‘Homo_sapiens.GRCh38.115.gtf.gz’. Output BAM files were processed with samtools^105^ to identify Y chromosome reads. Fastq files were filtered to remove Y chromosome reads using custom shell scripts. Salmon^106^ was used to quantify transcript abundances from filtered fastq files. Salmon index was built using ‘gencode.v48.transcripts.fa.gz’. pytximport^107,114^ was used to process salmon output ‘quant.sf’ files into RNA counts matrix (**Table S1**). RNA-seq data for other primary and induced cell types were downloaded from public repositories (see **Key Resources Table**). Fastq files for Barbar_* samples were aligned using salmon and processed into RNA counts matrix using pytximport. Data sets were combined using pandas.

RNA counts were normalized to 1e6 per sample (TPM: transcripts per million) and log10-transformed (log10(x+1)). Differential gene expression was calculated using PyDESeq2^108^. Genes were considered differentially expressed if baseMean > 10, abs(log2FoldChange) > 2, and padj < 1e-3. Heatmaps were generated using seaborn^100^. For principal component analysis (PCA), datasets were combined using ‘pandas.DataFrame.concat(join=”outer”)’ and filtered to only include genes with >1e2 TPM across at least 3 samples for “This Study” or 6 samples for “Other Studies”. Data was normalized using ‘sklearn.preprocessing.StandardScaler’ and PCA was performed using ‘sklearn.decomposition.PCA’ with the parameter ‘n_components=2’^99^. Enrichment scores were calculated by taking the median log_10_(TPM) of samples within a group minus the largest median log_10_(TPM) of samples from each out-group. In-group, out-group combinations used to calculate enrichment scores were: {‘iAstro’: [’iNeuro’, ’iMGL’, ’hPSC’], ‘pAstro’: [’iNeuro’, ’iMGL’, ’pEndo’, ’pOligo’, ’hPSC’], ‘iNeuro’: [’iAstro’, ’pAstro’, ’iMGL’, ’hPSC’], ‘iMGL’: [’iAstro’, ’pAstro’, ’iNeuro’, ’hPSC’], ‘pEndo’: [’iAstro’, ’pAstro’, ’iNeuro’, ’iMGL’, ’pOligo’, ’hPSC’], ‘pOligo’: [’iAstro’, ’pAstro’, ’iNeuro’, ’iMGL’, ’pEndo’, ’hPSC’], ‘hPSC’: [’iAstro’, ’pAstro’, ’iNeuro’, ’iMGL’, ’pEndo’, ’pOligo’]}.

### Reactive astrocyte induction assays

Reactive astrocyte induction assays were performed on H1 AAVS1-NBS9 and H1 CLYBL-AB9 CRISPRi-iRFP [C2] hPSC lines (see **Table S3**). iAstrocytes were cultured in EM, FGF, and MM media which include 1% serum until day 10 but are fully serum-free for 11 days before assay. iAstrocytes were induced until day 20 and then treated for 24 hours with individual interferons or combinations of TNFα, IL1α, and C1q (TIC). TIC treatment was based on a method to induce neurotoxic astrocytes^12^ and used 3 ng/mL human recombinant IL1α, 30 ng/mL human recombinant TNFα, and 400 ng/mL human purified C1. Interferon treatment used 50 μg/mL IFNαU, IFNα2b, IFNα6, IFNβ, and IFNγ or 32 ng/mL - 50 μg/mL for IFNβ and IFNγ titration experiments.

### Live culture imaging

hPSCs with PiggyBac expression of mGreenLantern-CAAX^115^, TdTomato-MyrPalm^116^, iRFP670-CAAX^117^, GCaMP8f (subcloned from Addgene: 162376)^118^, or RCaMP3 (synthesized as gBlock)^119^ were differentiated into iNeurons or iAstrocytes in co-culture. Co-cultures were maintained and imaged in NBM, NBM-A g10, or BrainPhys Imaging medium (see **Table S3**). Co-cultures were imaged using Opera Phenix. Chamber was pre-equilibrated to 37°C and 5% CO_2_ before imaging. Images were taken using a 10x objective and 405, 488, 568, and 647 fluorescent channels. Images were analyzed using SIMA. Cell regions were selected using an absolute intensity threshold, divided into objects using find cells method “P” or find nuclei method “A”, and filtered for objects > 20 μm^2^ size to remove debris. Metrics for each well were averaged over a 3×3 field of view (FOV) grid.

### Recording neuronal calcium activity

hPSCs with PiggyBac expression of GCaMP8f (green-shifted) or RCaMP3 (red-shifted) calcium indicators were differentiated into iNeurons and co-cultured with an array of astrocyte models. iNeurons were co-cultured in Neurobasal medium variants (NBM, NBM-A g10) with B27 and Dox and freshly added supplements (CTNF, BDNF, GDNF, dbcAMP, and ascorbic acid). For medium switching experiments, cells were given a full medium change on day of imaging to BrainPhys Imaging Medium with freshly added supplements. For pharmacological experiments, iNeurons were treated with molecules for 1 hour before imaging. High throughput fluorescence imaging of calcium indicators was performed using Opera Phenix. Chamber was pre-equilibrated to 37°C and 5% CO_2_ before recording. Videos were recorded using a 10x objective and 488 or 568 fluorescent channels, 320 ms exposure time, and 3 frames per second. For 2-color live fluorescence imaging, channels were imaged sequentially every frame with exposure times: 488: 100 ms 568: 160 ms. Recordings were exported to TIFF image stacks using Harmony software.

### Analyzing neuronal calcium activity

Semi-automated live fluorescence imaging analysis of calcium transients was performed on TIFF stacks using custom Python scripts [Gonzalez-Ramos et al. (manuscript in preparation)]. In brief, neuronal somas were automatically detected as regions of interest (ROIs) using CellPose-SAM^109,120^. From each ROI, fluorescence intensity was extracted across the time series. Resulted datasheets containing raw fluorescence calcium traces were used as inputs for all algorithms. Sliding window normalization was applied to generate ΔF/F0 profiles for each ROI. Prior to event detection, traces were smoothed using a Savitzky-Golay filter^121^. Calcium transients were then detected using an algorithm based on height, prominence, and minimum inter-peak distance parameters. The initial 5 seconds of each recording were excluded from analysis to avoid artifacts from photobleaching following laser activation. ROIs with fewer than 2 events per minute were excluded from subsequent analysis. Detected events were displayed as raster plots, with mean frequency and inter-event interval (IEI) calculated across all ROIs for each recording and used for comparison between conditions. Global Network Activity (GNA) was quantified using a sliding window and determined as the number of ROIs exhibiting at least one calcium transient within a time window or bin of 1.5 seconds, using a step size of 0.3 seconds. Cumulative plots for active ROIs at a given time window were represented as a histogram and shown along with a GNA sliding window trace above each raster plot, along time of recording. Then, the fraction of active ROIs (A) was calculated as the number of ROIs (≥ 1 event within the window) divided by the total number of ROIs (A = active ROIs / total ROIs). Synchronous network events (collective events) were identified as peaks in the GNA trace where A ≥ 0.30, meaning ≥30% of ROIs were co-active. The A fraction of synchronous events were used to compare network synchronicity between conditions, as mean A * total number of collective events. Videos with timestamps were exported to ‘.avi’ files using ImageJ.

## Data and Code Availability

All sequencing data have been deposited in Gene Expression Omnibus and are available under the accession: GSE334385 (https://www.ncbi.nlm.nih.gov/geo/query/acc.cgi?acc=GSE334385). Sources for other code used in this study are indicated in the **Key Resources Table**.

## References

1. Chung, W.-S., Allen, N. J. & Eroglu, C. Astrocytes Control Synapse Formation, Function, and Elimination. Cold Spring Harb. Perspect. Biol. 7, a020370 (2015).

2. Allen, N. J. Astrocyte regulation of synaptic behavior. Annu. Rev. Cell Dev. Biol. 30, 439–463 (2014).

3. De Strooper, B. & Karran, E. The Cellular Phase of Alzheimer’s Disease. Cell 164, 603–15 (2016).

4. Green, G. S. et al. Cellular communities reveal trajectories of brain ageing and Alzheimer’s disease. Nature 633, 634–645 (2024).

5. Habib, N. et al. Disease-associated astrocytes in Alzheimer’s disease and aging. Nat. Neurosci. 23, 701–706 (2020).

6. Guttenplan, K. A. et al. Knockout of reactive astrocyte activating factors slows disease progression in an ALS mouse model. Nat. Commun. 11, 1–9 (2020).

7. Ling, E. et al. A concerted neuron–astrocyte program declines in ageing and schizophrenia. Nature 627, 604–611 (2024).

8. Labarta-Bajo, L. & Allen, N. J. Astrocytes in aging. Neuron 113, 109–126 (2025).

9. Boisvert, M. M., Erikson, G. A., Shokhirev, M. N. & Allen, N. J. The Aging Astrocyte Transcriptome from Multiple Regions of the Mouse Brain. Cell Rep. 22, 269–285 (2018).

10. Leng, K. et al. CRISPRi screens in human iPSC-derived astrocytes elucidate regulators of distinct inflammatory reactive states. Nat. Neurosci. 25, 1528–1542 (2022).

11. Sadick, J. S. et al. Astrocytes and oligodendrocytes undergo subtype-specific transcriptional changes in Alzheimer’s disease. Neuron 110, 1788–1805.e10 (2022).

12. Liddelow, S. A. et al. Neurotoxic reactive astrocytes are induced by activated microglia. Nature 541, 481–487 (2017).

13. Serrano-Pozo, A. et al. Astrocyte transcriptomic changes along the spatiotemporal progression of Alzheimer’s disease. Nat. Neurosci. 27, 2384–2400 (2024).

14. Clarke, L. E. et al. Normal aging induces A1-like astrocyte reactivity. Proc. Natl. Acad. Sci. U. S. A. 115, E1896–E1905 (2018).

15. Sekar, A. et al. Schizophrenia risk from complex variation of complement component 4. Nature 530, 177–183 (2016).

16. Naj, A. C. et al. Common variants at MS4A4/MS4A6E, CD2AP, CD33 and EPHA1 are associated with late-onset Alzheimer’s disease. Nat. Genet. 43, 436–441 (2011).

17. Johnson, M. B. & Stevens, B. Pruning hypothesis comes of age. Nature 554, 438–439 (2018).

18. Nehme, R. et al. Combining NGN2 Programming with Developmental Patterning Generates Human Excitatory Neurons with NMDAR-Mediated Synaptic Transmission. Cell Rep. 23, 2509–2523 (2018).

19. Lu, C. et al. Overexpression of NEUROG2 and NEUROG1 in human embryonic stem cells produces a network of excitatory and inhibitory neurons. FASEB J. 33, 5287–5299 (2019).

20. Pietiläinen, O. et al. Astrocytic cell adhesion genes linked to schizophrenia correlate with synaptic programs in neurons. Cell Rep. 42, 111988 (2023).

21. Pembroke, W. G., Hartl, C. L. & Geschwind, D. H. Evolutionary conservation and divergence of the human brain transcriptome. Genome Biol. 22, 52 (2021).

22. Oberheim, N. A., Wang, X., Goldman, S. & Nedergaard, M. Astrocytic complexity distinguishes the human brain. Trends Neurosci. 29, 547–553 (2006).

23. Oberheim, N. A. et al. Uniquely hominid features of adult human astrocytes. J. Neurosci. 29, 3276–3287 (2009).

24. Rapino, F. et al. Small-molecule screen reveals pathways that regulate C4 secretion in stem cell-derived astrocytes. Stem Cell Reports 11, 363–379 (2022).

25. Guttikonda, S. R. et al. Fully defined human pluripotent stem cell-derived microglia and tri-culture system model C3 production in Alzheimer’s disease. Nat. Neurosci. 24, 343–354 (2021).

26. Krencik, R. & Zhang, S. C. Directed differentiation of functional astroglial subtypes from human pluripotent stem cells. Nat. Protoc. 6, 1710–1717 (2011).

27. Barbar, L. et al. CD49f Is a Novel Marker of Functional and Reactive Human iPSC-Derived Astrocytes. Neuron 107, 436–453.e12 (2020).

28. Jovanovic, V. M., et al. A defined roadmap of radial glia and astrocyte differentiation from human pluripotent stem cells. Stem Cell Reports 18, (2023).

29. TCW, J., et al. An Efficient Platform for Astrocyte Differentiation from Human Induced Pluripotent Stem Cells. Stem Cell Reports 9, 600–614 (2017).

30. Berryer, M. H. et al. Robust induction of functional astrocytes using NGN2 expression in human pluripotent stem cells. iScience 26, 106995 (2023).

31. Caiazzo, M. et al. Direct conversion of fibroblasts into functional astrocytes by defined transcription factors. Stem Cell Reports 4, 25–36 (2015).

32. Canals, I. et al. Rapid and efficient induction of functional astrocytes from human pluripotent stem cells. Nat. Methods 15, 693–696 (2018).

33. Baranes, K. et al. Transcription factor combinations that define human astrocyte identity encode significant variation of maturity and function. Glia 71, 1870–1889 (2023).

34. Lam, I. et al. Rapid iPSC inclusionopathy models shed light on formation, consequence, and molecular subtype of α-synuclein inclusions. Neuron 112, 2886–2909.e16 (2024).

35. Li, X. et al. Fast Generation of Functional Subtype Astrocytes from Human Pluripotent Stem Cells. Stem Cell Reports 11, 998–1008 (2018).

36. Tchieu, J. et al. NFIA is a gliogenic switch enabling rapid derivation of functional human astrocytes from pluripotent stem cells. Nat. Biotechnol. 37, 267–275 (2019).

37. Cvetkovic, C. et al. Assessing Gq-GPCR–induced human astrocyte reactivity using bioengineered neural organoids. J. Cell Biol. 221, (2022).

38. Yi, R. et al. A single-cell transcriptomic dataset of pluripotent stem cell-derived astrocytes via NFIB/SOX9 overexpression. Sci. Data 11, 1–11 (2024).

39. Lee, H. et al. Contributions of Genetic Variation in Astrocytes to Cell and Molecular Mechanisms of Risk and Resilience to Late-Onset Alzheimer’s Disease. Glia 73, 1166–1187 (2025).

40. Sellgren, C. M. et al. Increased synapse elimination by microglia in schizophrenia patient-derived models of synaptic pruning. Nat. Neurosci. 22, 374–385 (2019).

41. Alerasool, N., Segal, D., Lee, H. & Taipale, M. An efficient KRAB domain for CRISPRi applications in human cells. Nat. Methods 17, 1093–1096 (2020).

42. Stevens, B. et al. The Classical Complement Cascade Mediates CNS Synapse Elimination. Cell 131, 1164–1178 (2007).

43. Limone, F. et al. Efficient generation of lower induced motor neurons by coupling Ngn2 expression with developmental cues. Cell Rep. 42, 111896 (2023).

44. Shan, X. et al. Fully defined NGN2 neuron protocol reveals diverse signatures of neuronal maturation. Cell Reports Methods 4, 100858 (2024).

45. Abud, E. M. et al. iPSC-Derived Human Microglia-like Cells to Study Neurological Diseases. Neuron 94, 278–293.e9 (2017).

46. Sloan, S. A. et al. Human Astrocyte Maturation Captured in 3D Cerebral Cortical Spheroids Derived from Pluripotent Stem Cells. Neuron 95, 779–790.e6 (2017).

47. Rzechorzek, N. M., et al. Circadian clocks in human cerebral organoids. bioRxiv (2024). doi:10.1101/2024.02.20.580978

48. Zhang, Y. et al. Purification and Characterization of Progenitor and Mature Human Astrocytes Reveals Transcriptional and Functional Differences with Mouse. Neuron 89, 37–53 (2016).

49. Uenaka, T. et al. Prevention of transgene silencing during human pluripotent stem cell differentiation. Cell Stem Cell 33, 517–530.e8 (2026).

50. Foo, L. C. et al. Development of a method for the purification and culture of rodent astrocytes. Neuron 71, 799–811 (2011).

51. Shaner, N. C. et al. A bright monomeric green fluorescent protein derived from Branchiostoma lanceolatum. Nat. Methods 10, 407–409 (2013).

52. Dejanovic, B. et al. Changes in the Synaptic Proteome in Tauopathy and Rescue of Tau-Induced Synapse Loss by C1q Antibodies. Neuron 100, 1322–1336.e7 (2018).

53. Wu, T. et al. Complement C3 Is Activated in Human AD Brain and Is Required for Neurodegeneration in Mouse Models of Amyloidosis and Tauopathy. Cell Rep. 28, 2111–2123.e6 (2019).

54. Hong, S. et al. Complement and microglia mediate early synapse loss in Alzheimer mouse models. Science (80-.). 352, 712–716 (2016).

55. Wilton, D. K. et al. Microglia and complement mediate early corticostriatal synapse loss and cognitive dysfunction in Huntington’s disease. Nat. Med. 29, 2866–2884 (2023).

56. Lui, H. et al. Progranulin Deficiency Promotes Circuit-Specific Synaptic Pruning by Microglia via Complement Activation. Cell 165, 921–35 (2016).

57. Werneburg, S. et al. Targeted Complement Inhibition at Synapses Prevents Microglial Synaptic Engulfment and Synapse Loss in Demyelinating Disease. Immunity 52, 167–182.e7 (2020).

58. Howell, G. R. et al. Molecular clustering identifies complement and endothelin induction as early events in a mouse model of glaucoma. J. Clin. Invest. 121, 1429–1444 (2011).

59. Prakash, P. et al. Proteomic profiling of interferon-responsive reactive astrocytes in rodent and human. Glia 72, 625–642 (2024).

60. Gomes, C. et al. Induction of astrocyte reactivity promotes neurodegeneration in human pluripotent stem cell models. Stem Cell Reports 19, 1122–1136 (2024).

61. Walker, D. G., Kim, S. U. & McGeer, P. L. Expression of complement C4 and C9 genes by human astrocytes. Brain Res. 809, 31–38 (1998).

62. Dolan, M.-J. et al. Spatiotemporal Analysis of Remyelination Reveals a Concerted Interferon-Responsive Glial State That Coordinates Immune Infiltration. bioRxiv 2025.04.22.649486 (2025).

63. Voulgaris, D., Nikolakopoulou, P. & Herland, A. Generation of Human iPSC-Derived Astrocytes with a mature star-shaped phenotype for CNS modeling. Stem Cell Rev. Reports 18, 2494–2512 (2022).

64. Askenazi, M. et al. Compilation of reported protein changes in the brain in Alzheimer’s disease. Nat. Commun. 14, 4466 (2023).

65. Morshed, N. et al. Quantitative phosphoproteomics uncovers dysregulated kinase networks in Alzheimer’s disease. *Nat*. Aging 1, 550–565 (2021).

66. Johnson, E. C. B. et al. Large-scale deep multi-layer analysis of Alzheimer’s disease brain reveals strong proteomic disease-related changes not observed at the RNA level. Nat. Neurosci. 25, 213–225 (2022).

67. Huang, Z. et al. Brain proteomic analysis implicates actin filament processes and injury response in resilience to Alzheimer’s disease. Nat. Commun. 14, 2747 (2023).

68. Higginbotham, L. et al. Integrated proteomics reveals brain-based cerebrospinal fluid biomarkers in asymptomatic and symptomatic Alzheimer’s disease. Sci. Adv. 6, (2020).

69. Seifar, F. et al. Large-scale deep proteomic analysis in Alzheimer’s disease brain regions across race and ethnicity. Alzheimer’s Dement. 20, 8878–8897 (2024).

70. Allen, N. J. & Eroglu, C. Cell Biology of Astrocyte-Synapse Interactions. Neuron 96, 697–708 (2017).

71. Jen, Y. H. L., Musacchio, M. & Lander, A. D. Glypican-1 controls brain size through regulation of fibroblast growth factor signaling in early neurogenesis. Neural Dev. 4, 1–19 (2009).

72. Swain, S. et al. Low-Glucose Culture Conditions Bias Neuronal Energetics Towards Oxidative Phosphorylation. J. Neurochem. 169, (2025).

73. Grosjean, P. et al. Network-aware self-supervised learning enables high-content phenotypic screening for genetic modifiers of neuronal activity dynamics. *Nat*. Mach. Intell. 7, 2009–2025 (2025).

74. Fernandopulle, M. S. et al. Transcription Factor–Mediated Differentiation of Human iPSCs into Neurons. Curr. Protoc. Cell Biol. 79, 1–48 (2018).

75. Afshar-Saber, W. et al. An Open-Source Pipeline for Calcium Imaging and All-Optical Physiology in Human Stem Cell-Derived Neurons. Adv. Sci. 13, e15887 (2026).

76. Hyvärinen, T. et al. Functional characterization of human pluripotent stem cell-derived cortical networks differentiated on laminin-521 substrate: comparison to rat cortical cultures. Sci. Rep. 9, 17125 (2019).

77. Patel, M. D. et al. Human Astrocytes Synchronize Neural Organoid Networks. bioRxiv 1–23 (2024). doi:10.1101/2024.10.17.618921

78. Voufo, C. et al. Circuit mechanisms underlying embryonic retinal waves. Elife 12, 1–22 (2023).

79. Moreno-Juan, V. et al. Prenatal thalamic waves regulate cortical area size prior to sensory processing. Nat. Commun. 8, 14172 (2017).

80. Weissman, T. A., Riquelme, P. A., Ivic, L., Flint, A. C. & Kriegstein, A. R. Calcium Waves Propagate through Radial Glial Cells and Modulate Proliferation in the Developing Neocortex. Neuron 43, 647–661 (2004).

81. Pan, F., Mills, S. L. & Massey, S. C. Screening of gap junction antagonists on dye coupling in the rabbit retina. Vis. Neurosci. 24, 609–18 (2007).

82. Cruikshank, S. J. et al. Potent block of Cx36 and Cx50 gap junction channels by mefloquine. Proc. Natl. Acad. Sci. U. S. A. 101, 12364–12369 (2004).

83. Abudara, V. et al. The connexin43 mimetic peptide Gap19 inhibits hemichannels without altering gap junctional communication in astrocytes. Front. Cell. Neurosci. 8, 1–8 (2014).

84. Zhang, Y. et al. Rapid Single-Step Induction of Functional Neurons from Human Pluripotent Stem Cells. Neuron 78, 785–798 (2013).

85. Clayton, B. L. L. & Liddelow, S. A. Heterogeneity of Astrocyte Reactivity. Annu. Rev. Neurosci. 48, 231–249 (2025).

86. Hasel, P., Rose, I. V. L., Sadick, J. S., Kim, R. D. & Liddelow, S. A. Neuroinflammatory astrocyte subtypes in the mouse brain. Nat. Neurosci. 24, 1475–1487 (2021).

87. Choi, B. J., Chen, Y. & Desplan, C. Retinal calcium waves coordinate uniform tissue patterning of the Drosophila eye. Science (80-.). 390, eady5541 (2025).

88. Ackman, J. B., Burbridge, T. J. & Crair, M. C. Retinal waves coordinate patterned activity throughout the developing visual system. Nature 490, 219–25 (2012).

89. Peinado, A., Yuste, R. & Katz, L. C. Extensive dye coupling between rat neocortical neurons during the period of circuit formation. Neuron 10, 103–114 (1993).

90. Cina, C., Bechberger, J. F., Ozog, M. A. & Naus, C. C. G. Expression of connexins in embryonic mouse neocortical development. J. Comp. Neurol. 504, 298–313 (2007).

91. Maxeiner, S. et al. Spatiotemporal transcription of connexin45 during brain development results in neuronal expression in adult mice. Neuroscience 119, 689–700 (2003).

92. Miller, J. A. et al. Transcriptional landscape of the prenatal human brain. Nature 508, 199–206 (2014).

93. Doench, J. G. et al. Optimized sgRNA design to maximize activity and minimize off-target effects of CRISPR-Cas9. Nat. Biotechnol. 34, 184–191 (2016).

94. Horlbeck, M. A. et al. Compact and highly active next-generation libraries for CRISPR-mediated gene repression and activation. Elife 5, 1–20 (2016).

95. González, F. et al. An iCRISPR platform for rapid, multiplexable, and inducible genome editing in human pluripotent stem cells. Cell Stem Cell 15, 215–226 (2014).

96. Cerbini, T. et al. Transcription activator-like effector nuclease (TALEN)-mediated CLYBL targeting enables enhanced transgene expression and one-step generation of dual reporter human induced pluripotent stem cell (iPSC) and neural stem cell (NSC) lines. PLoS One 10, 1–18 (2015).

97. McKinney, W. Data Structures for Statistical Computing in Python. in Proceedings of the 9th Python in Science Conference 41, 56–61 (2010).

98. Van Der Walt, S., Colbert, S. C. & Varoquaux, G. The NumPy array: A structure for efficient numerical computation. Comput. Sci. Eng. 13, 22–30 (2011).

99. Virtanen, P. et al. SciPy 1.0: fundamental algorithms for scientific computing in Python. Nat. Methods 17, 261–272 (2020).

100. Waskom, M. Seaborn: Statistical Data Visualization. J. Open Source Softw. 6, 3021 (2021).

101. Hunter, J. D. Matplotlib: A 2D graphics environment. Comput. Sci. Eng. 9, 99–104 (2007).

102. Charlier, F. et al. Statannotations: v0.6. Zenodo (2022). doi:10.5281/zenodo.7213391

103. Schindelin, J., et al. Fiji: an open-source platform for biological-image analysis. Nat. Methods 9, 676–682 (2012).

104. Dobin, A. et al. STAR: Ultrafast universal RNA-seq aligner. Bioinformatics 29, 15–21 (2013).

105. Danecek, P. et al. Twelve years of SAMtools and BCFtools. Gigascience 10, 1–4 (2021).

106. Patro, R., Duggal, G., Love, M. I., Irizarry, R. A. & Kingsford, C. Salmon provides fast and bias-aware quantification of transcript expression. Nat. Methods 14, 417–419 (2017).

107. Kuehl, M., Wong, M. N., Wanner, N., Bonn, S. & Puelles, V. G. Gene count estimation with pytximport enables reproducible analysis of bulk RNA sequencing data in Python. Bioinformatics 40, 0–3 (2024).

108. Muzellec, B., Teleńczuk, M., Cabeli, V. & Andreux, M. PyDESeq2: a python package for bulk RNA-seq differential expression analysis. Bioinformatics 39, 2019–2021 (2023).

109. Pachitariu, M., Rariden, M. & Stringer, C. Cellpose-SAM: superhuman generalization for cellular segmentation. bioRxiv (2025). doi:10.1101/2025.04.28.651001

110. Thomson, J. A. et al. Embryonic stem cell lines derived from human blastocysts. Science 282, 1145–7 (1998).

111. Kreitzer, F. R. et al. A robust method to derive functional neural crest cells from human pluripotent stem cells. Am. J. Stem Cells 2, 119–31 (2013).

112. Pantazis, C. B. et al. A reference human induced pluripotent stem cell line for large-scale collaborative studies. Cell Stem Cell 29, 1685–1702.e22 (2022).

113. Quist, E., Ahlenius, H. & Canals, I. Transcription Factor Programming of Human Pluripotent Stem Cells to Functionally Mature Astrocytes for Monocultures and Cocultures with Neurons. Methods Mol. Biol. 2352, 133–148 (2021).

114. Soneson, C., Love, M. I. & Robinson, M. D. Differential analyses for RNA-seq: Transcript-level estimates improve gene-level inferences. F1000Research 4, 1–23 (2016).

115. Campbell, B. C. et al. mGreenLantern: a bright monomeric fluorescent protein with rapid expression and cell filling properties for neuronal imaging. Proc. Natl. Acad. Sci. U. S. A. 117, 30710–30721 (2020).

116. Shaner, N. C. et al. Improved monomeric red, orange and yellow fluorescent proteins derived from Discosoma sp. red fluorescent protein. Nat. Biotechnol. 22, 1567–1572 (2004).

117. Shcherbakova, D. M. & Verkhusha, V. V. Near-infrared fluorescent proteins for multicolor in vivo imaging. Nat. Methods 10, 751–754 (2013).

118. Zhang, Y. et al. Fast and sensitive GCaMP calcium indicators for imaging neural populations. Nature 615, 884–891 (2023).

119. Yokoyama, T. et al. A multicolor suite for deciphering population coding of calcium and cAMP in vivo. Nat. Methods 21, 897–907 (2024).

120. Stringer, C., Wang, T., Michaelos, M. & Pachitariu, M. Cellpose: a generalist algorithm for cellular segmentation. Nat. Methods 18, 100–106 (2021).

121. Savitzky, A. & Golay, M. J. E. Smoothing and Differentiation of Data by Simplified Least Squares Procedures. Anal. Chem. 36, 1627–1639 (1964).

